# A Self-Supervised Foundation Model for Robust and Generalizable Representation Learning in STED Microscopy

**DOI:** 10.1101/2025.06.06.656993

**Authors:** Anthony Bilodeau, Frédéric Beaupré, Julia Chabbert, Kamylle Thériault, Andréanne Deschênes, Jean-Michel Bellavance, Koraly Lessard, Renaud Bernatchez, Paul De Koninck, Christian Gagné, Flavie Lavoie-Cardinal

## Abstract

Foundation Models (FMs) have dramatically increased the potential and power of deep learning algorithms through general capacities over a variety of tasks. The performance increase they offer is obtained without elaborated specific trainings for domains such as natural language processing and computer vision. However, their application in specialized fields like biomedical imaging and fluorescence microscopy remains difficult due to distribution shifts and the scarcity of high-quality annotated datasets. The high cost of data acquisition and the requirement for in-domain expertise further exacerbate this challenge in microscopy. To address this we introduce STED-FM, a foundation model specifically designed for super-resolution STimulated Emission Depletion (STED) microscopy. STED-FM leverages a Vision Transformer architecture trained at scale with Masked Autoencoding on a new dataset of nearly one million STED images. STED-FM learns expressive latent representations without requiring extensive annotations, yielding robust performance across diverse downstream microscopy image analysis tasks. Unsupervised experiments demonstrate the discriminative structure of its learned latent space. These representations can be leveraged for multiple downstream applications, including fully supervised classification and segmentation with reduced annotation requirements. Moreover, STED-FM representations enhance the performance of deep learning–based image denoising and improve the quality of images generated by diffusion models, enabling latent attribute manipulation for the data-driven discovery of subtle nanostructures and phenotypes, as well as algorithmic super-resolution. Moreover, its powerful structure retrieval capabilities are integrated into automated STED microscopy acquisition pipelines, paving the way for smart microscopy. In sum, we demonstrate that STED-FM lays a robust foundation for state-of-the-art algorithms across a wide array of tasks, establishing it as a highly valuable and scalable resource for researchers in super-resolution microscopy.

## 1 Introduction

Deep learning has become a powerful tool for predictive and quantitative analysis, driven by its ability to learn expressive representations of complex, high-dimensional data. The scaling hypothesis suggests that performance improves with larger models, datasets, and computational resources [1], which gave rise to the development of Foundation Models (FMs) [2]: large-scale pre-trained models capable of excelling at various tasks. This is in contrast with most classical machine learning approaches, where models are trained to accomplish one specific task (e.g., classification, regression), based on a dataset associated to the domain.

They have been extensively studied for natural language processing [3, 4] and computer vision [5]. FMs have benefited from large curated datasets for training [6–9] and show impressive generalization capability on downstream tasks [10]. However, in specialized fields such as biomedical imaging, their effectiveness remains uncertain due to significant distribution shifts between downstream and upstream domains [11]. In biomedical imaging, limited data availability poses a challenge, as both fine-tuning of models pre-trained on natural images and training a new model from scratch often result in suboptimal outcomes [12]. Consequently, there is a growing interest in developing FMs tailored specifically to biomedical and bioimaging applications [13–16].

One of the most significant challenges associated with the development of FMs for bioimaging tasks is the scarcity of high-quality annotated datasets [17]. This can be explained by the cost of data acquisition as well as the in-domain expertise and time required to annotate [18–20]. Moreover, variability in the imaging modalities and experimental settings, including batch effects, will inevitably increase the complexity of dataset aggregation and curation [21]. It will also lead to heterogeneous data distributions that need to be learned by the FMs. This can be challenging in a context where available datasets are of limited size [21]. These challenges can hinder the training of expressive FMs for biomedical imaging.

Yet, wide-ranging successes in other domains have motivated the design and training of FMs adapted to the bioimaging research field. Efforts have been put into the collection and curation of large bioimaging datasets [13, 22–25] and of datasets in more specialized fields, such as chemistry [26] and microscopy [15, 27–31]. Leveraging such datasets, researchers have proposed to use the FM paradigm to either train deep learning models in a fully-supervised fashion [13, 25, 30], using a combination of fully-supervised and self-supervised learning (SSL) [12], or with self-supervised learning only [18, 19]. SSL refers to the training procedures that aim to learn general representations from a dataset without any annotations [32–36]. On natural images, it has been shown that SSL can achieve similar or superior performance to their fully-supervised counterparts, in particular when few annotations are available [18]. One family of SSL techniques uses the strategy of corrupting inputs and training networks to reconstruct the corruption-free versions [35]. For example, masked autoencoders [37] (MAEs) learn to reconstruct masked patches from a transformer encoder using only the visible encoded patches and a lightweight decoder. The encoder can afterwards be used as a general-purpose model. Such methods have been shown to scale well and to generalize efficiently to unseen datasets and tasks [37], including in biomedical imaging [31, 38, 39].

Due to their ability to learn meaningful representations from unannotated image data, SSL methods have already been successfully applied in microscopy for tasks such as image denoising and resolution enhancement [15, 40–45], object detection [46], and segmentation [47]. Recent work pointed out that in-domain pre-training can significantly impact the performance on downstream tasks, especially in the small data regime [16, 25]. In this work, we develop a FM tailored for super-resolution STimulated Emission Depletion (STED) microscopy, hence called STED-FM. Super-resolution microscopy, or optical nanoscopy, has transformed our ability to investigate biology at the molecular scale [48]. However, its complexity and extreme dependence on optimal parameters for acquisition and sample preparation [49] further limit data availability. Meanwhile, the richness of hidden information that could be extracted remains largely under-exploited [50]. Our model was specifically designed to perform a wide array of downstream tasks on small STED imaging datasets. To this end, we created a dataset of nearly one million STED images for the training of STED-FM [51], a general-purpose vision transformer (ViT) [52] (Fig. 1a). Our approach leverages the task-agnostic, auto-regressive MAE framework, originally introduced for natural images to emulate self-supervised training strategies used in natural language processing [37] (Fig. 1b). MAE is a well established method for downstream tasks such as classification and segmentation in biomedical [39, 53] and specialized fields, such as microscopy [31, 38].

**Figure 1:**
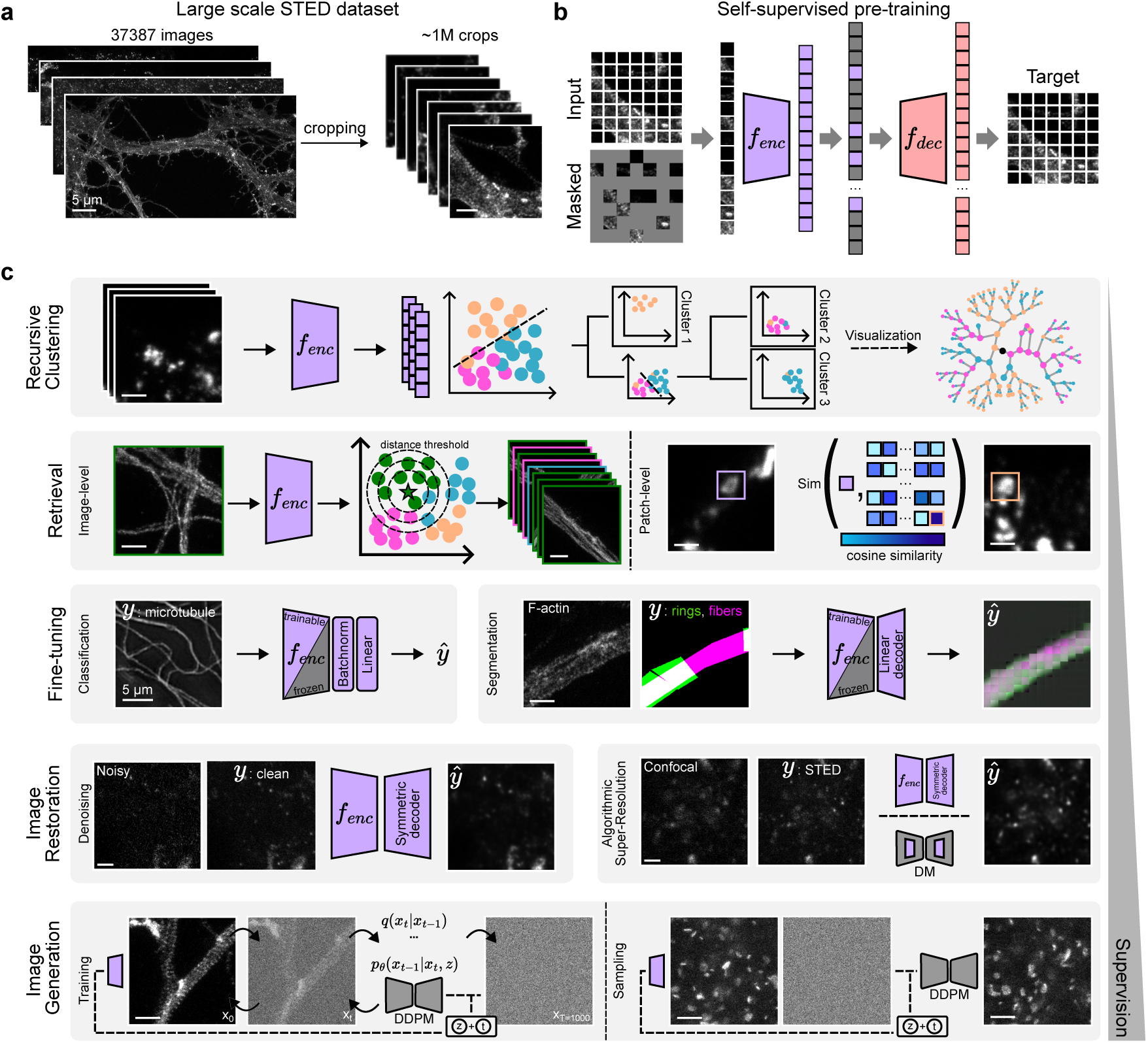
Overview of the STED-FM deep learning framework. **a)** Large dataset of STED images cropped into smaller regions of 224 *×* 224 pixels resulting in a dataset of nearly one million crops used to train STED-FM. **b)** MAE training procedure for pre-training ViTs. **c)** Schematized overview of the different tasks performed by the pre-trained STED-FM model, listed in increasing order of supervision required. Scale bars: 1 µm unless otherwise specified.

We conduct a comprehensive evaluation of STED-FM’s learned latent representations and demonstrate their utility across various tasks. The expressiveness of the learned manifold is validated through clustering (Fig. 1c: *Recursive Clustering* ) and instance retrieval, where similar structures are grouped coherently (Fig. 1c: *Retrieval* ). On standard downstream tasks such as classification and segmentation, STED-FM reduces the amount of required annotated data for training (Fig. 1c: *Fine-tuning* ). STED-FM is also validated on low-level tasks such as image denoising and algorithmic super-resolution. Furthermore, conditioning diffusion-based generative models on latent codes generated by STED-FM improves the quality and visual fidelity of generated images (Fig. 1c: *Image Generation*), enabling attribute manipulation and pattern discovery. Finally, STED-FM’s learned representations can support structural feature detection and image quality assessment as part of automated image acquisition workflows.

## 2 Results

We created a large curated dataset of approximately 1 million STED images (224 × 224 pixels crops, extracted from 37,387 large images of variable sizes) to enable the training of STED-FM (*STED-FM dataset* , Methods, Fig. 1a) [51]. The dataset consisted of images of more than 20 different fluorescently-labeled proteins in different cell types (hippocampal and cortical neurons, HEK, NG108, Caco2), imaging conditions (live and fixed cells), and sample type (cell culture, tissue slices) acquired on an Abberior STED Expert Line microscope (Methods). We initially considered four ViT sizes for STED-FM: ViT-tiny, -small, -base, and -large. Since the ViT-Small (ViT-S) and ViT-base models showed similar performance in early downstream classification experiments (Supplementary Fig. 1), we selected the ViT-S architecture to minimize compute resources. STED-FM refers to the ViT-S weights obtained after pre-training on the *STED-FM dataset* .

For baseline comparison, we used the publicly available pre-trained weights from *ImageNet* [6] and also pre-trained the ViT-S architecture on three broadly used microscopy datasets: the Joint Undertaking in Morphological Profiling Cell Painting dataset (*JUMP*, 3.7M images) [27], the Human Protein Atlas dataset (*HPA*, 1.1M images) [54], and a Structured Illumination Microscopy (*SIM*, 205k images) dataset [15]. The pre-training datasets cover a wide range of domains and dataset sizes: natural images, microscopy, and super-resolution microscopy (Supplementary Table 1).

### 2.1 Unsupervised Analysis

As an initial evaluation, we aimed to assess the quality and robustness of the representations learned by STED-FM during pre-training, and to explore the breadth of capabilities enabled by the latent space. We validated the learned representations of STED-FM against that of the ViT-S pre-trained on *ImageNet*, *JUMP*, *HPA*, and *SIM*.

The attention mechanism of ViTs provides a direct probe into how the model assigns importance to image regions, which should intuitively align with an expert’s notion of biological relevance. To quantify this alignment, we sampled 40 F-actin images, extracted the attention maps from the models pre-trained on different datasets (Methods), and asked experts to select their preferred map (Fig. 2a,b). STED-FM’s attention maps were selected over 60% of the time. This suggests that STED-FM’s attention aligns better with human preference, making its representations and subsequent predictions more interpretable.

**Figure 2:**
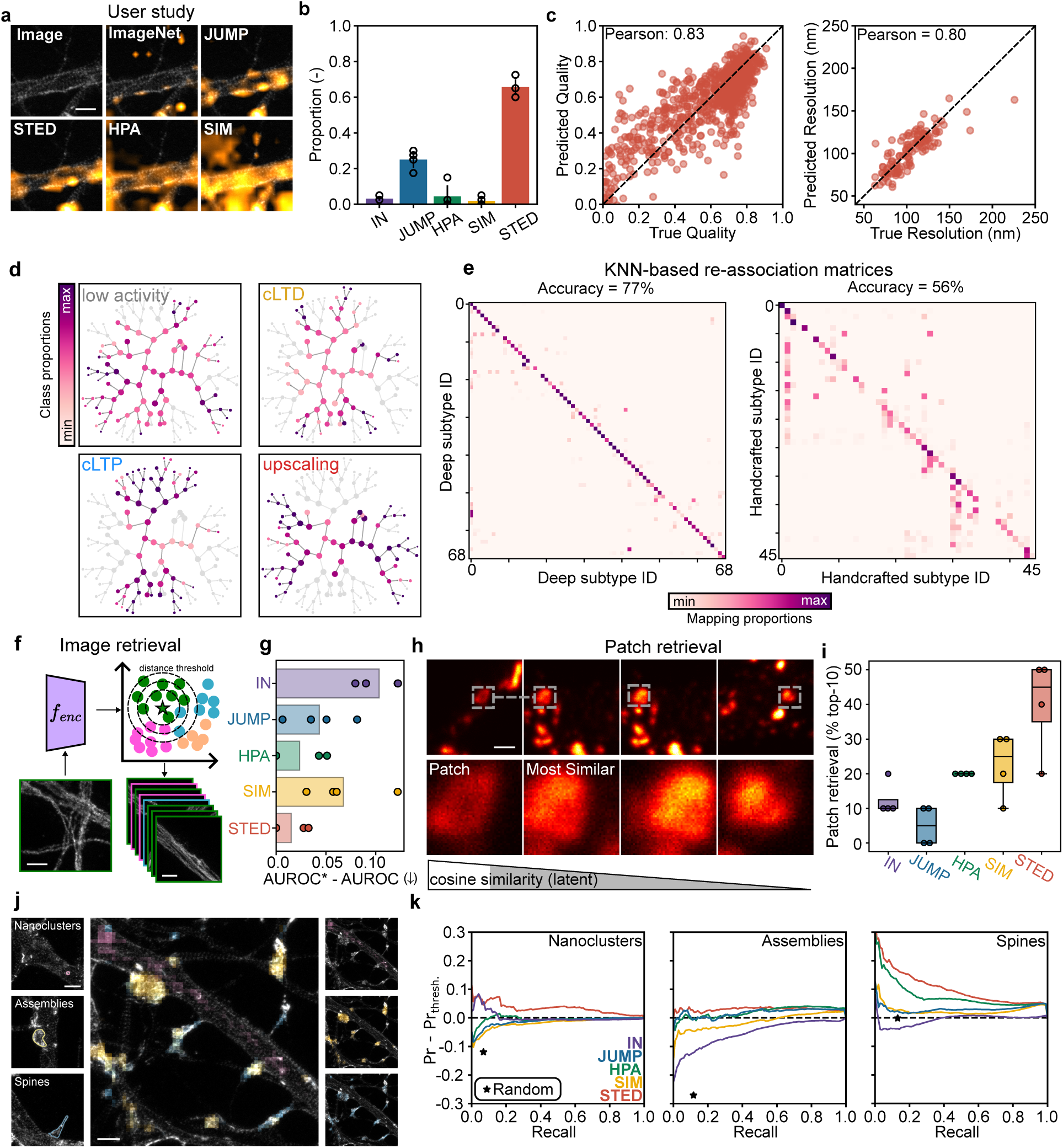
Unsupervised analysis of the latent space learned during pre-training. **a)** Attention maps generated from the learned representation of STED-FM and ViT-S pre-trained on *ImageNet*, *JUMP*, *HPA* and *SIM* datasets used in the user study. **b)** Alignment of the different models’ attention maps with expert preference. **c)** Correlation between ground truth and inferred image attribute values obtained from a ridge regression model trained on STED-FM’s image embeddings. **d)** Graph visualizations of the recursive consensus clustering (RCC) algorithm performed on the *NAS Dataset*. The different neuronal stimulation conditions are associated with distinct patterns in the RCC graph, supporting the emergence of activity-dependent synaptic subtypes. Node sizes are given by the number of embeddings in the node. Nodes are color-coded by the proportion of the given class. **e)** Matrices obtained from re-assigning every image embedding to the majority class of its 10 nearest neighbors, where class refers to leaf node identifier. RCC performed on the latent manifold is more coherent than when performed on the hand-crafted manifold. **f)** Schematic of the image retrieval task. **g)** Image-based retrieval of distinct biological structures in 4 downstream datasets (*SO, NAS, Px, PR*). For each dataset and pre-trained model, we report the difference to the best performing model (AUROC*) for structure-specific retrieval. On average, STED-FM shows near-optimal performance, with AUROC*-AUROC close to 0. **h)** Schematic and retrieved examples from the patch retrieval task using STED-FM templates for the identification of perforated synapses in the *Synaptic Protein Zoo Dataset*. **i)** Patch retrieval performance of the different pre-trained models for the identification of perforated synapses averaged over 4 experts’ preferences. **j)** Retrieval of different F-actin conformations as templates: nanoclusters, compact assemblies, and spines using STED-FM on the *F-Actin Conformation Dataset*. **k)** Segmentation precision over the range of recall for each pre-trained model compared to using an intensity threshold-based approach on the pixel intensity distribution. Manual annotations were obtained for each conformation on 79 images. *Random* corresponds to re-assigning the manual annotations randomly on the foreground. Scale bars: 1 µm.

Next, we used ridge regression to inspect whether physical attributes, such as image quality and spatial resolution, were encoded in the latent space (Methods, Fig. 2c). For both attributes, a Pearson correlation coefficient of at least 0.80 is obtained by STED-FM, indicating that it encodes relevant image features in its latent space. Interestingly, other pre-trainings lead to decreased correlation, suggesting that these attributes are not as efficiently learned from these datasets (Supplementary Fig. 2).

We investigated the global structure of the latent manifold using recursive consensus clustering (RCC) applied to STED-FM’s embeddings from the *Neural Activity States Dataset (NAS)*, which contains images of the post-synaptic protein PSD95 in fixed hippocampal neurons treated with different activity-modulating solutions (Methods) [55]. The RCC procedure, adapted from Sonpatki & Shah [56], involves recursively applying consensus clustering [57] to STED-FM’s embeddings until leaf nodes are formed (Methods). This process, also applied to manually extracted features (Supplementary Fig. 3), allows visualization of the clustering tree (Fig. 2d) to evaluate if STED-FM extracts discriminative structural features associated with different neuronal activity states (Supplementary Fig. 4a & 5 & Supplementary Table **??**). To quantify the manifold’s quality we performed *K*-Nearest Neighbors (KNN) classification (*K* = 9, Methods) of the data onto associated leaf nodes. STED-FM embeddings achieved an accuracy of 77% (Fig. 2e, left), outperforming the 56% accuracy obtained with hand-crafted features (Fig. 2e, right). This demonstrates that the latent manifold learned by STED-FM offers superior coherence, representational power, reduction of redundancy, and discriminative capability compared to manifolds built from manually extracted features (Supplementary Fig.3 and 4).

To address the time-consuming visual inspection required by experts in microscopy image analysis to identify specific structural phenotypes, we leveraged the latent representations of STED-FM for automated image and patch retrieval of similar sub-structures in large STED images [58–60]. We evaluated performance on datasets acquired on the same microscope as pre-training data (In Domain STED (ID-STED); *STED Optimization Dataset - SO* [61], *NAS* [55]) and on different STED microscopes (Out Of Domain STED (OOD-STED); *Peroxisome Dataset - Px* [62], *Polymer Rings Dataset - PR* [63]). For image retrieval, we used the cosine distance to measure the similarity between a template image embedding and all other embeddings (Fig. 2f & Supplementary Figure 6, Methods). STED-FM consistently achieved the best or near-optimal performance on all downstream dataset (Fig. 2g).

For patch retrieval, we applied STED-FM to automate the challenging task of retrieving rare structures, focusing on perforated synaptic structures from the annotated *Synaptic Protein Zoo Dataset - SPZ* (ID-STED) [64] (Fig. 2h). A user study demonstrated that STED-FM is better at retrieving patches matching the perforated synapse template compared to the other ViT-S pre-trainings (Fig. 2i, Methods). This highlights STED-FM’s capacity to encode subtle nano-organization patterns required to identify these rarely observable structures in the *SPZ* (*↑* 3%).

Finally, we leveraged the patch-retrieval capability of the pre-trained models to automatically highlight regions of interest on the *F-actin Conformation Dataset* (ID-STED) [65, 66] (Fig. 2j), automating a task that traditionally requires time-consuming manual annotation [65, 67]. Using a single template per class, we retrieved patches corresponding to: 1) F-actin nanoclusters, 2) F-actin compact assemblies, and 3) dendritic spines (Fig. 2j). Compared to an intensity threshold-based approach and other ViT-S pre-trainings, STED-FM outperforms other methods for the segmentation of all structures, according to manual ground truth annotations (Fig. 2k).

### 2.2 Supervised post-training

We evaluated the quality of the learned representations by using them as initialization for supervised post-training on two downstream tasks: 1) classification and 2) segmentation. For both tasks, we added a linear layer to the STED-FM backbone to predict the final output (Methods). We considered two training strategies: linear probing (freezing the backbone and training only the linear layer) and end-to-end fine-tuning (training the entire network). To simulate common bioimaging limitations, we also assessed performance using limited fine-tuning data. To evaluate the influence of the pre-training dataset size on FM performance, we compared the classification accuracy of JUMP-FM, HPA-FM, and STED-FM trained on 976 022 images (Supplementary Fig. 7; Methods). Downsampling the JUMP dataset from 3.7 million to 976 022 images resulted in a substantial reduction in accuracy, whereas no significant effect was observed for the HPA dataset. Consequently, subsequent analyses were conducted using FMs trained on their respective full-size datasets.

For downstream classification, we used two ID-STED (*SO, NAS* ) and two OOD-STED (*Px, PR*) datasets (Extended Data Fig. 1a, Supplementary Table 3). After linear probing, STED-FM performed best overall (Extended Data Fig. 1b), with the largest performance gaps observed on ID-STED datasets in the small data regime (*SO, NAS*, Supplementary Fig. 8). All models using the STED-FM backbone exceed the performance of models trained from scratch. This demonstrates the advantage of using similar-domain, large-scale pre-trained networks over training from scratch on limited super-resolution microscopy data. Results for end-to-end fine-tuning (Extended Data Fig. 1c) were similar, with STED-FM remaining the best overall, though the performance gap narrowed. This is likely due to the larger number of trainable parameters compensating for less effective pre-trainings.

For downstream segmentation, we used two ID-STED (*F-actin* [65], *SPZ* [64]) and two OOD-STED (*Foot processes (FP)* [68], *Lioness* [29]) datasets (Extended Data Fig. 1d, Supplementary Table 4, and Supplementary Fig. 9). Results from both linear probing (Extended Data Fig. 1e) and end-to-end fine-tuning (Extended Data Fig. 1f) confirmed that STED-FM achieved the best overall performance across all datasets and data regimes (Supplementary Fig. 10, Supplementary Tables 5-6). Notably, *ImageNet* initialization dropped below random initialization performance, highlighting the importance of in-domain pre-training for dense prediction tasks. We observe a strong inverse correlation (Pearson = -0.92) between segmentation performance and the perimetric complexity [69] of the target structures (Methods, Supplementary Fig. 11), confirming that morphologically complex structures are inherently more challenging to segment. Finally, we show that adding a symmetric decoder can improve segmentation performance, particularly for more complex tasks (Methods, *FP* in Supplementary Fig. 12).

We investigated the generalization properties of STED-FM on four out-of-distribution microscopy (OOD-MIC) datasets for classification (*BBBC026*, *BBBC052*, *BBBC053*, and *HPA-Classification*) and two OOD-MIC dataset for segmentation (*LCN*, and *DeepD3* ). STED-FM performs robustly even on these OOD-MIC tasks (Supplementary Fig. 13-15). To quantify domain distance independent of shift-sensitive pixel metrics, or dataset-dependent deep features [70–74], we opted for a power spectrum density analysis [75], which uses differences in frequency content (Extended Data Fig. 2a,b). Intuitively, closer domains should yield better performance, yet STED-FM achieves better-than-average performance across all tasks even when its pre-training domain is not the closest (top-right region, Extended Data Fig. 2c). We reasoned this superior performance stems from factors beyond domain closeness. Investigating pre-training dataset diversity, measured by the average intra-distance between images, we found a Pearson correlation of 0.60 between dataset diversity and downstream performance on classification and segmentation tasks, with STED-FM having the highest diversity (Extended Data Fig. 2d), suggesting that a more diverse pre-training dataset likely leads to better learned representations. Given the improved latent representation of STED-FM in both unsupervised and supervised downstream analysis in comparison to the other pre-trained FM, we only consider STED-FM for further downstream tasks.

### 2.3 Image Restoration

In modern microscopy, image restoration has become an important component of the imaging pipeline, allowing for the enhancement of signal quality while minimizing the necessary light dose [76]. Specifically, we tackled two image restoration tasks: denoising (improving signal-to-noise ratio (SNR)) and algorithmic super-resolution (generating structures unseen in low-resolution input).

We first evaluated using STED-FM’s latent representations for denoising by adding a symmetric decoder to the frozen backbone (Methods). We compared this model against four baselines: CARE2D [76], Noise2Void (N2V) [77], pix2pix [78], and the STED-specific baseline UNet-RCAN [79] (Methods). The baselines were trained and benchmarked across four OOD-STED datasets (Supplementary Table 7, Methods). In simpler denoising tasks, where the difference in SNR between noisy and high SNR is small (Supplementary Fig. 16), STED-FM performed similarly to CARE2D and UNet-RCAN (Supplementary Fig. 17), but in more challenging tasks, when the foreground signal is barely above the background noise level, STED-FM outperforms other baselines (Supplementary Fig.18-20).

Next, we addressed algorithmic super-resolution on a dataset of paired confocal and STED images of the synaptic protein VGAT (Supplementary Table 8, OOD-STED). We compared the ability of STED-FM and the three baselines CARE2D [76], pix2pix [78], and UNet-RCAN [79] to reconstruct a super-resolved STED image from a reference confocal (Methods). STED-FM outperforms all baselines across all computed pixel-wise metrics (Supplementary Fig. 21). For a task requiring the generation of complex structures unresolved in the confocal images, we trained two generative models (pix2pix and a diffusion model (DM)) on the F-actin dendritic dataset [80]. To evaluate their performance, we introduced an additional metric assessing a pre-trained U-Net’s ability to segment F-actin rings and fibers in the generated images relative to expert’s ground truth annotations (Methods). The DM significantly outperforms pix2pix on this segmentation task and all pixel-wise metrics (MSE, PSNR, and SSIM) (Methods, Fig. 3d-e & Supplementary Fig. 22). We then evaluated if we could enhance the DM generations by leveraging STED-FM’s expressive representations. We used the DRaFT framework [81] to fine-tune the DM using STED-FM’s latent representation similarity between synthetic and generated images as a reward (Methods, Fig. 3c). Although pixel-wise metrics remain unchanged (Fig. 3d,e & Supplementary Fig. 22), a user-study (N=4) confirmed that experts perceived the fine-tuned DM images as better aligned with real STED images, demonstrating superior subjective quality (Methods, Fig. 3f). Importantly, STED-FM is not required after fine-tuning.

**Figure 3:**
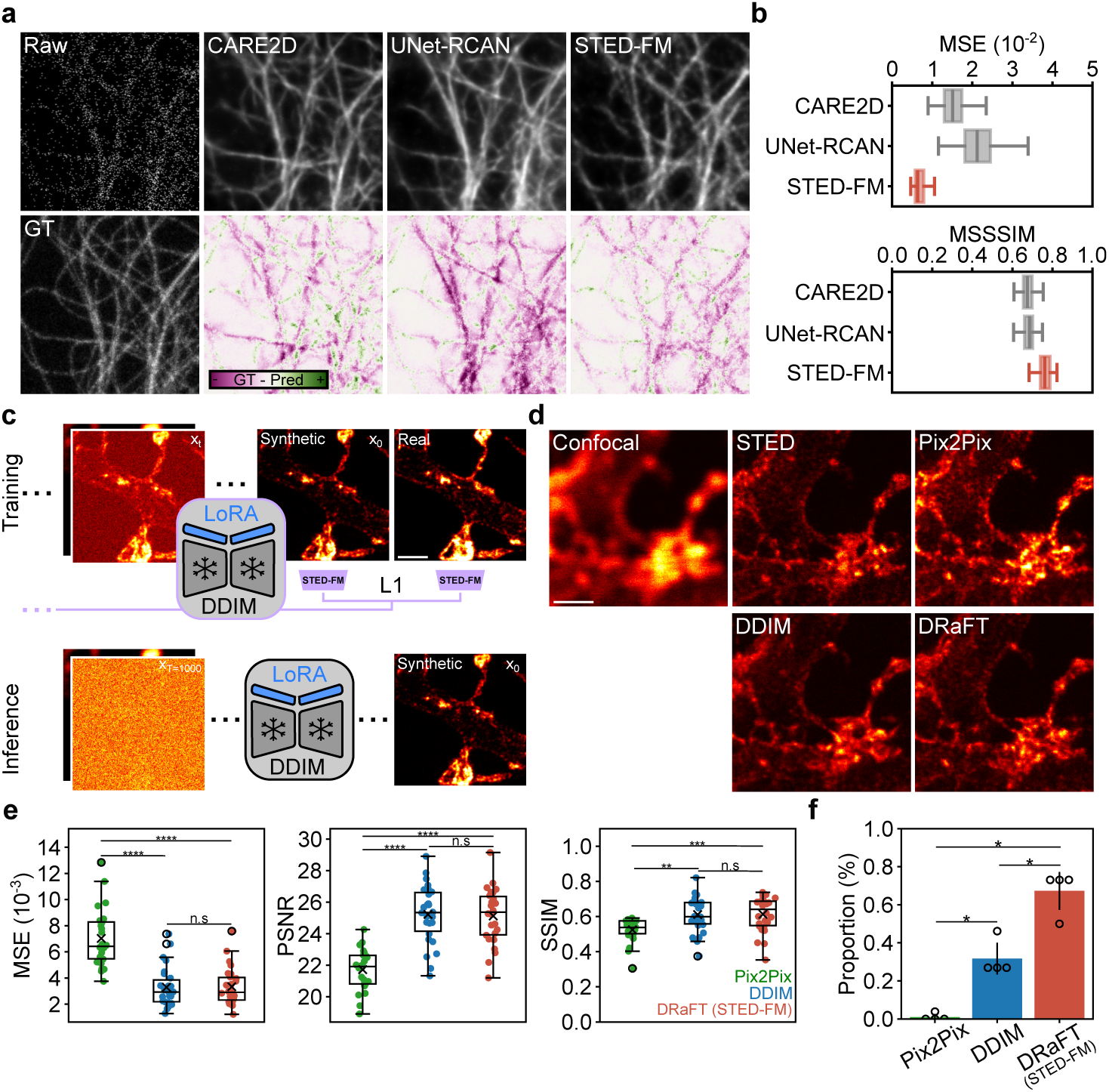
Evaluation of STED-FM on image restoration tasks. **a)** Image denoising task evaluation on the *ω*-tubulin (OOD-STED) dataset. (Top, left) Raw, noisy input image and the corresponding ground truth image (Bottom, left). (Top) Denoised outputs generated by the baselines and the STED-FM model. (Bottom) Residual image, calculated by subtracting the denoised output from the ground truth and colorcoded for regions where the denoised signal is too high (magenta) or too low (green). **b)** Evaluation of the denoising performance using pixel-wise metrics: MSE (mean squared error) and MSSSIM (multiscale structural similarity). Higher MSSSIM and lower MSE indicates better performance. Boxplots present the the sample median, the first and third quantiles. The whiskers are 1.5 times the interquartile range. Statistical analysis is provided in Supplementary Fig. 18. (N=100) **c)** Algorithmic super-resolution using the DRaFT framework. Low-Rank Adaptation (LoRA) layers are added to the frozen diffusion model. During training, an image is sequentially denoised using the DDIM sampling procedure. STED-FM’s latent representations of the synthetic image (*x*_0_) and the real STED image are compared using a mean squared error loss. This loss is backpropagated through the sampling procedure to update the LoRA layers (Methods). At inference, STED-FM is not required. **d)** Example images of generated F-Actin nanostructures with pix2pix, DDIM and DRaFT models from a reference confocal image and compared with the ground truth STED image. **e)** Evaluation of the algorithmic super-resolution performance using standard pixel-wise metrics: MSE, PSNR (peak signal-to-noise ratio), and SSIM (structural similarity). Higher SSIM and PSNR as well as lower MSE indicate better performance. **f)** Results of a user study indicating the proportion of instances in which human participants (N=4) selected each model’s reconstruction as more similar to the ground-truth STED image (N=26). The DRaFT fine-tuning with STED-FM method was selected more frequently than other baselines. Boxplots present the the sample median, the first and third quantiles, and ’x’ marks the mean. The whiskers are 1.5 times the interquartile range. The p-values for all metrics and the user study are given in Supplementary Tables 9, 10, 11.

These results demonstrate that STED-FM’s latent representations provide a powerful foundation for microscopy image restoration, enabling both discriminative and generative models to be used in OOD-STED contexts and outperform task-specific baselines. They demonstrate the utility of STED-FM for enhanced signal recovery and show that generation of faithful and quality nanostructures can be significantly improved through STED-FM conditioning of generative models.

### 2.4 Image Generation

#### 2.4.1 Diffusion models conditioned on pre-trained latent codes

We explored using STED-FM’s latent representations as conditioning signal for a denoising diffusion probabilistic model (DDPM [82]). This *latent guidance* framework allows the decoding of latent representations back to pixel space, serving as a powerful tool to analyze and interpret STED-FM’s learned manifold. We argue that using latent guidance to train the DDPM is superior to the more commonly used classifier-free guidance framework [83]. It improves the conservation of biological structures in the generated images and provides an interpretable tool to investigate the representation space of STED-FM.

Our approach (Fig 4a) closely follows Preechakul *et al.* [84], adapting the standard DDPM by adding a conditioning signal **z** corresponding to STED-FM’s embedding of the input image (Methods). The intuition is that conditioning on this rich latent code should improve the quality and visual fidelity of the generated images (Fig. 4a).

**Figure 4:**
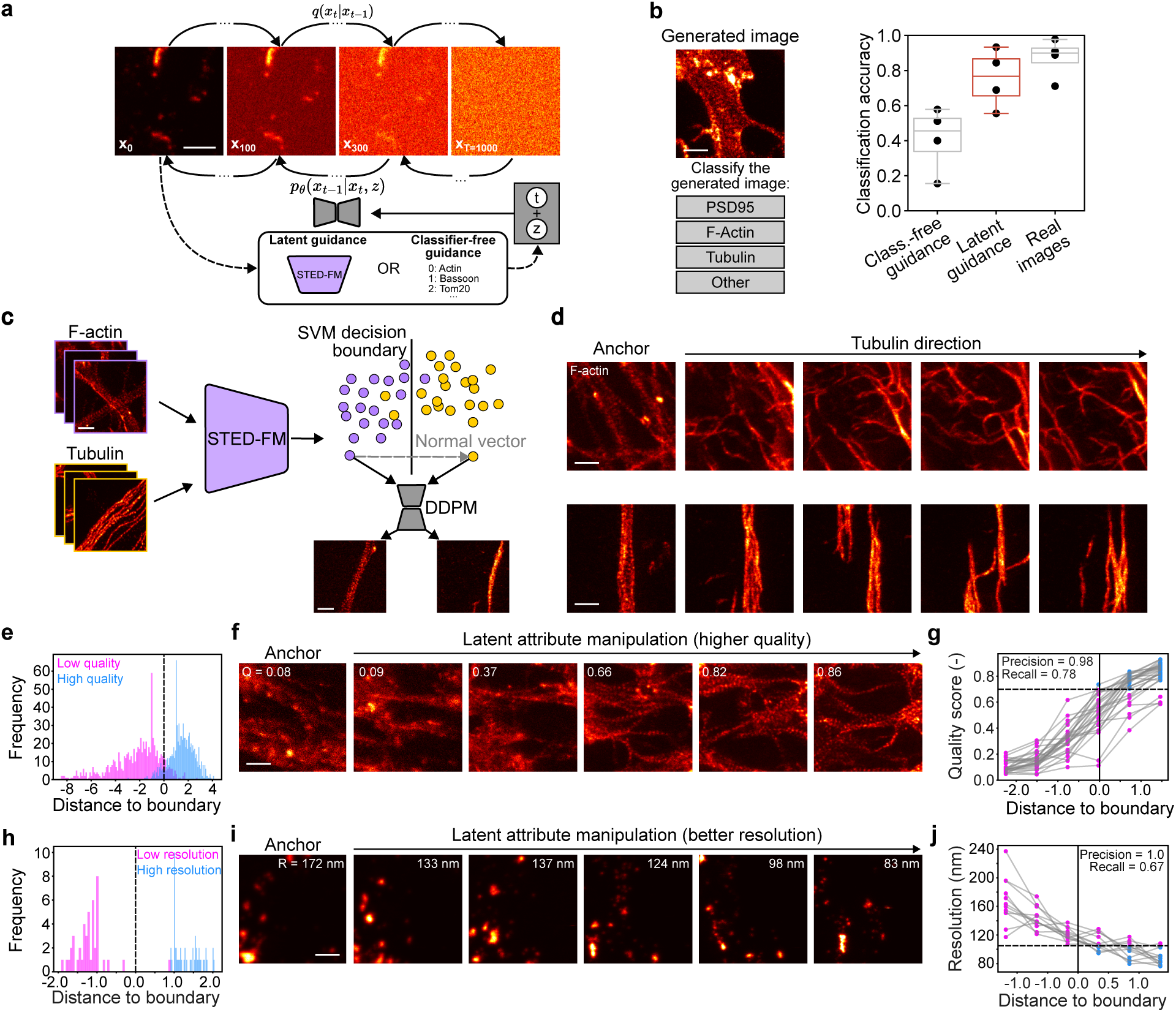
Image generation and latent attribute manipulation. **a)** Overview of the latent guidance and classifier-free guidance frameworks for training DDPMs. (*q*: diffusion kernel, *p*: learned denoising kernel, **t**: timestep embedding, **z**: conditioning vector) **b)** A user study was designed to evaluate experts’ classification accuracy on real and generated images (N=4 experts). **c)** Schematic of the latent attribute manipulation pipeline. A SVM is trained on the latent representations from STED-FM to discriminate between two groups. The latent code of an image is manipulated by interpolation in the direction perpendicular to the decision boundary (normal vector). **d)** The latent code manipulation pipeline allows translation of structures in images of F-actin into features associated with tubulin. **e-j)** Latent code manipulation of spatial features encoded in super-resolution microscopy images, such as image quality (**e-g**) and spatial resolution (**h-j**) **e, h)** Histogram of embeddings’ distances to the SVM’s decision boundary. **f, i)** Image generation trajectories using latent code manipulation. **g, j)** The quality score and spatial resolution correlate with the distance to the boundary, enabling precision and recall measurements (Methods). Every data point is color-coded according to a threshold (high quality: *>* 0.7; high resolution: *<* 105*nm*; dashed horizontal line). By splitting the graph into four quadrants using this quality (g) or resolution (j) threshold and the SVM’s decision boundary (solid vertical line), we define false positive, true positive, false negative and true negative regions (quality: read clockwise from the top-left region; spatial resolution: read clockwise from the bottom-left region). An ideal system, perfectly classifying points to the right of the boundary as high quality (high resolution) and those to the left as low quality (low resolution), would achieve recall and precision scores of 1.0. Scale bars: 1 µm.

To assess our latent-guided DDPM’s performance, we conducted a user study in which experts (N=4) identified the primary structure (F-actin, Tubulin, PSD95, or Other for unidentifiable patterns) in 45 images sampled from the *SO* dataset (Fig. 4b, left). We compared expert classification accuracy on real and synthetic images generated via latent or classifier-free guidance (Fig. 4b, right). Accuracy for the latent guidance synthetic images is similar to that of real images and superior to classifier-free guidance (Supplementary Figure 23). This supports the use of latent guidance to generate high-quality, high-fidelity images with features that are coherent and identifiable, matching those observed in real data.

#### 2.4.2 Latent code manipulation reveals encoded attributes

A functionality inherent to the latent guidance procedure is the ability to manipulate the conditioning vector in the latent space of STED-FM. We can use the DDPM to decode the manipulated vectors and visually inspect the associated generated images, which helps further assess the encoding quality and representation capacity of STED-FM. We adapted the methodology outlined by Shen *et al.* [85] and investigate whether STED-FM can encode nanoscopy features and semantics. The process, schematized in Fig. 4c, begins by classifying image embeddings into two classes corresponding to a chosen attribute. A support vector machine (SVM) is then trained on these classified embeddings. The normal vector to the SVM’s decision boundary provides the attribute direction in the latent space (Methods). The main idea is to manipulate the latent codes of unseen images along this direction. The resulting manipulated code is subsequently used as the conditioning vector for the latent-guided DDPM. This allows the DDPM to generate a version of the input image that exhibits features associated with the opposite class.

We first demonstrate STED-FM’s ability to encode specific biological structures in an experiment that aimed to interpolate images of one protein (F-actin) into another (tubulin) (Fig. 4d). The latent *F-actin to tubulin* direction helps generate images that progressively lose the typical F-actin periodic structures and longitudinal fibers [65]. Instead, these transform into continuous filaments, typical of STED tubulin images. This result highlights the ability of STED-FM to encode and discriminate between these two complex image attributes.

Next, we aimed to probe and manipulate spatial features in super-resolution microscopy images : image quality [61] and spatial resolution. For both attributes, the SVM’s decision boundary defined the direction of manipulation in the latent space (Fig. 4e,f & h,i). We demonstrate attribute manipulation capability by generating latent trajectories that successfully transformed input images towards the opposite attribute class (*e.g.*, low-quality to high-quality (N=27); Fig. 4f & Supplementary Fig. 24 or low-resolution to high-resolution (N=12); Fig 4i & Supplementary Fig. 25). Quantifying these trajectories revealed that the distance to the SVM boundary correlates with the inferred attribute scores (Fig. 4g,j). Specifically, the quality manipulation achieves a high precision and recall of 0.98 and 0.78, respectively, and the resolution manipulation achieved a precision and recall of 1.00 and 0.67, respectively (Methods). These results confirm that STED-FM effectively encodes relevant, separable imaging attributes within its latent representation, which can be manipulated by a latent-guided DDPM to enhance interpretability.

### 2.5 Pattern discovery

We now use the representational encoding of STED-FM with the decoding ability of the latent-guided DDPM to *discover* how biological phenotypes evolve as we navigate in the latent space. We first explored the age-dependent remodeling of synaptic structures in a dataset consisting of STED images of the synaptic protein PSD95 in cortical neuronal cultures that were fixed at 12 and 25 days in vitro (DIV12 and DIV25; *Synaptic development dataset* [86]). We sampled independent images from DIV12 neurons (*N* = 62), and manipulated their latent representations towards DIV25 neuron embeddings (Methods, Fig. 5a, Supplementary Fig. 26). We then analyzed how 7 structural features used to describe synaptic nanoclusters in super-resolution microscopy images [55] (Methods) changed along the manipulation trajectories (Fig. 5b, Supplementary Fig. 27). When moving in the latent space from representations associated with DIV12 to DIV25 neurons, we observed a general increase in features associated with synaptic protein cluster size in the synthetic images, while the density of clusters was less affected. We validated the plausibility of these changes and found that the trajectories measured in the synthetic images closely matched the distributions of synaptic features observed in the real data (Fig. 5b, Supplementary Fig. 27). We next evaluated whether the features of generated images fell within the distribution of real data points (Methods). We found that the percentage of images scored as out-of-distribution, based solely on their measured features (*e.g.*, area, density, etc.), is similar between real and synthetic data (4.48% and 5.72% respectively; Supplementary Table 12). This consistency indicates that the generated images accurately reflect the feature distribution characteristics found in the real images of DIV12 and DIV25 neurons.

**Figure 5:**
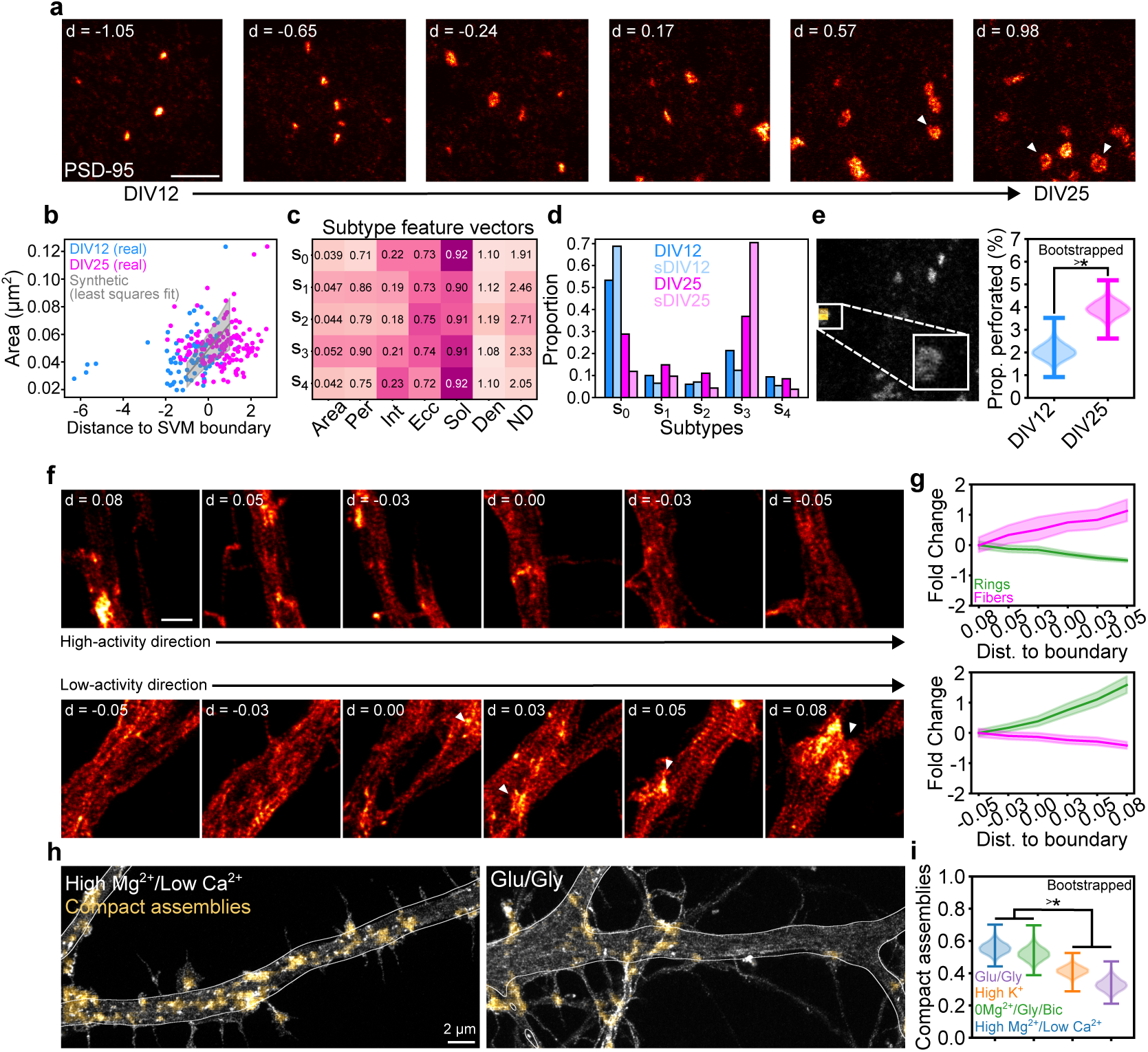
STED-FM latent features encode subtle phenotypic modulation for biological discovery. **a)** Latent attribute manipulation trajectory of PSD95 images from DIV12 neurons, manipulated towards images of DIV25 neurons. **b)** Raw image feature values for the area of synaptic protein cluster as a function of distance to the SVM decision boundary for real DIV12 (blue)and DIV25 (fuschia) images. The grey line represents the average fit obtained by fitting a linear function on all synthetic trajectories (shade: standard deviation). **c)** Average hand-crafted feature vectors of the different PSD95 subtypes found in synthetic data. **d)** Proportion of PSD95 subtypes in the data of DIV12 (blue; real: DIV12; synthetic: sDIV12) and DIV25 neurons (fuschia; real: DIV25; synthetic: sDIV25). **e)** Example of a perforated synaptic protein cluster found using STED-FM. The proportion of perforated synapses is greater in DIV25 neurons compared to DIV12 neurons (Methods). *p*-value = 1.7918 *×* 10^−9^. **f)** Latent decoding from low- to high-activity and high- to low-activity in F-actin images. Example of a trajectory in the high-activity (top) and low-activity (bottom) directions from an anchor image. **g)** Quantification of the presence of F-actin rings (green) and fibers (magenta) depending on the direction (top: to high-activity; bottom: to low-activity). **h)** Quantification of compact assemblies from the images using a similar procedure as in Fig. 2j. **i)** The proportion of compact assemblies (Methods) is reduced with increasing activity. Scale bars: 1 µm unless otherwise noted.

Next, we performed consensus clustering on STED-FM’s representations of generated images to extract synaptic subtypes associated with the PSD95 nanoclusters (Supplementary Fig. 28). We previously showed that analyzing changes in synaptic protein organization at the subtype level provides greater sensitivity and interpretability, as it accounts for the inherent heterogeneity of the feature distributions [55] (Fig. 5b, Supplementary Fig.29; Methods). Consistent with previous biological findings [87], the PSD95 cluster size, and nanodomain number increase for the synthetic images of neurons fixed at DIV25, in comparison to DIV12. Specifically, subtype *s*_3_, characterized by relatively high area, perimeter, and ND became more frequent for later timesteps (Supplementary Fig 29). Subtypes *s*_0_, characterized by smaller area, perimeter, and ND, decrease in proportion for synthetic images of older neurons. We confirmed that the proportion of these subtypes were also similarly modulated in real DIV12 and DIV25 images (Fig. 5d). Finally, visual inspection of the generated trajectories revealed an apparent increase in the number of perforated synaptic proteins in later timesteps (shown with white arrows in Fig. 5a). Leveraging the patch retrieval capabilities of STED-FM, we initially quantified that the prevalence of perforated synapses was indeed higher in the synthetic older neurons (sDIV25 compared to sDIV12; Supplementary Fig. 30). We then validated that this hypothesis held true in the real experimental data, observing a similar modulation in the proportion of perforated synapses (Fig 5e). This increase in the number of perforated synapses during maturation has also been previously reported by other groups, confirming its biological plausibility [88, 89].

We assessed the model’s ability to differentiate phenotypes during the activity-dependent remodeling of neuronal F-actin in dendrites (Fig. 5e & Supplementary Fig. 31), using the segmentation of F-actin rings and fibers for validation [65]. As expected, latent attribute manipulation towards high-activity increases the prevalence of F-actin fibers while it decreases for F-actin rings, and vice versa (Fig. 5e, f). The transition from high- to low-activity revealed regions of high F-actin density, previously reported as compact assemblies in Lavoie-Cardinal *et al.* [65] (Fig. 5f, arrow heads). This observation prompted a quantitative investigation of their abundance in an activity-dependent manner using STED-FM’s patch-retrieval capabilities (Methods, Fig. 5f). The results show that increasing neuronal activity gradually decreases the prevalence of compact assemblies (Fig. 5h,i & Supplementary Fig. 32). Combining STED-FM with the DDPM enables data-driven discovery of overlooked phenotypes without the manual annotations required for supervised analysis [65].

### 2.6 Microscopy automation

In recent years, smart microscopy has become more popular, offering the possibility to adapt microscope acquisition parameters (regions of interest, imaging parameters, reactive microscopy, etc.) to address specific research questions [61, 90–93]. The application of data-driven microscopy generally requires the development of pipelines that can automatically and reliably detect events/structures of interest. We propose to leverage the retrieval capabilities of the STED-FM backbone to automate the region selection process and enable the detection of small substructures like dendritic spines labeled with F-actin. We draw inspiration from feature-based segmentation methods [94] and demonstrate the integration of STED-FM into an automated acquisition pipeline. A microscopist first annotates a few regions of interest (ROIs) on a confocal overview image. Patch-level features corresponding to these annotations are then extracted using STED-FM and used to train a Random Forest classification model (Methods, Fig. 6a, Supplementary Fig. 33). The predicted ROIs are subsequently used for automated region selection in machine learning-assisted parameter optimization pipelines. The annotation pipeline achieves a detection F1-score *>* 0.75 by requiring annotation of only 5% of the image (Methods, Supplementary Fig. 34).

**Figure 6:**
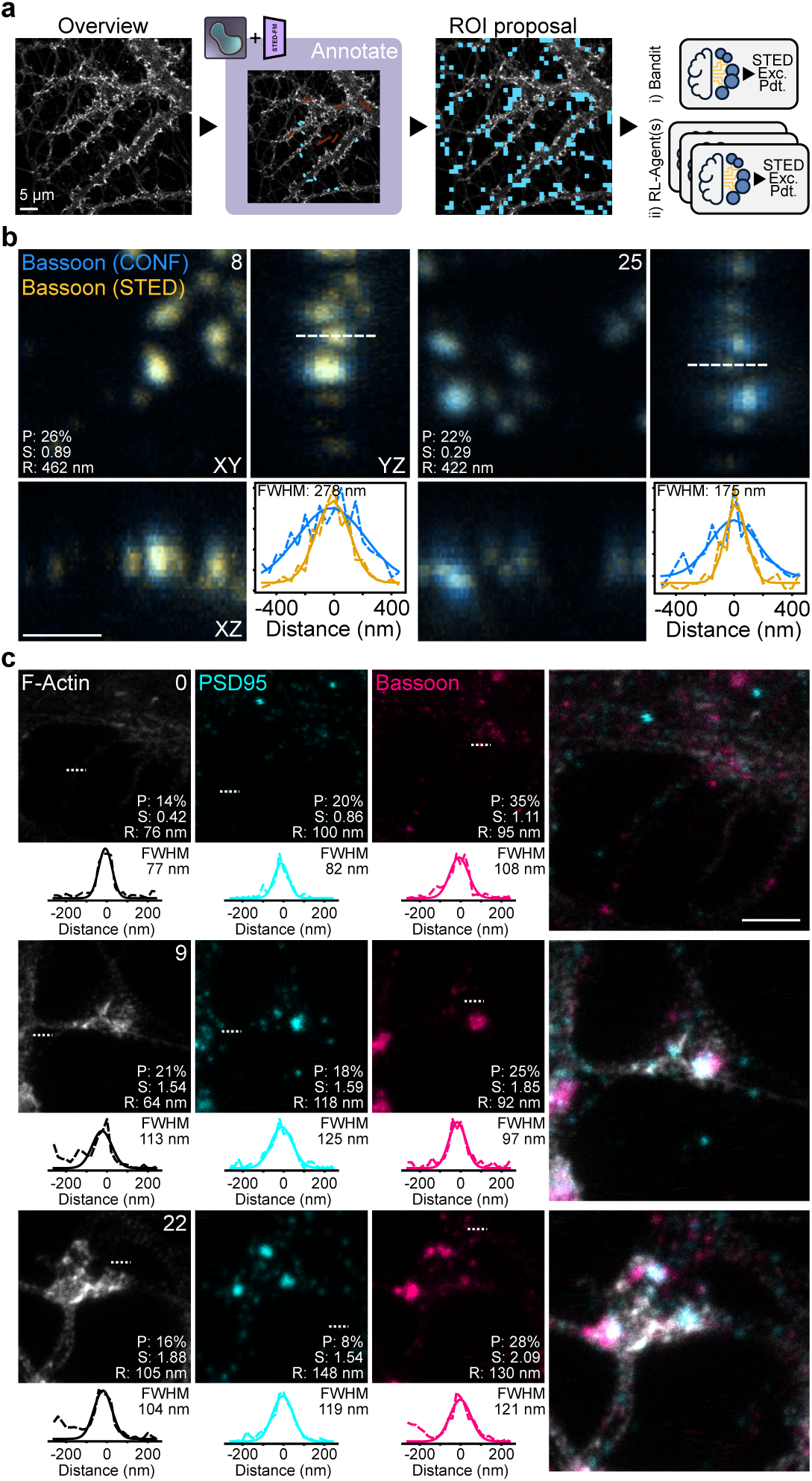
Automation of acquisition pipelines with STED-FM. **a)** Automatic region of interest (ROI) proposal is done by interactively training a Random Forest on the embedded patches from STED-FM on a large confocal overview (cyan: positive annotations; red: negative annotations; see Methods). In this example, spines are detected. The ROIs containing spines are used as part of the optimization pipeline of the imaging parameters. **b)** The imaging parameters of 3D DyMIN acquisition (Excitation and STED laser power, and pixel dwell time) of synaptic protein Bassoon are optimized using the Bandit framework. The imaging optimization objectives improve during the imaging session (P: Photobleaching, S: Signal Ratio, R: z-Resolution). The confocal (Blue) and DyMIN (Yellow) images of Bassoon are overlayed. The XY, XZ, and YZ plane are obtained from maximal projections. FWHM is calculated from manually traced line profiles (dashed). **c)** Imaging parameters (Excitation and STED laser powers, and pixel dwell time) are simultaneously optimized using a multi-agent RL pipeline for a 3-color STED acquisition (F-actin, PSD95, and Bassoon). The imaging optimization objectives improve during the imaging session (P: Photobleaching, S: Signal Ratio, R: z-Resolution). FWHM is obtained from manually traced line profiles shown under each image. Scale bars: 1 µm unless otherwise noted.

We present two imaging pipelines: i) optimization of excitation laser power, STED laser power, and pixel dwell time in a 3D-DyMIN [95] acquisition of the presynaptic protein Bassoon, and ii) a multi-agent reinforcement learning (RL) pipeline for the simultaneous acquisition of 3-color STED images of F-actin, Bassoon, and PSD95. For the 3D-DyMIN acquisition, we used the LinTSDiag procedure proposed in Bilodeau *et al.* [90] to simultaneously optimize three imaging objectives: photobleaching, signal ratio, and resolution (Methods). As expected, the z-resolution of the acquired images improved, reaching values below 200 nm in the final images (Methods, Fig. 6b). This improvement in resolution was achieved at the cost of a reduced signal ratio, while photobleaching was effectively stabilized (Supplementary Fig. 35).

We then adapted the RL pipeline originally proposed in Bilodeau *et al.* [90] to simultaneously run an independent RL model for each fluorophore. At each ROI detected by STED-FM, the RL models selected the set of imaging parameters for their respective fluorophore (Methods). The imaging optimization objectives were evaluated, and the next ROI was processed. For all channels, the models effectively improved the signal ratio while reducing photobleaching and causing a small decrease in resolution (Fig. 6c, Supplementary Fig. 36 & 37).

In a different experiment, the expressive latent representation of STED-FM was leveraged as a training objective within an automated acquisition pipeline. We combined STED-FM’s embedding with a pre-trained SVM, initially trained to classify image quality (Figure 4e-g). By defining the imaging objective simply as the signed distance to the SVM decision boundary, a Bandit optimization algorithm was able to guide the selection of imaging parameters. This approach successfully selected imaging parameters that iteratively improved the quality metric during the imaging session (Extended Data Fig. 3).

Integrating STED-FM’s representations into automated microscopy offers a novel approach to enhance acquisition throughput, enabling automated subregion selection with minimal user interaction and providing a real-time evaluation pipeline for optimization routines.

## 3 Discussion

We introduced STED-FM, a generalist FM for STED microscopy, trained on what is, to our knowledge, the largest STED image dataset compiled to date. This work addresses a central challenge in super-resolution microscopy, the scarcity of large and annotated datasets, by leveraging the abundance of unlabeled data to train large generalizable models [13, 25, 30]. Using SSL, we shift from traditional task-specific training on small curated datasets to the development of generalist models trained on large unlabeled datasets [16]. STED-FM, based on a ViT backbone and trained with the MAE procedure, learns a high-quality latent manifold with expressive and discriminative properties, which translates into strong downstream performance across diverse microscopy tasks, even with minimal supervision.

Our unsupervised analysis of STED-FM goes beyond conventional performance metrics by examining key foundation model capabilities such as the encoding of known physical attributes and the identification of subtle biological patterns. Models pre-trained on unrelated domains such as ImageNet were less effective at capturing these properties due to domain shift [11]. This observation is consistent with recent findings showing that self-supervised pre-training improves both in-domain task performance and generalization in out-of-distribution (OOD) or low-data regimes [36, 96]. These results raise fundamental questions about how domain shifts should be quantified. Although we manually defined domain categories (*e.g.*, ID-STED, OOD-MIC), our quantitative analysis (Extended Data Fig. 2) revealed substantial variability in similarity measures even within the same group, emphasizing the ambiguity of what constitutes a truly OOD task [97]. Traditional deep feature metrics such as the Inception Score (IS) or Fréchet Inception Distance (FID) have been widely used to compare image distributions, yet recent work shows that they can fail to capture diversity and perceptual quality in specific contexts [98, 99]. Moreover, because these metrics rely on feature extrators pre-trained on natural images, they may not be ideally suited for OOD microscopy data [100]. Consequently, the development of new, robust, domain-specific quantification metrics is an open challenge. Such metrics are crucial to accurately estimate fine-tuning data requirements and select optimal pre-training sources for superior downstream performance. We anticipate that our initial efforts at quantitative domain assessment using a metric based on the images power spectrum help lay the foundation for this emerging direction.

We demonstrated the quality of STED-FM’s latent representation across a comprehensive suite of real-world STED microscopy applications, including clustering, retrieval, classification, segmentation, denoising, super-resolution, image generation, and automated microscopy. In data-scarce conditions, models fine-tuned from STED-FM consistently outperformed ViT-S models pre-trained on other microscopy datasets. The high performance achieved by fine-tuning only a linear layer underscores the quality of the learned representations, alleviating the need for extensive annotations and reducing training data requirements for downstream tasks.

Although more complex classification or segmentation heads such as symmetric decoders (Supplementary Fig. 12) could further improve absolute performance, we intentionally evaluated with simple linear layers to better reflect the inherent quality and utility of the STED-FM latent representations. We systematically evaluated classification and segmentation performance across 15 distinct datasets, selected for diversity in: i) imaging modalities (ID-STED, OOD-STED, and OOD-MIC) ii) biological structures, and iii) image quality. Benchmarking on this large and diverse ensemble of datasets enabled us to quantitatively analyze the influence of domain distance on model robustness at an unprecedented scale for super-resolution microscopy. Furthermore, by publicly releasing the results and all associated algorithms and models, this benchmark paves the way for researchers aiming to develop and validate novel, general algorithms or datasets, for future FMs in microscopy.

We showed that STED-FM’s representations can be leveraged for image restoration tasks, including denoising and algorithmic super-resolution. Furthermore, these representations enabled the fine-tuning of a DM through its sampling procedure, producing outputs that align more closely with expert assessments. To our knowledge, this represents the first demonstration of DM fine-tuning in bioimaging, establishing a foundation for large-scale DM that can latter be adapted to a specific task [81]. Additionally, the STED-FM’s representations can be used as a conditioning input for a DDPM to modulate specific features such as image quality and spatial resolution. This manipulation pipeline enables discovery-driven experiments by generating synthetic images that smoothly interpolate between conditions, reflecting realistic experimental outcomes. This capability allowed us to identify subtle PSD95 and F-actin nanostructures that were overlooked during the initial screening, highlighting its potential as a powerful tool for visualizing small structural changes obscured by biological variability. These findings can subsequently guide targeted analyses in real experimental datasets for thorough validation.

Additionally, STED-FM can be integrated into microscopy acquisition pipelines for automated ROI selections with minimal user interactions. The encoding of specific features, such as image quality, within the latent representations of STED-FM can be exploited as a reward signal in machine learning–based parameter optimization loops. Together, these capabilities demonstrate the potential of using FM for fully automated, high-throughput acquisition workflows tailored to specific structures, actively supporting and accelerating the work of microscopists.

In this work, we not only introduced STED-FM but also released the large-scale unlabeled dataset used for its training [51], along with several smaller, curated, and annotated datasets for evaluating performance across different downstream tasks (*e.g.*, semantic segmentation:*SPZ* and *F-actin conformations dataset*, denoising: *VGAT-LQHQ*, or super-resolution: *VGAT-SR* and *Gephyrin*). By making these resources publicly available, we aim to facilitate their broad adoption by the bioimaging community and integration into diverse training pipelines. Combining the STED-FM dataset with external microscopy datasets could enable the creation of multimodal training repositories encompassing multiple microscopy modalities and even extending to timelapse imaging [101]. Increasing dataset diversity by incorporating images from complementary techniques would further strengthen the development of foundation models in bioimaging. This is supported by our results showing a correlation between dataset diversity and performance on downstream classification tasks (Extended Data Fig. 2) as well as by recent work in computer vision, where, for instance, Oquab *et al.* [59] demonstrated that larger and more heterogeneous pretraining datasets improve transfer performance and robustness across downstream tasks. Our adaptation of the power spectrum analysis to evaluate image diversity could also contribute to improve the screening of large bioimaging datasets to automatically identify and add groups of images that significantly contribute to the overall diversity of the dataset.

Although systematic screening of large-scale super-resolution microscopy datasets represents a promising strategy to enhance pre-training diversity, such aggregated datasets remain scarce. For instance, in this work we aggregated SIM images from four publicly available datasets but only managed to obtain 3656 images, a number that falls well short of the large datasets available for conventional fluorescence microscopy, such as *JUMP* or *HPA*. This disparity underscores the critical need for broader dissemination of acquired super-resolution datasets into public repositories [102], as has been successfully achieved in the conventional microscopy community and exemplified here by the release of our STED dataset. The strong cross-domain performance of this dataset on multiple classification and segmentation tasks demonstrates its capacity to generalize beyond STED imaging, suggesting that STED-FM could be integrated into a broad range of quantitative bioimaging pipelines.

We envision a continuous improvement of STED-FM through the continuous aggregation of newly acquired STED images into its training dataset. This iterative scaling of the training data holds the potential to unlock even greater performance gains, aligning with the scaling hypotheses observed in natural language processing and natural image computer vision [1]. By fostering community-driven expansion, we anticipate further improvements to STED-FM’s robustness, generalizability, and overall performance, creating an even more powerful tool for the entire super-resolution microscopy field. As the amount of publicly available data grows, pre-training on this larger scale will enable the training of even larger models, which have been shown to have higher predictive power and robustness [103].

## Acknowledgments

We thank the Neuronal Cultures Platform of the CERVO Brain Research Center for the preparation of the dissociated hippocampal cultures, and the Digital Research Alliance of Canada for the compute resources that were used to train and validate the deep learning models. We thank Renée Hložek for fruitful discussion on the development of foundation models and William Witteman for careful proofreading of the manuscript.

Funding was provided by grants from the Natural Sciences and Engineering Research Council of Canada (RGPIN-2017-06171, P.D.K., and RGPIN-2019-06704, F.L.C., and RGPIN-2019-06706 to C.G.), Fonds de Recherche Nature et Technologie (FRQNT) Team Grant (2021-PR-284335 to F.L.C, C.G., and P.D.K.), Canadian Institutes of Health Research (CIHR) (202109PJT-471107-NSB-CFBA-12805 to F.L.C. and P.D.K.), Neuronex Initiative (National Science Foundation 2014862, Fond de recherche du Québec - Santé 295824 to F.L.C. and P.D.K.), CERVO Brain Research Center Foundation (F.L.C.), the Canadian Foundation for Innovation (32786 to P.D.K. and 39088, F.L.C.), a New Frontiers in Research Fund Exploration Grant (NFRFE-2020-00933 to F.L.C., and C.G.) F.L.C. is a Canada Research Chair Tier II (CRC-2019-00126, F.L.C.) and C.G. is a CIFAR AI-Chair.

A.B. and K.L. were supported by scholarships from NSERC. K. L. was supported by a scholarship from the Institute for Intelligence and Data. F.B. was supported by scholarships from FRQNT, NeuroQuébec, and the Québec Bio-Imaging Network. A.B., R.B., K.T. and F.B. were awarded excellence scholarships from the FRQNT strategic cluster UNIQUE.

## Author contributions

A.B. and F.B. developed the pipeline for the foundation model design and training. A.B. and F.B. performed the unsupervised, supervised post-training, image generation, latent attribute manipulation, and automated microscopy experiments. A.B., F.B., and F.L.C designed the user studies. J.C. acquired and annotated the F-actin conformations dataset. K.T. prepared the samples and acquired the denoising and algorithmic super-resolution datasets of Gephyrin and VGAT in brain slices. J.M.B and F.L.C. designed the experimental protocol and acquired the synaptic development dataset. R.B. managed the Synaptic Protein Zoo platform and aggregated the data and annotations for the corresponding dataset. A.D. acquired the image resolution dataset. K.L. performed preliminary self-supervised learning experiments. A.B., F.B., and F.L.C wrote the manuscript. P.D.K, C.G and F.L.C co-supervised the project.

## Data availability

The in-house datasets (*Synaptic protein zoo* [64], *Synaptic development dataset* [86], *F-actin conformations dataset* [66], *VGAT-LQHQ* [104], *Gephyrin* [104], *VGAT-SR* [105], and the *STED-FM dataset* [51]) are publicly available from https://github.com/FLClab/STED-FM. Other datasets used in this study are available from the original publications. A comprehensive collection of the information pertaining to this publication is available on the accompanying website: https://flclab.github.io/stedfm/.

## Code availability

All the code used in this manuscript is open source and available at: https://github.com/FLClab/STED-FM.

## 4 Methods

### 4.1 Datasets

#### 4.1.1 Self-supervised pre-training datasets

The *STED-FM dataset* [51] consisted of 37 387 images of varying size which were split into 224 × 224 crops. The resulting size of the dataset was 976 022 crops, all of which were used for pre-training STED-FM. Within the *STED-FM dataset*, the protein of interest is known in 238 683 crops. In total, 24 different proteins were identified in this subset, and the total number of crops per protein is reported in Supplementary Table 13.

We benchmarked STED-FM against pre-training of ViT-S on four other datasets which cover the natural image, microscopy and super-resolution microscopy domains. In order of similarity to the STED image domain: Structured Illumination Microscopy images (*SIM* ; [15]), the Human Protein Atlas (*HPA*; [54]), a subset of the JUMP-CP dataset (*JUMP* ; [27]), and *ImageNet-1k* [6]. For *ImageNet*, we did not re-train our own ViT-S model on the ImageNet-1k dataset, but used the pre-trained weights available from the Pytorch Image Models library [106].

For *SIM*, we used all the reconstructed images from the datasets used in [15]. The dataset is aggregated from the 3D-RCAN dataset [107, 108], the BioSR dataset [109, 110], DeepBacs dataset [111–113] and the EMTB dataset [114, 115].]

In the case of *HPA*, we treated the 4 channels of an image as 4 independent images. We performed random cropping to sample 224 × 224 crops from the 512 × 512 original images during training.

For *JUMP*, we used the JUMP Cell Painting datasets [27], available from the Cell Painting Gallery [116] on the Registry of Open Data on AWS. We sampled 3.7M images from the different cpg-0016 sub-folders. Since images were of variable sizes, we performed random cropping of the images during training to obtain crops of size 224 × 224.

Each image in the dataset was normalized to the [0, 1] range using min/max values (*JUMP*, *HPA*), or the 0.001 and 0.999 quantile (*STED-FM dataset*, *SIM* ). The number of independent images and the number of crops for each dataset can be found in Supplementary Table 1.

##### Size-matched datasets

We trained ViT-Small on HPA and JUMP with a dataset size equal to that of STED-FM. This downsamples JUMP from 3.7M images and HPA from 1.1M images to 976 022, exactly (the size of STED-FM’s pre-training dataset). Images were selected randomly. We did not alter the pre-training of ImageNet, as we use the publicly available weights from torchvision [117]. We also elected not to downsample every dataset to the SIM dataset size of 205 946 considering that the first downsampling step had already altered significantly the performance of the JUMP pre-training.

#### 4.1.2 Supervised downstream datasets

For the different downstream experiments performed, we used a total of 20 super-resolution microscopy datasets and 7 confocal microscopy datasets.

##### Classification

We used 9 downstream classification datasets: 2 in-distribution (ID-STED), 2 out of distribution STED (OOD-STED), and 5 out of distribution microscopy (OOD-MIC) (Supplementary Table 3).

- *STED optimization* (*SO* ; ID-STED): Dataset of STED microscopy images from Durand *et al.* [61] containing 4 different structures: the cytoskeleton proteins F-actin and Tubulin, the calcium-dependent protein kinase II (CaMKII), and the post-synaptic protein PSD95. Only the images with an image quality above 0.7 were used. The quality rating was obtained by a manual expert annotation in Durand *et al.* [61]. The train and test splits from the original publication were used. A validation dataset was created by sampling 10% of the training images from each class. The downstream task consisted in classifying the four aforementioned structures (Extended Data Fig. 1).
- *Neural activity states* (*NAS* ; ID-STED): STED microscopy dataset of PSD95 imaged under 4 different experimental conditions from Wiesner *et al.* [55]. It highlights activity-dependent remodeling of PSD95 following different stimulations: High Mg^2+^/Low Ca^2+^ (low activity), 0Mg^2+^*/Gly/Bic* (cLTP), Glutamate/Glycine (cLTD), and Tetrodotoxin (upscaling). The dataset was split into train (70%), validation (15%) and test (15%) sets. The downstream task consisted in classifying the four experimental conditions: low activity, cLTP, cLTD, and upscaling (Extended Data Fig. 1).
- *Peroxisome* (*Px* ; OOD-STED): STED microscopy dataset of images containing peroxisome organelles from de Lange *et al.* [62][118]. It contains images of yeast cells grown for four different time periods in a medium containing 0.5% methanol (MeOH): 4hMeOH, 6hMeOH, 8hMeOH, and 16hMeOH. The dataset was acquired in triplicate. We combined two replicates for training and validation, with a random split of 80%/20% respectively. The third replicate was used for testing. The downstream task consisted in classifying the four time periods (Extended Data Fig. 1). Images were padded with zeros to match the required image size of 224 × 224 pixels.
- *Polymer rings* (*PR*; OOD-STED): STED microscopy images from Hurtig *et al.* [63][119] showing two proteins which self-assemble into filamentous polymers, and which are part of the cell division (Cdv) system: CdvB1 and CdvB2. We randomly split the dataset into training, validation, and testing (respectively 70%, 15%, 15%). The downstream task consisted in classifying CdvB1 vs. CdvB2 (Extended Data Fig. 1). Images were padded with zeros to match the required image size of 224 × 224 pixels.
- *DL-SIM* (OOD-MIC): SIM images of four different sub-cellular structures obtained from Jin *et al.* [120]: Microtubules, Adhesions, Mitochondria, and F-actin. We used the high-quality reconstructed SIM images and the training and testing dataset splits from the original publication. We subsampled 30% of the training dataset to obtain a validation dataset. The downstream task consisted in classifying the aforementioned structures (Extended Data Fig. 1).
- *BBBC026* (OOD-MIC): We used image set BBBC026v1 [121], available from the Broad Bioimage Benchmark Collection [122]. The classification task consisted in predicting the negative and positive controls. Images were randomly split into training, validation, and testing (respectively 70%, 15%, 15%). Crops of 224 × 224 pixels were extracted from the images without overlap for training.
- *BBBC052* (OOD-MIC): We used image set BBBC052v1 [123], available from the Broad Bioimage Benchmark Collection [122]. This dataset was acquired on a confocal microscope. The downstream task was a four-class classification task: CK-666/KD, Cofilin1KD/KD, PFN1KO/KO, TBeta4KD/KD. Images were randomly split into training, validation, and testing (respectively 70%, 15%, 15%). Crops of 224 × 224 pixels were extracted from the images without overlap for training.
- *BBBC053* (OOD-MIC): We used image set BBBC053v1 [123], available from the Broad Bioimage Benchmark Collection [122]. This dataset was acquired on a confocal microscope. The downstream task consisted in classification of images of mitochondria treated with either DMSO or CCCP to induce and detect changes in mitochondrial morphology. Images were randomly split into training, validation, and testing (respectively 70%, 15%, 15%). Crops of 224 × 224 pixels were extracted from the images without overlap for training.
- *HPA-Classification* (OOD-MIC): We used the classification labels provided as part of the Human Protein Atlas Image Classification competition [124]. The downstream task was a multi-label, multi-class classification task with 28 possible labels. Images were randomly split into training, validation, and testing (respectively 70%, 15%, 15%). We performed random cropping to sample 224 × 224 crops from the 512 × 512 original images during training. Only the green channel was used for this task. Following the evaluation procedure of the competition, we report the F1-score as a performance metric instead of the classification accuracy.

##### Segmentation

We used 6 downstream segmentation datasets : 2 ID-STED, 2 OOD-STED, and 2 OOD-MIC (Supplementary Table 4). Example images and ground truth segmentations are shown in Supplementary Fig. 9.

- *F-actin* (ID-STED): STED microscopy dataset of images containing two segmentation classes: F-actin rings and F-actin fibers from Lavoie-Cardinal *et al.* [65]. The training, validation, and test splits from the original publication were used. The downstream task consisted in segmenting regions containing F-actin rings and/or fibers (Extended Data Fig. 1). The ground truth segmentation masks consist of polygonal bounding boxes around the structure of interest, making them weak labels.
- *Synaptic Protein Zoo* (*SPZ*, ID-STED): STED microscopy dataset of PSD95 and Bassoon images from Wiesner *et al.* [55] were annotated using the citizen science platform *Zooniverse* [64]. The ground truths are semantic segmentation masks obtained from the majority class defined from 10 annotations per synaptic nanocluster for each task (segmentation and classification). The segmentation masks correspond to the aggregation (pixel-wise) of all voted masks, defining a probability at every pixel. Not all synaptic proteins in the images have been annotated, resulting in weak labels. The classes correspond to nanocluster morphology: round, elongated, perforated and multi-domain proteins. The downstream task consisted in the semantic segmentation of the four nanocluster morphologies (Extended Data Fig. 1).
- *Foot processes* (*FP* ; OOD-STED): STED microscopy dataset of podocyte foot processes from Unnersjö-Jess *et al.* [68]. The ground truth masks are two-channel segmentations of foot process and slit diaphragm regions. We used the training and testing splits provided by the authors of the original publication. We subsampled 20% of the training set to obtain the validation dataset. The downstream task consisted in the semantic segmentation of foot process and slit diaphragm regions (Extended Data Fig. 1).
- *Lioness* (OOD-STED): STED microscopy dataset of structures and their boundaries acquired using the LIONESS procedure from Velicky *et al.* [29][125]. The four imaging volumes from the original publication were split into two volumes for training, one for validation, and one for testing. The downstream task consisted in semantic segmentation of structures and their boundaries (Extended Data Fig. 1).
- *Lacunar Canalicular Network* (*LCN* ; OOD-MIC): Confocal image stacks of the lacunar canalicular network in cortical rat bone [126, 127]. The downstream task consisted in a 3-class segmentation of the canalicui, lacunae, and canals. We did not use all frames from the confocal stack to create the dataset splits, skipping every 10 frames to simulate a small dataset. We split the dataset into training, validation, and testing splits at the stack level (respectively 70%, 15%, 15%) and extracted crops of 224 × 224 pixels.
- *DeepD3* (OOD-MIC): We used the dataset from DeepD3, an open deep learning-based framework to robustly quantify dendritic spines in microscopy data [128, 129]. The downstream task consisted in a 2-class segmentation of the dendrite and the spine. We used all frames from the dataset. We used the training split provided by the authors for the training and validation datasets (respectively 90% and 10%). The validation split provided by the authors was used for testing. We cropped all frames to 224 × 224 pixels without overlap, making sure that a crop contained more than 1% of foreground. The foreground was obtained by Otsu thresholding.

##### Denoising

We used 4 OOD-STED downstream denoising datasets (Supplementary Table 7). Example of pairs of low SNR and high SNR images are shown in Supplementary Fig. 17-19.

- *ω-tubulin* (OOD-STED): We used the STED denoising dataset of *ω*-tubulin dataset published by Osuna-Vargas *et al.* [130][131]. During training we split the training dataset into training and validation (respectively 90% and 10%). We use the testing split provided by the authors. The images in the dataset are 256 × 256 pixels, we crop the image to 224 × 224 pixels from the top left corner. We registered the low SNR image on the high SNR version with translation using the SimpleITK library [132].
- *ε-tubulin* (OOD-STED): We used the STED dataset of *ε*-tubulin provided in [79]. As in the original publication, low SNR frames were summed to generate the high SNR image. Prior to summing, low SNR images were registered to one another using a rigid body transformation. The StackReg plugin implemented in ImageJ was used for registration [133]. The images were cropped to 224 × 224 pixels. We kept 1200 crops for training as in the original publication and used 100 crops for validation and 100 crops for testing.
- *VGAT-LQHQ* (OOD-STED): We acquired pairs of low and high SNR STED images on our Abberior Infinity line microscope, a different microscope than the one used for the generation of the STED-FM dataset [104]. The Vesicular Inhibitory Amino Acid Transporter (VGAT) in fixed acute mouse brain slices. The low and high SNR STED image acquisition only differed in the number of accumulation: 1 accumulation for the low SNR image and 10 accumulation for the high SNR STED. We registered the low SNR image on the high SNR version with translation using the SimpleITK library [132]. Images were then cropped to 224 × 224 pixels for training.
- *Gephyrin* (OOD-STED): We acquired pairs of low and high SNR STED images on our Abberior Infinity line microscope, a different microscope than the one used for the generation of the STED-FM dataset [104]. The synaptic protein gephyrin was immunostained in fixed acute mouse brain slices. The low and high SNR STED image acquisition only differed in the number of accumulation: 1 accumulation for the noisy image and 10 accumulation for the high SNR STED. We registered the low SNR image on the high SRN version with translation using the SimpleITK library [132]. Images were then cropped to 224 × 224 pixels for training.

##### Super-Resolution

We used 2 OOD-STED downstream super-resolution datasets (Supplementary Table 7)

- *VGAT-SR* (OOD-STED): We acquired pairs confocal and STED images on our Abberior Infinity line microscope, a different microscope than the one used for the generation of the STED-FM dataset [105]. The *VGAT-SR* dataset was acquired on the same samples as the *VGAT-LQHQ* dataset. The low resolution images in this dataset correspond to the confocal images of the *VGAT-LQHQ* dataset. We registered the confocal image on the STED with translation using the SimpleITK library [132]. Images were then cropped to 224 × 224 pixels for training.
- *F-actin-SR* (ID-STED): We used the publicly available dataset from Bouchard *et al.* [80] containing pairs of confocal and STED images of the protein F-actin. We used the provided dataset splits for training, validation, and testing. The training and validation datasets are already provided as crops. For the testing dataset we extracted 224 × 224 pixels size crops from the image without overlap. We ensured that at least 10% of foreground was found within the image. For all crops, we normalized crops (confocal and STED) using a min/max normalization.

##### Latent attribute manipulation

For the latent attribute manipulation experiments (including those from Section 2.5), we used variants of datasets introduced beforehand, as well as new ones. Each dataset was split into training, validation, and test sets. The train and validation sets were used for training the SVM and obtaining the normal vector to the decision boundary. Afterwards, we sampled images from the test set to serve as anchors to the manipulation trajectories.

- *F-actin to tubulin dataset* : To train the SVM, we sampled all F-actin and tubulin images from the *SO* dataset with image quality scores above 0.7 [61]. This resulted in 1 101 images for training, 111 images for validation, and 60 F-actin test images as anchors (Fig. 4d).
- *Image quality dataset* : The ground truth quality scores (ranging from 0 to 1) for this dataset were obtained from the annotations of the original dataset in Durand *et al.* [61]. A small convolutional neural network was trained to assign a quality score to images of F-Actin. We used this network to measure the quality of the generated images. Images with quality scores lower than 0.60 were assigned to the low quality class, while those with scores higher than 0.70 belonged to the high quality class. Images with scores between these thresholds were discarded to reduce the ambiguity near the SVM’s decision boundary. This resulted in 1,623 images for training, 147 for validation, and 40 low quality F-actin images as candidate anchors (Fig. 4e-g). We discarded anchors with a quality score lower than 0.2, as we found those to be too far out-of-distribution for the algorithm to work as desired. This resulted in 27 anchors used for the analysis.
- *Image resolution dataset* : The image resolution dataset consisted of images of the pre-synaptic protein Bassoon (STAR635P) imaged using two different STED laser powers [134]. The high-resolution images were acquired with a STED depletion power of 67.7 mW and the low-resolution images with a depletion power of 22.4 mW. The resulting images had average resolutions of 86 nm (high resolution) and 117 nm (low resolution; Supplementary Fig. 38). The dataset consists of 97 training images, 23 validation images, and 12 low resolution image anchors (Fig. 4h-j). The image resolution was measured using the decorrelation analysis [135].
- *Synaptic development dataset* : The synaptic development dataset consisted of STED images of PSD95 from neuronal cortical cultures fixed at : 12 and 25 days in vitro (DIV12, DIV 25) [86]. We used the DIV12 and DIV25 classes for the latent attribute manipulation. We filtered out images with low signal (foreground intensity *<* 0.15 after min-max normalization). The final dataset resulted in 311 images for training, 144 images for validation, and 62 images of the DIV12 class as anchors (Fig. 5a-e).
- *Activity-dependent F-actin dataset* : We used the F-actin activity-dependent dataset provided in Bilodeau *et al.* [20] for training the SVM. For the latent attribute manipulation experiment (Fig. 5f,g) we used the images of DIV13 neurons treated with a High Mg^2+^/Low Ca^2+^ solution (low activity) and a glutamate/glycine solution (high activity). For the quantification of the prevalence of compact assemblies in F-actin stained images, we used all images from DIV13 neurons provided in Lavoie-Cardinal *et al.* [65] (Fig. 5f-i).

##### Image retrieval

For the quantification of the image retrieval experiments, we used a dataset of F-actin images (Fig. 2j).

- *F-actin conformations dataset* : The F-actin conformations dataset consists of F-actin STED images from several rat hippocampal neuronal cultures fixed between DIV12 and DIV15 (79 images) [66]. Nanoclusters are defined as very dense small round F-actin aggregation (around 100 nm in diameter) along the dendritic shaft. Compact assemblies are large dense F-actin aggregations that are on the dendritic shaft. Spines are identified as protrusions that extend from the dendritic shaft.

### 4.2 Deep neural network pre-training

#### 4.2.1 Vision Transformers - Masked autoencoders

The vision transformers (ViTs) were pre-trained on microscopy datasets using the masked autoencoding procedure introduced in [37]. In Supplementary Fig. 1, we show the training of ViTs of different architecture sizes: tiny, small, base, and large. In all cases, a patch size of 16 was used. We used the architectures provided in the PyTorch Image Models library [106]. According to the results obtained, we selected the ViT-S architecture for STED-FM as it offered the best overall performance. Given an input size of 224 × 224, this resulted in a sequence length of 196 for all models, excluding the class token. All models were trained for 1000 epochs with a batch size of 1024. We used the same training hyper-parameters as in the seminal implementation of the masked autoencoder (MAE) [37]. We used the AdamW optimizer [136] with a learning rate of 1.5 *×* 10^−3^, weight decay of 0.05, and decay rates *ε*_1_ and *ε*_2_ of 0.9 and 0.95, respectively. We used a cosine annealing scheduler with a period of 20 epochs. For the configuration of the MAE, we used a masking ratio of 0.75 and a fixed decoder embedding dimension of 512 for all models. After pre-training, the lightweight decoder of the MAE procedure was removed and the ViT encoder served as the vision foundation model.

### 4.3 Unsupervised analysis

#### 4.3.1 ViT attention maps

To extract an attention map from the pre-trained ViT-S, the self-attention of the class token from the 6 attention heads in each of the 12 transformer blocks was averaged, resulting in 12 attention maps. The global attention map of the model was obtained by summing these 12 maps. A 0.5 quantile threshold was applied to the sum, and the thresholded attention map was then rescaled to input size. The global attention maps were overlayed on the input intensity image and shown to domain experts in a user study (Fig. 2a, b).

#### 4.3.2 Ridge regression

We trained a ridge regression model using the scikit-learn library [137] to predict the spatial resolution, or image quality, given the latent feature vector of an image (Fig. 2c). A ridge regression model was trained independently for each pre-training dataset. A value of *ω* = 1 was used for the regularization term (l2-norm).

#### 4.3.3 Recursive consensus clustering

We performed clustering on the deep feature space of STED-FM for the *NAS* dataset. We followed the work of Sonpatki & Shah [56], which is a recursive version of the consensus clustering algorithm introduced in Monti *et al.* [57].

##### Consensus

Consensus clustering is a method which performs *n* trials of a clustering algorithm. In each trial we sample a subset representing ratio *P* of the dataset. These *n* resampling trials were performed for *C* different clustering algorithms or variants of the same clustering algorithm (such as K-Means with different values of *K*), such that the final number of trials is given by *N* = *n* × *C*. At the end of the trials, for every possible pair of elements, the number of times the pair clusters together is divided by the number of trials in which the two elements were both part of the sampled subset. This produces a consensus matrix *M* . For our experiment, we used *N* = 30, *P* = 0.8, and used the K-Means clustering algorithm with values of *K* ranging from 2 to 10 (*C* = 8).

##### Recursion

We determined the best number of clusters for recursion by computing the consensus matrices for the different values of *K*. We calculated how close each matrix is to a perfect consensus matrix (one which would have all elements either 0 or 1). The selected *K^↑^* which produces a matrix closest to a perfect one was chosen as the optimal number of clusters. Recursion was then performed on those *K^↑^* clusters.

##### Stopping

The recursion was performed until all recursion paths were completed. As proposed in Sonpatki & Shah [56], this happens when clustering returns an intra-cluster stability *<* 0.80, or an inter-cluster overlap *>* 0.20. Letting *S* be the set of pairs that cluster together and *W* be the set of pairs that do not, the intra-cluster stability and inter-cluster overlap were defined as:

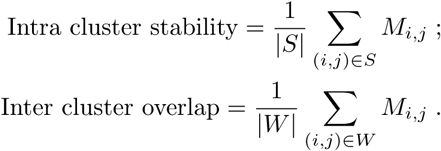

#### 4.3.4 Retrieval Image retrieval

For the image retrieval task, latent embeddings of all images in the ID-STED and OOD-STED classification datasets (*SO*, *NAS*, *Px*, *PR*) were generated using the STED-FM model. Subsequently, an iterative retrieval process was performed. In each iteration, a single image embedding served as the *template*. The cosine similarity was computed between this template embedding and all other image embeddings in the dataset. These similarity scores were then used as prediction scores for performance evaluation.

More specifically, to evaluate the retrieval performance, the Area Under the Receiver Operating Characteristic curve (AUROC [138]) was employed. The ROC curve, in this binary classification context, was constructed by calculating the true and false positive rates across a range of similarity score thresholds (Fig. 2f). For each template, a one-vs-all approach was adopted, where the positive class consisted in images belonging to the same class as the template, and all other images constituted the negative class. The AUROC was calculated for each template and averaged across all images for all classification datasets to provide a single metric performance. On all downstream datasets, the performance of a model was reported as the difference between its performance and the best model’s performance.

##### Patch retrieval

The patch retrieval task follows a similar procedure, but instead of considering image embeddings as templates and candidates, we used patch embeddings, which can be easily retrieved from the ViT architecture. We started by choosing a patch in an image (or a collection of patches) containing a structure of interest, and extracted its embedding from ViT-S as template. In the case of a structure of interest spanning more than a single patch (a region of more than 16 pixels by 16 pixels), we averaged the patch embeddings in which the structure accounts for more than 50% of the patch’s pixels. For all images of a given dataset, we gathered patch embeddings and rank them in order of similarity to the template patch using cosine distance.

We performed the patch retrieval experiment with all pre-trained models, and for a given template, only keep the top 10 candidates from each model (50 images in total from 5 pre-trained models). For quantification we performed a user study in which experts were asked to rank the 50 images (pairwise ranking, corresponding to the 50 most similar embeddings set) according to their similarity to the template image. This produced an expert-defined similarity ranking from which we extracted the top 10 most similar images. We counted the proportion of these 10 candidates which were produced by each of the different pre-trained models (Fig. 2h,i), which was used to quantify patch retrieval performance.

#### 4.3.5 Automatic patch retrieval experiment

The automatic patch retrieval experiment was performed on the *F-actin conformation dataset* [66]. The template was obtained by manually annotating a structure of interest in an image. In the case of a template spanning more than a single patch, we averaged the patches in which the template accounts for more than 50% of the patch’s pixels.

This template embedding was compared to the embeddings of all patches on large field of views. The large field of view was cropped by extracting overlapping crops of 224 *×* 224 pixels (25% overlap). To avoid potential orientation bias of the selected structure of interest, the initial template was augmented through rotation. Four distinct template orientations were generated by applying successive 90*^↔^* rotations. Each of these four rotated template embeddings was then compared against the embeddings of the image crops from the field of view. The maximum similarity score (cosine distance) achieved between any of the four templates and an image patch was used for that patch. For overlapping crops the similarity score was averaged. The resulting maximum projection heatmap was upscaled to input size and overlayed on the large of field of view in Fig. 2j.

The capacity of the pre-trained models to accurately identify relevant regions was evaluated against a manually annotated dataset. The dataset was composed of 79 manually annotated images for three structures: nanoclusters, compact assemblies, and spines. The performance of each pre-trained model was compared to a baseline detection method employing class-specific intensity thresholds. The primary evaluation metric was the difference in precision between a pre-trained model (Pr) and the thresholding method (Pr_thresh._), analyzed over the range of recall. Additionally, the performance of a random baseline was reported. This random model simulated chance-level detection by randomly reassigning the expert annotations within the foreground pixels of the field of view of the image.

### 4.4 Fine-tuning

#### 4.4.1 Classification

Downstream classification performance was evaluated by adding a linear layer with batch normalization on top of the pre-trained encoders. We then either keep the pre-trained features frozen and only fine-tune the linear layer (linear probing), or allow for end-to-end fine-tuning (fine-tuning). All models were trained with the cross-entropy loss. Early stopping was used to checkpoint the best validation loss during fine-tuning. In all classification experiments, training was repeated with five random seeds (42, 43, 44, 45, 46).

##### Linear probing

We use stochastic gradient descent (SGD) with a linear warmup schedule that takes the learning rate from 0.0 to 0.001 in the first 30 epochs. Afterwards, a cosine annealing scheduler [139] with a period of 30 epochs was applied for the remaining 270 epochs, for a total of 300 epochs of fine-tuning. Cosine annealing was done with maximum and minimum learning rate values of 10^−3^ (the value reached after the linear warmup) and 10^−5^, respectively.

##### Fine-tuning

The models were fine-tuned for 300 epochs. The Adam optimizer was used. The scheduler uses a linear warmup schedule that takes the learning rate from 0.0 to 10^−4^ in the first 30 epochs. A cosine annealing scheduler was applied for the remaining training epochs, annealing the learning rate from 10^−4^ and 10^−6^.

##### Random initialization - from scratch

We also trained the models from scratch (random initialization) using a fully-supervised training scheme with the annotated data associated with the downstream task. For this configuration we used the Adam optimizer with a linear warmup scheduler for 30 epochs followed by a cosine decay scheduler over the remaining epochs.

#### 4.4.2 Segmentation

We evaluate segmentation performance by adding a linear layer to the pre-trained encoder. Letting *B* be the batch size, *N_p_*the number of patches, *D_p_*the patch embedding dimension, *P* the patch size (in pixels, assuming square patches) and *C* the number of channels to predict (classes), the encoder outputs a tensor of dimensions (*B, N_p_, D_p_*). The linear layer was trained to map these patch embeddings to pixel values (in the range [0, 1]), transforming the encoder’s output (excluding the class token embedding, hence *N_p_ −* 1 patches) to a tensor of dimension (*B, N_p_* − 1,*P* ^2^*C*). The output of this linear layer was then reconstructed into image space (a process called “unpatchifying”) to match the original input dimensions, enabling the computation of a Mean Squared Error (MSE) loss against the ground truth segmentation mask. For classification, we consider linear probing and fine-tuning. For all training, early stopping was used to checkpoint the best validation loss during fine-tuning. The models were trained for 300 epochs. In all segmentation experiments, training was repeated with five random seeds (42, 43, 44, 45, 46).

##### Linear Probing

When using linear probing, we use SGD with a linear warmup of 30 epochs, followed by cosine annealing for the remaining epochs using maximum and minimum learning rates of 10^−3^ and 10^−5^, respectively, along with a period of 30 epochs.

##### Fine-tuning

For end-to-end fine-tuning we use the same schedulers, this time removing the 30-epoch period and replacing the SGD optimizer with the AdamW optimizer, using a learning rate of 10^−3^, weight decay of 0.1 and *ε* parameters of 0.9 and 0.95.

##### Random initialization - from scratch

When training from scratch, only the hyper-parameters of the AdamW optimizer were different from the end-to-end case, with learning rate of 10^−3^, weight decay of 0.05, and *ε* parameters of 0.9 and 0.999.

##### UNETR decoder

For some experiments, we used a larger decoder than a simple linear layer in order to maximize performance gains. The decoder used closely follows the UNETR architecture [140]. It was used in segmentation, denoising and algorithmic super-resolution experiments (Supplementary Fig.17-21). The performance gains obtained from using this large decoder rather than the simple linear layer for segmentation are also shown in Supplementary Fig. 12.

#### 4.4.3 Small data regime

For both classification and segmentation, we also considered the small data regime, in which the number of annotated instances for fine-tuning was limited. For classification, we used *{*10, 25, 50, 100*}* annotated instances per class for the linear probing or fine-tuning steps. For datasets with insufficient instances to meet these counts, the maximum available number of instances per class was used (Supplementary Table 3). For segmentation tasks, we considered a percentage of training samples rather than a raw count, using a ratio of *{*0.01, 0.1, 0.25, 0.50*}* of available annotated samples for linear probing or fine-tuning steps.

To reduce variability in performance caused by different number of weight updates when using smaller datasets, we adopted a constant training budget. The budget was set to match the number of parameter updates used in the full dataset fine-tuning scenario, with an upper limit of 1000 training epochs. More specifically, we define the training budget in the full dataset scenario as *B_f_* = *N_t_* − *N_e_*, where *N_t_* is the number of annotated training instances and *N_e_* is the fixed number of training epochs. The small data regime training budget *B_s,_*_classif_ in classification was computed as:

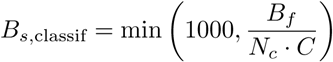

where *N_c_* is the number of training instances per class and *C* is the number of classes. In segmentation, the training budget *B_s,_*_seg_ was computed as:

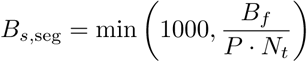

where *P* is the percentage of training samples. The optimizers, loss functions, and hyper-parameters (other than those pertaining to training epochs) were identical to those described in the full-dataset experiments.

### 4.5 Structural complexity and model generalizability

#### 4.5.1 Quantifying domain distance

To establish a quantitative metric for describing the distance between the pre-training dataset and each downstream task dataset, we used a frequency-based approach. Measuring distances between image domains presents an open challenge. Metrics based on deep learning features (*e.g.* FID, IS) are common in computer vision [70–74] but recent work showed that they can fail to capture diversity and perceptual quality in specific contexts [98, 99]. Moreover, because these metrics rely on feature extrators pre-trained on natural images, they may not be ideally suited for OOD microscopy data [100]. Metrics based on direct pixel comparison (*e.g.*, MSE or SSIM) are highly sensitive to spatial shifts and thus unreliable to measure the distance between images with different biological structures.

To assess domain distance between images from different biological structures and acquired with various microscopy techniques, we decided to rather use the frequency content of the images, leveraging a normalized power spectrum analysis. As Koho *et al.* [75] demonstrate, microscopy images have distinct power spectrum signature that can be used to infer the quality. They also point out that spectral domain approaches are more easily applicable to comparing different images, than pixel-wise metrics. The distance between two datasets was quantified by measuring the Kullback-Leibler divergence between the normalized power spectrum density of two images [141, 142]. The distance between two datasets corresponds to the average Kullback-Leibler divergence between all pairs of images. For large datasets we sampled *N* = 5000 images.

#### 4.5.2 Correlation with downstream performance

The power spectrum-based distance was correlated with the downstream model performance across all evaluated tasks (classification and segmentation). To enable the comparison across tasks with different dynamic ranges, both the distance values and the performance metrics (accuracy for classification (F1-score for *HPA-Classification*) and AUPR for segmentation) were Z-score standardized within each specific task.

#### 4.5.3 Evaluating pre-training dataset diversity

As a metric for dataset diversity, we computed the average intra-dataset distance between images, using the power spectrum-based metric detailed above. A larger average intra-dataset distance indicates greater variability in structural texture and spatial frequency content across the pre-training image set. The Pearson correlation coefficient between the dataset diversity and the average downstream performance across all tasks was then computed (accuracy for classification (F1-score for *HPA-Classification*) and AUPR for segmentation). The performance within each downstream dataset were Z-score standardized.

#### 4.5.4 Quantifying segmentation difficulty

To analyze the relationship between structural properties and performance of the downstream segmentation tasks, we incorporated a quantitative measure of segmentation difficulty based on object morphology. Following prior work [69], we used the Perimetric Complexity (PC) as the metric. PC is defined by the following equation:

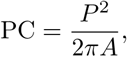

where *P* denotes the perimeter of the segmented structure and *A* denotes its area. Higher PC values correspond to increasingly complex or irregular object morphology.

### 4.6 Image restoration

#### 4.6.1 Denoising

To enable image-to-image denoising, a dedicated decoder was integrated into the STED-FM architecture. The decoder design was heavily inspired by the UNETR (U-Net Transformer) model, adated specifically for 2D segmentation tasks [140].

The decoder structure utilizes skip connections originating from the ViT-small encoder’s intermediate layers, specifically *{z*_3_*, z*_6_*, z*_9_*, z*_12_*}*. These features are projected and concatenated into the corresponding upsampling blocks. The feature map depths in the decoder blocks that process these skip connections were configured with filter counts *{*64, 128, 256, 512*}*, respectively. Each subsequent deconvolution block performs upsampling while systematically halving the number of filters. The final output block consisted of two sequential convolutional layers, followed by a single terminal convolutional layer that projected the feature maps to the desired number of output classes. The model’s final output was then passed through a sigmoid activation function.

During training, the STED-FM encoder was kept frozen. We trained the decoder for 300 epochs with the AdamW optimizer with a weight decay of 0.01 and betas of (0.9, 0.999). The learning rate was set at 1 × 10^−4^ and used a linear warmup schedule that takes the learning rate from 0.0 to 1 × 10^−4^ in the first 30 epochs. Afterwards, a cosine annealing scheduler [139] with a period of 30 epochs was applied for the remaining 270 epochs, for a total of 300 epochs of fine-tuning. Cosine annealing was done with maximum and minimum learning rate values of 10^−4^ (the value reached after the linear warmup) and 10^−6^, respectively.

We compared STED-FM with four baselines: CARE2D [76], Noise2Void (N2V) [77], pix2pix [78], and UNet-RCAN [79]. For CARE2D, N2V, and pix2pix we used the implementation provided in ZeroCostDL4Mic and DL4MicEverywhere [143, 144]. In all cases, we used the default training parameters and trained for 300 epochs. For the UNet-RCAN, we used the training scripts provided in the original publication. We kept the same training parameter configuration and trained for 200 epochs as in the seminal paper.

#### 4.6.2 Algorithmic super-resolution

For simple algorithmic super-resolution task involving the *VGAT-SR* dataset, we used the same training procedure as with the denoising task.

For the algorithmic super-resolution task on the *F-Actin-SR* dataset, we used a denoising diffusion implicit model (DDIM) following the implementation of Dhariwal & Nichol [145]. We used 1000 diffusion timesteps for training the model and 100 denoising steps for the DDIM sampling. We used a linear beta schedule. The denoising network within the DDIM was based on a U-Net architecture with residual connections and attention mechanisms. It consisted of three encoder stages with channel multipliers (1, 2, 4) applied to a base dimension of 64, followed by a bottleneck and three symmetric decoder stages with skip connections. Each encoder and decoder contained two ResNet blocks with RMS normalization and SiLU activation functions, followed by an attention layer. We employed linear attention for computational efficiency in the outer stages and full self-attention in the innermost stage. Timestep information was embedded using sinusoidal positional encodings and processed through a multi-layer perceptron. These timestep embeddings modulate the ResNet block features via affine transformations (scale and shift parameters). We trained for 300 epochs, using the Adam optimizer with a learning rate of 2 × 10^−4^ and beta coefficients of 0.9 and 0.99.

We then fine-tuned the DDIM using the procedure described in DRaFT [81]. Rather than optimizing the standard denoising objective, DRaFT directly optimizes a differentiable reward function by backpropagating through the deterministic DDIM sampling process. We employed the negative mean squared error between STED-FM’s embeddings of the real and generated images as reward signal (Supplementary Fig. 22a). To enable memory-efficient fine-tuning, we applied Low-Rank Adaptation (LoRA) [146] to the linear and convolution layers of the denoising network with a rank of 4, thereby only training a small subset of adapter parameters while keeping the base DDIM frozen. We also used gradient checkpointing during the backpropagation through sampling steps, which reduces memory consumption by recomputing intermediate activations during the backward pass. The total loss to minimize combined the negative reward objective (with a weight of 1.0) with the standard denoising loss (with a weight of 0.1) to maintain training stability. We used 100 deterministic DDIM sampling steps (*ϖ* = 0) and backpropagated through only the last 5 sampling steps (DRaFT-5 variant) to balance computational efficiency with gradient signal quality. Optimization was performed using the Adam optimizer with a learning rate of 1 × 10^−4^ and beta coefficients of 0.9 and 0.99. Training was performed over 15 epochs, which given our dataset of 4331 samples, amounts to 64965 reward queries.

To evaluate the generation of the F-actin nanostructures we used a similar procedure to Bouchard *et al.* [80]. Pixel-wise metrics such as MSE, PSNR, SSIM were used to compare the synthetic images with the real images. To assess the realism and position of the F-actin nanostructures within the images, the pre-trained U-Net model from Lavoie-Cardinal *et al.* [65] was used to predict the presence of two structures: F-actin rings and fibers. The predictions of the model were then compared with that of manual annotations using the Dice score. To assess the visual fidelity of the synthetic images we created a user-study. We showed four experts a true STED image (N=26) as well as three generated images obtained from pix2pix, the pre-trained diffusion model (named DDIM for the sampling procedure used) and the DRaFT-finetuned diffusion model using STED-FM, all conditioned on the confocal version of the original image. The experts were then asked to select which of the three generated images was most similar to the true STED.

### 4.7 Image generation

#### 4.7.1 Latent-guided denoising diffusion probabilistic models (DDPMs)

Following the latent guidance framework of Preechakul *et al.* [84], we conditioned the DDPM using a 384- dimensional latent vector **z** obtained by passing an image through STED-FM. To condition the DDPM, we map **z** to the dimensionality of the timestep embedding **t**’s latent space (256 dimensions). The resulting embeddings were then added (**z** + **t**, Fig. 4a), and serve as the conditioning input. Once the latent-guided DDPM was trained, a noise image **x**_T_ can be given to the DDPM along with the latent code **z** of a test image **x**_0_. The DDPM generates a new image which preserves the features present in image **x**_0_.

For classifier-free guidance [83], a conditioning vector was obtained by creating an embedding layer for 24 protein classes that were defined in a subset of the *STED-FM dataset* (Supplementary Table 13), producing a 256-dimensional vector which matches the timestep embedding dimension. The latent vector was then added to the timestep embedding for conditioning (**z** + **t**, Fig. 4a).

#### 4.7.2 Latent attribute manipulation

For latent attribute manipulation, we followed the method from Shen *et al.* [85], originally developed for generative adversarial networks (GANs) and later adapted to DDPMs by Preechakul *et al.* [84]. The method manipulates image semantics in latent space using the normal vector of a SVM’s decision boundary, trained to separate two classes (Fig. 4c). The SVM’s decision boundary is a hyperplane in d-dimensional space, from which we can extract its normal vector **n** *↔* R*^d^*. For an image embedding **z** in this latent space, we compute the signed distance from **z** to the hyperplane as

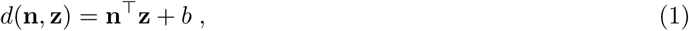

where *b* is the intercept of the decision boundary. To manipulate the image embedding **z** in the direction of the normal vector to a target distance *d_t_* from the hyperplane, we update it along the direction of the normal vector **n**

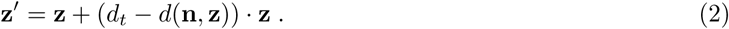

Manipulating image semantics requires partitioning of an image dataset into two classes that represent opposing ends of the desired semantic attributes. In our case, the images were embedded using STED-FM, and a SVM was trained to classify the image embeddings. The direction of the normal vector to the SVM’s hyperplane corresponds to the semantic attribute direction. We can then investigate how the image has changed as a result of the manipulation by decoding the manipulated embedding **z***^↘^* back to pixel space using the conditioned DDPM described in the previous section. In all latent attribute manipulation experiments, we shifted each image’s latent representation from the mean distance to the SVM boundary of one distribution toward the mean distance of the other distribution.

### 4.8 Pattern discovery

#### 4.8.1 Synaptic development of PSD95

To investigate age-dependent changes in PSD95 morphology, we used the *Synaptic development dataset* [86]. A SVM was trained to discriminate between the synaptic structures from cultured neurons fixed at two different time points (Day In Vitro): DIV12 and DIV25. The normal vector to the decision boundary of the SVM in the latent space was then used to manipulate the latent representation of an image of PSD95 corresponding to DIV12 neurons towards images corresponding to PSD95 in neurons at DIV25. For each DIV12 anchor image, we generated a series of six interpolated latent codes at constant intervals between the average distance of the DIV12 and DIV25 classes from the SVM boundary (Supplementary Figure 26). Manipulated latent codes were input to the DDPM to synthesize the corresponding images, which were then subjected to further analysis.

To quantify the morphological changes induced by the latent space manipulation, we first identified protein clusters in the generated images using a wavelet segmentation method, similar to the approach described in Wiesner *et al.* [55]. Several morphological and intensity features were extracted from the segmentation: area, perimeter, eccentricity, solidity, density, intensity, and number of nanodomains. These features were averaged across all identified protein clusters within each image.

To evaluate the plausibility of the features extracted from synthetic images, we calculated the percentage of images whose features were significantly different (*p <* 0.05) from the corresponding target distribution. For example, for both real and synthetic DIV12 images, the target distribution for a given feature was defined as the distribution of that feature’s values in the real DIV12 dataset. The p-value of a data point was obtained from computing its Z-score in the target the distribution (Supplementary Table 12.

To gain a deeper understanding of the features that best discriminate between PSD95 nanostructures at DIV12 and DIV25, we performed a consensus clustering analysis. K-means clustering was applied by varying the number of clusters (*K*) from 2 to 25. A pair-wise consensus matrix was constructed, indicating the frequency with which each pair of data points was clustered together out of the 23 independent K-Means runs. Hierarchical clustering was then applied on the consensus clustering, from which we were able to identify five subtypes of PSD95 synaptic clusters in the images. The proportion of each subtype was measured at each step of the latent space manipulation. To assign one instance to one of the subtypes, we computed the euclidean distance between the image’s and the subtypes’ features and assign the image to the closest subtype (Supplementary Fig. 29).

##### Perforated synapses

We observed an increased prevalence of perforated synaptic structures when manipulating the latent representations of PSD95 clusters towards those of DIV25 neurons. To further characterize this effect, we performed a detailed analysis using the patch retrieval capabilities of STED-FM. A template image was generated from the synapse shown in Fig. 5a (d = 0.98; right image), and the procedure described in Automatic patch retrieval experiment was applied to compute similarity maps across the training data from DIV12 and DIV25 neurons in the *Synaptic development dataset*. To obtain an approximate segmentation and count of perforated synapses, a similarity threshold of 0.65 was applied, and the total number of synapses was determined using the regionprops function from the scikit-image Python library [147]. This threshold was manually validated by an expert. The reported proportion corresponds to the number of detected perforated synapses divided by the total number of synapses identified via wavelet-based segmentation in the PSD95 images of the *Synaptic development dataset*.

We calculated the bootstrap distribution of perforated synapse proportion by resampling the number of perforated synapses found per image 10 000 times. For each resampling iteration, the proportion was recalculated. For statistical testing, we performed a two-tailed z-test with the null hypothesis (*H*_0_) that the mean proportions are equal (*H*_0_ : *µ*_DIV12=_*_µ_*_DIV25_ ). The standard score (*Z*) was calculated as:

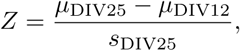

where *s* is the standard deviation. The *p*-value was obtained using 2Φ(*−|Z|*), where Φ is the standard normal cumulative distribution function.

#### 4.8.2 F-actin nanostructures

The activity-dependent remodeling of F-actin from rings to fibers was investigated using the training and validation datasets provided in Bilodeau *et al.* [20]. To ensure unambiguous training, image crops containing both F-actin rings and fibers were excluded from the dataset used to train the SVM boundary.

For image generation, the images of the testing dataset provided in Lavoie-Cardinal *et al.* [65], corresponding to High Mg^2+^/Low Ca^2+^ (low-activity) and Glu/Gly (high-activity), conditions were used as anchors. For each crop in the testing dataset, its latent code was manipulated along the normal vector of the SVM boundary. We generated a series of six interpolated latent codes at constant intervals between the average distance of the low- and high-activity classes from the SVM boundary (Supplementary Fig. 31). Five images were generated for each anchor with uniform step sizes along the manipulation vector using the DDPM conditioned on the manipulated embeddings. To validate the morphological changes, we used the pre-trained U-Net model from Lavoie-Cardinal *et al.* [65] to quantify the evolution of segmented F-actin rings and fibers as a function of image manipulation.

##### F-actin compact assemblies

Given the increased number of compact F-actin assemblies observed when manipulating the latent representations toward the low-activity condition, we performed a more detailed analysis of this pattern using the model’s patch retrieval capability. A template representing a compact F-actin assembly was manually annotated from a real image. Using the procedure detailed in Automatic patch retrieval experiment, we computed similarity maps for this template across the activity-dependent F-actin dataset and applied a threshold of 0.65 on those maps to generate the segmentation masks. The area of compact F-actin assemblies was quantified within dendritic regions identified using the immunostained protein MAP2 as a reference signal. Segmented regions intersecting the dendritic mask were included in the proportion calculation. The bootstrap distribution of the average proportion was calculated using 10,000 resamplings and is reported in Fig. 5i.

### 4.9 Automated Microscopy

#### 4.9.1 STED microscopy experiments

##### STED imaging

Super-resolution imaging was performed on an Abberior Expert-Line STED system (Abberior Instruments GmbH, Germany). This system was equipped with a 100x oil immersion objective lens with a numerical aperture of 1.4 (Olympus, model UPLSAPO100XO), a motorized stage, and an auto-focus unit. Three excitation wavelengths were used (485 nm, 561 nm, and 640 nm) for the experiments. The system is equipped with a 775 nm depletion laser (40 MHz, 1.2 ns pulse duration; MPB Communications). All lasers were operated at 40 MHz. The emitted fluorescence photons were detected by avalanche photodiode detectors (APDs) through a pinhole set at approximately 1 Airy unit. Red dyes were detected with a FF02-615/20-25 filter (Semrock) and far-red dyes were detected with a ET685/70 filter (Chroma). The acquired images were visualized and processed using the FIJI software (ImageJ, version 1.54p) or Python.

##### Neuronal cell culture

Extraction and culture of hippocampal neuronal cells was performed following the protocol of Nault & De Koninck [148] and in accordance to guidelines of the animal care committee of Université Laval. Briefly, neuronal cultures from the hippocampus were obtained using neonatal Sprague-Dawley rats. P0-P1 rats were sacrificed by decapitation before dissection of the hippocampi. Cells were dissociated using enzymatic digestion and mechanical dissociation, and then seeded onto 12 mm coverslips coated with poly-D-lysine and laminin (40,000 cells/coverslip). Neurons were cultivated in a growth medium composed of Neurobasal and B27 (in a 50:1 ratio), enriched with penicillin/streptomycin (25 U/mL; 25 *µ*g/mL) and 0.5 mM L-GlutaMAX (by Invitrogen). After 5 days, Ara-C (5 *µ*M; from Sigma-Aldrich) was added into the medium to limit the proliferation of non-neuronal cells. Cultures were fed twice a week, by replacing *↗*50% of the growth medium with serum- and Ara-C–free medium.

##### Sample preparation and staining procedures

Neuronal cultures were fixed for 20 minutes in 4% PFA solution (PFA 4%, Sucrose 4%, Phosphate Buffer 100mM, Na-EGTA 2mM). Permeabilization and blocking of aspecific binding sites were done simultaneously with a solution of Triton X-100 and 2% goat serum in PBS 20 mM, for 30 min. Primary and secondary antibodies were successively incubated for 2h and 1h, respectively. Labeling of F-actin with fluorescently-tagged phalloidin was performed in a subsequent step, for 2 h. Incubation of antibodies was performed in the blocking solution at room temperature. Immunostained coverslips were mounted in PVA with DABCO for imaging. Supplementary Tables 14 & 15 summarizes the antibodies used in this study with associated concentrations. F-actin was stained with phalloidin-STAR635 (Sigma Aldrich, cat. 30972-20µg, 1:50 dilution).

#### 4.9.2 Workflow

The automated microscopy experiment with automatic ROI selection were performed as follows:

1. A confocal overview image of a large field of view was acquired on the microscope (Fig. 6, Supplementary
2. Regions of interest (ROIs) were identified and extracted from the overview using the developed napari-plugin (Supplementary Fig. 33i). The STED-FM latent representation vectors were extracted from each image patch corresponding to the identified ROIs forming the training dataset for the random forest classifier (Supplementary Fig. 33ii). STED-FM extracts the latent representation vector of all patches in the image and the trained random forest is used to predict the associated class (Supplementary
3. The regions of interest (ROIs) were imaged using the STED microscope, with imaging parameters selected through machine learning–based optimization routines such as bandit optimization or reinforcement learning (RL) models. [90].

#### 4.9.3 Interactive ROI selection

For interactive ROI selection, a random forest model was trained on patch embeddings obtained from the STED-FM backbone of the overview confocal image. This approach is inspired by the work of Seifi *et al.* [94]. The interactive annotation was implemented in a napari-plugin, allowing users to annotate as many structures of interest as desired. In the experiments, the regions containing spines (or not) were annotated.

Since STED-FM requires images of 224 × 224 pixels, the overview image was tiled with a 50% overlap and fed to the model. A 16 × 16 pixel patch within these crops was considered annotated if at least 25% of its area had been annotated. The extracted patch embeddings and corresponding annotations were split into training and validation sets with a 80/20 ratio. The napari-plugin provides user-configurable parameters for the random forest model such as the number and maximum depth of trees. During the experiments, a configuration of 100 trees with a maximum depth of 5 was used.

To extract individual ROIs from the binary masks predicted by the random forest model, the following procedure was applied: 1) the predicted mask was converted into a labeled image, where each contiguous region (island) of positive pixels was assigned a unique ID, 2) the centroid of the bounding box center of each island was obtained by using regionprops from the scikit-image Python library [147]. In the experiments, ROIs of a constant, pre-defined size were used and centered on the centroid of the bounding box.

##### Benchmarking and evaluation

Quantifying the performance of an iterative annotation pipeline is non-trivial, as annotations can be iteratively updated until all ROIs were selected, making direct measurement difficult. To address this, we designed an evaluation strategy that simulated an annotator, allowing us to benchmark the performance of the random forest on a set of unseen images using a leave-on-out cross-validation scheme across five images.

We developed an iterative strategy to efficiently select the most informative training samples from unlabeled image patches, aiming to maximize classification performance while minimizing annotation effort. The process begins by extracting feature embeddings from an image using STED-FM, which then serve as the pool of available patches for training. The training set is initialized with a random 10% sample of positive (spines) patches. The same number of negative patches were added to keep the dataset balanced.

The core of the algorithm is an alternating sampling loop that sequentially adds one hard negative and one hard positive sample. The algorithm prioritizes patches that the current random forest classifier (trained on previously selected data) finds most uncertain or misclassifies (low-confidence predictions). To efficiently guide this sampling, *k*-means clustering is performed on the patch embeddings (16 clusters for positive patches and 64 for negative patches) to capture the heterogeneity within each class, reflecting the need for diverse training examples. Patches from high-confidence clusters were skipped to avoid redundant sampling.

After a new sample is tentatively added, the classifier is retrained and evaluated using the F1-score on the remaining patches. If the new sample improves the F1-score, it is permanently added to the training set. Otherwise, the patch yielding the highest tentative F1-score is selected to guarantee progressive model improvement. The process terminates when the F1-score on the current image reaches 0.9 or the selected number of samples hits 500 patches per image. This adaptive sampling strategy ensures exploration of the feature space by targeting misclassified examples as a user would do.

#### 4.9.4 Bandit optimization

##### Automated ROI selection and 3D acquisition

3D-DyMIN acquisitions were performed on neuronal cultures stained for Bassoon and F-actin. For interactive ROI selection using STED-FM as previously described, spines were chosen from a confocal overview of the F-actin. At each selected location, a volume of 2.88 µm *×* 2.88 µm *×* 2 µm was acquired. Only the STED and excitation powers, and pixel dwell time were optimized. Supplementary Table 16 provides the range of each parameter. DyMIN parameters were set at a threshold of 1 photon detected within 20% of the pixel dwell time for steps 1 and 2 [95]. The imaging optimization objectives were photobleaching, signal ratio and z-Resolution. Photobleaching and signal ratio were calculated following the procedure described in Bilodeau *et al.* [90]. Z-resolution was approximated by fitting a Gaussian function to the line profile obtained from the central slices (10 above and 10 below the center) of the maximal-projection of each frame. During the optimization process, the z-profile was approximated by a center-line across the summed volume, which accounts for the difference between the automatic resolution computation and the manually determined FWHM in Fig. 6b. The contextual LinTSDiag model was used for optimization with the default parameters of Bilodeau *et al.* [90].

##### STED-FM as a metric for image quality optimization

We designed an experiment using STED-FM as an optimization objective to guide image acquisition. To quantify the quality of a newly acquired image, the image was first embedded into STED-FM’s latent space. This latent representation was then passed to the the pre-trained SVM of Figure 4e-g, which was trained to differentiate between low- and high-quality images. A high quality prediction corresponds to an embedding far from the SVM’s decision boundary and having a positive score (*>*0). Acquisitions were performed on neuronal cultures stained for F-actin. ROIs were manually selected for this experiment, and at each location, an image of 4.48 µm × 4.48 µm was acquired. The optimization process focused on adjusting three imaging parameters: the STED power, the excitation power, and the pixel dwell time. The optimization was driven by the LinTSDiag model, using the default parameters from Bilodeau *et al.* [90]. The complete range of values for each parameter is provided in Supplementary Table 17.

#### 4.9.5 Reinforcement learning

3-color STED acquisitions were performed on neurons stained for Bassoon, PSD95, and F-actin. For each channel, a dedicated reinforcement learning (RL) model was used for parameter optimization. Specifically, we employed the same proximal policy optimization (PPO) model described and trained in Bilodeau *et al.* [90]. The range of imaging parameters for each channel is provided in Supplementary Table 18. The imaging session followed the same methodology and default parameters as in Bilodeau *et al.* [90]. Briefly, ROIs were selected using STED-FM as previously described from an initial confocal image, spines were chosen from the F-actin signal. A confocal acquisition of a ROI was then given as input to the corresponding RL model. Based on the acquisition history, the model selected the next imaging parameters, and a STED image was acquired. A second confocal image was then acquired to quantify photobleaching. The imaging optimization objectives were measured from the acquired images and fed back to the RL model to inform future parameter selections. This iterative process was repeated for a total of 30 ROIs with the STED images acquired in this order: F-actin, PSD95, and Bassoon.

### 4.10 Resampling statistics

We employed resampling to verify statistical differences between two groups [149]. The detailed procedure is described in Bilodeau *et al.* [20]. When analyzing more than two groups, the approach involved resampling of the F-statistic for 10,000 iterations [149]. We tested the null hypothesis that all groups originate from the same underlying mean distribution, rejecting it if the calculated p-value is below 0.05. Upon rejection, we conducted pairwise comparisons between each group using a two-group randomization, also performed with 10,000 repetitions [149]. This test randomly reassigned the data points between the two groups being compared and calculated the difference in their means. We then assessed the likelihood of observing our actual mean difference under the assumption of no true difference (the null hypothesis), again using a significance level of 0.05 to determine statistical significance.

## 5 Extended Data

**Extended Data Fig. 1:**
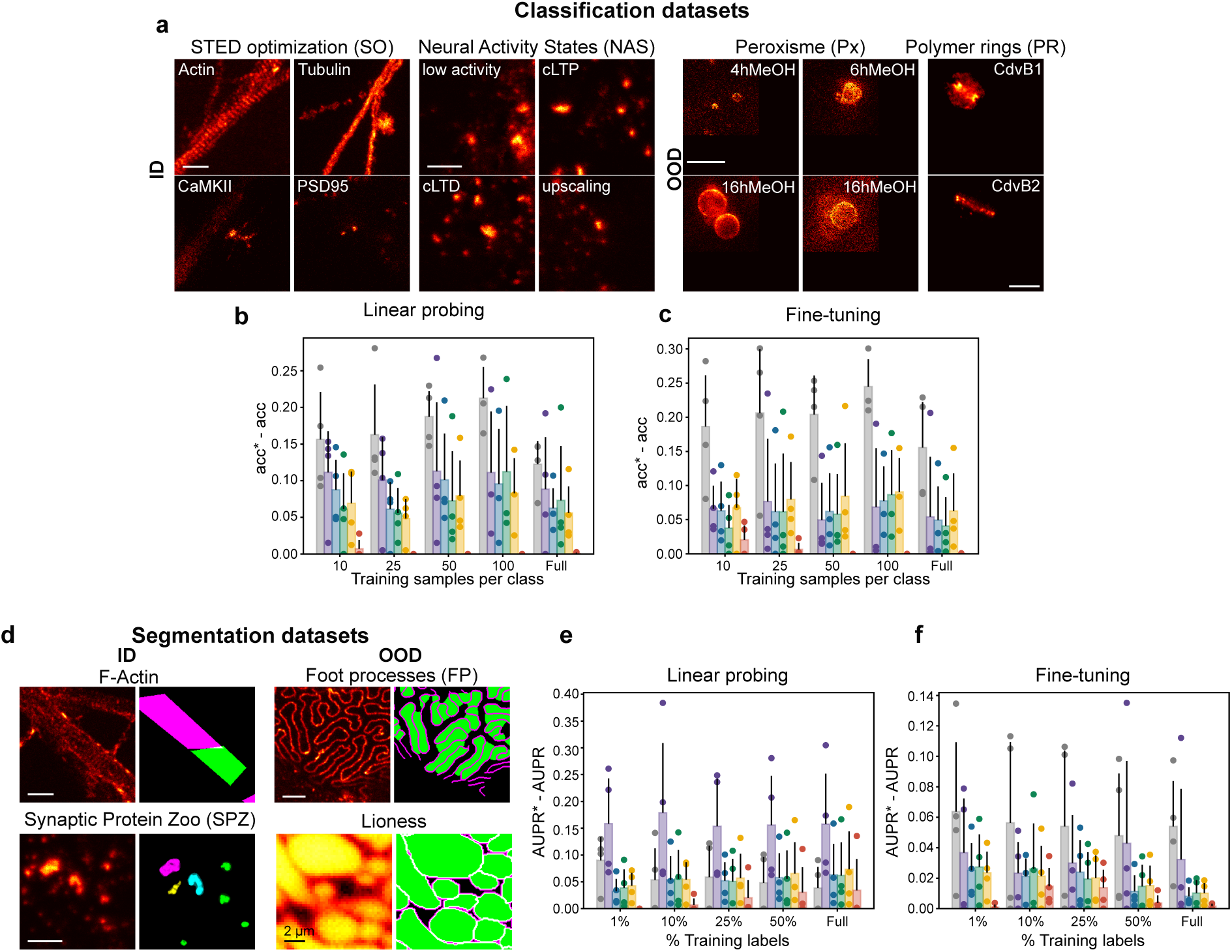
Supervised downstream tasks analysis. **a)** Example images and labels from 2 in-domain STED (ID-STED: *SO*, *NAS* ) and 2 out-of-domain STED (OOD-STED: *Px*, *PR*) datasets. **b)** Classification performance in linear probing in the small and full data regimes. Performance is reported as the difference from optimal accuracy (acc*^↑^ ↑* acc), where optimal accuracy represents the best performance achieved by any model for a given configuration. A model which performs best in all possible combinations of downstream dataset and number of samples per class would have acc*^↑^ ↑* acc = 0. **c)** Classification performance in end-to-end fine-tuning. **d)** Example images and segmentation ground truths from 2 in-domain STED (ID-STED: *F-Actin*, *SPZ* ) and 2 out-of-domain STED (OOD-STED: *FP*, *Lioness*) datasets. **e)** Segmentation performance in linear probing in the small and full data regimes. Performance is reported as the difference from optimal area under the precision recall curve (AUPR*^↑^ ↑* AUPR). **f)** Segmentation performance in end-to-end fine-tuning. Data points represent the average of five independent runs (different random seeds).

**Extended Data Fig. 2:**
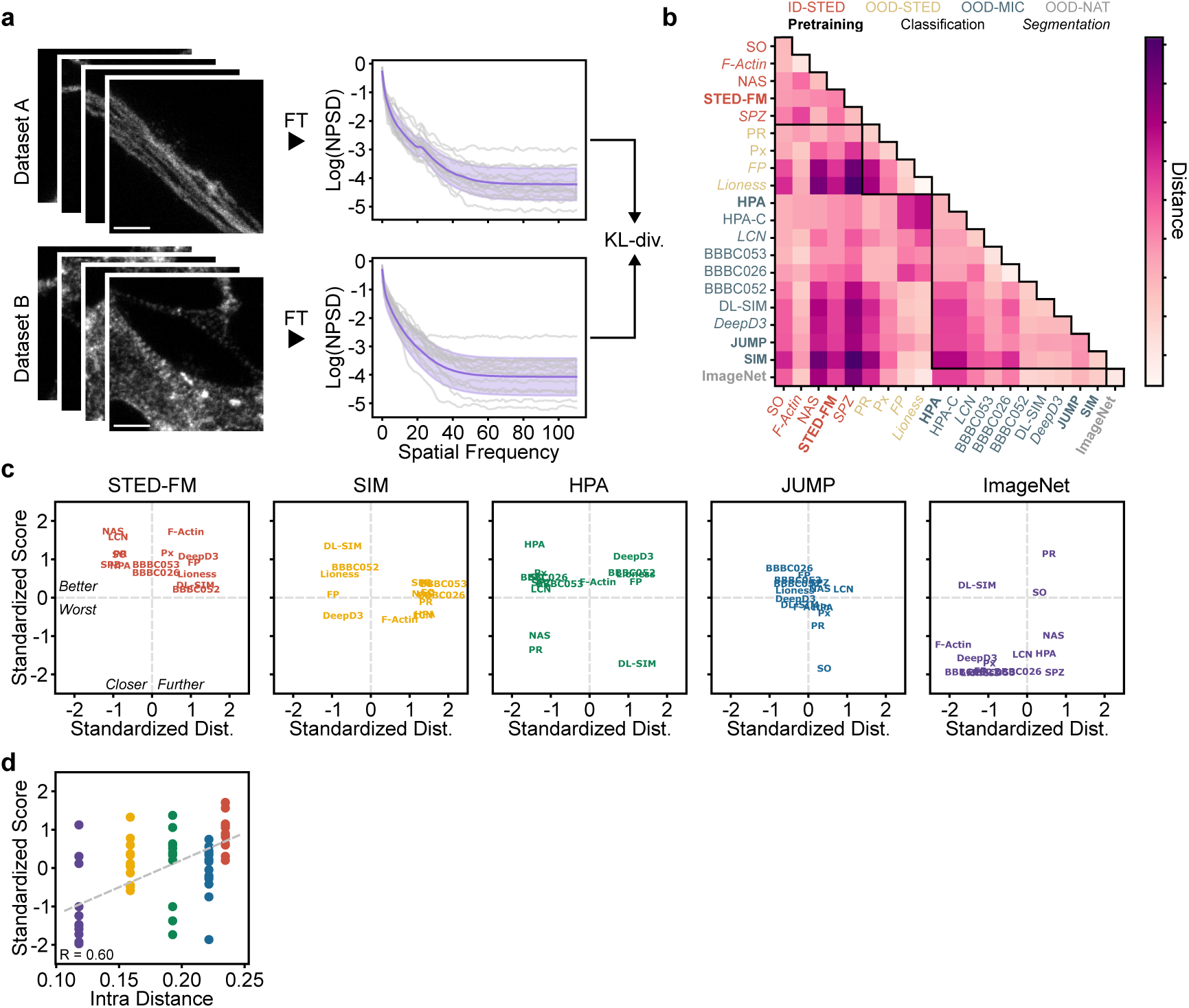
Correlation between domain distance and model performance on downstream classification and segmentation tasks. **a)** Schematic of the normalized power spectrum density distance. The normalized power spectrum density (NPSD) of images in both datasets is calculated. The Kullback Leibler divergence (KL-div) computes the distance between images. **b)** Measured distances between all datasets used in the manuscript. The intra-dataset distance corresponds to the diagonal. Colors are used to represent the ID-STED, OOD-STED, OOD-MIC, and OOD-NAT. Pretraining datasets are bold, classification datasets are in normal font, and segmentation datasets are in italic. **c)** For each classification and segmentation downstream tasks both the distance values and the performance metrics (accuracy for classification (F1-score for HPA classification) and AUPR for segmentation) were Z-score standardized within each specific task. STED pretraining is consistently over the average performance achieved on a given dataset even in cases where the distance of the pretraining dataset is further than other pretraining datasets (distance *>* 0 and score *>* 0). **d)** Pearson correlation coefficient between the dataset diversity (Intra Distance) and the average downstream performance across all tasks is computed (accuracy for classification (F1-score for HPA classification) and AUPR for segmentation). The performance within each downstream dataset was Z-score standardized.

**Extended Data Fig. 3:**
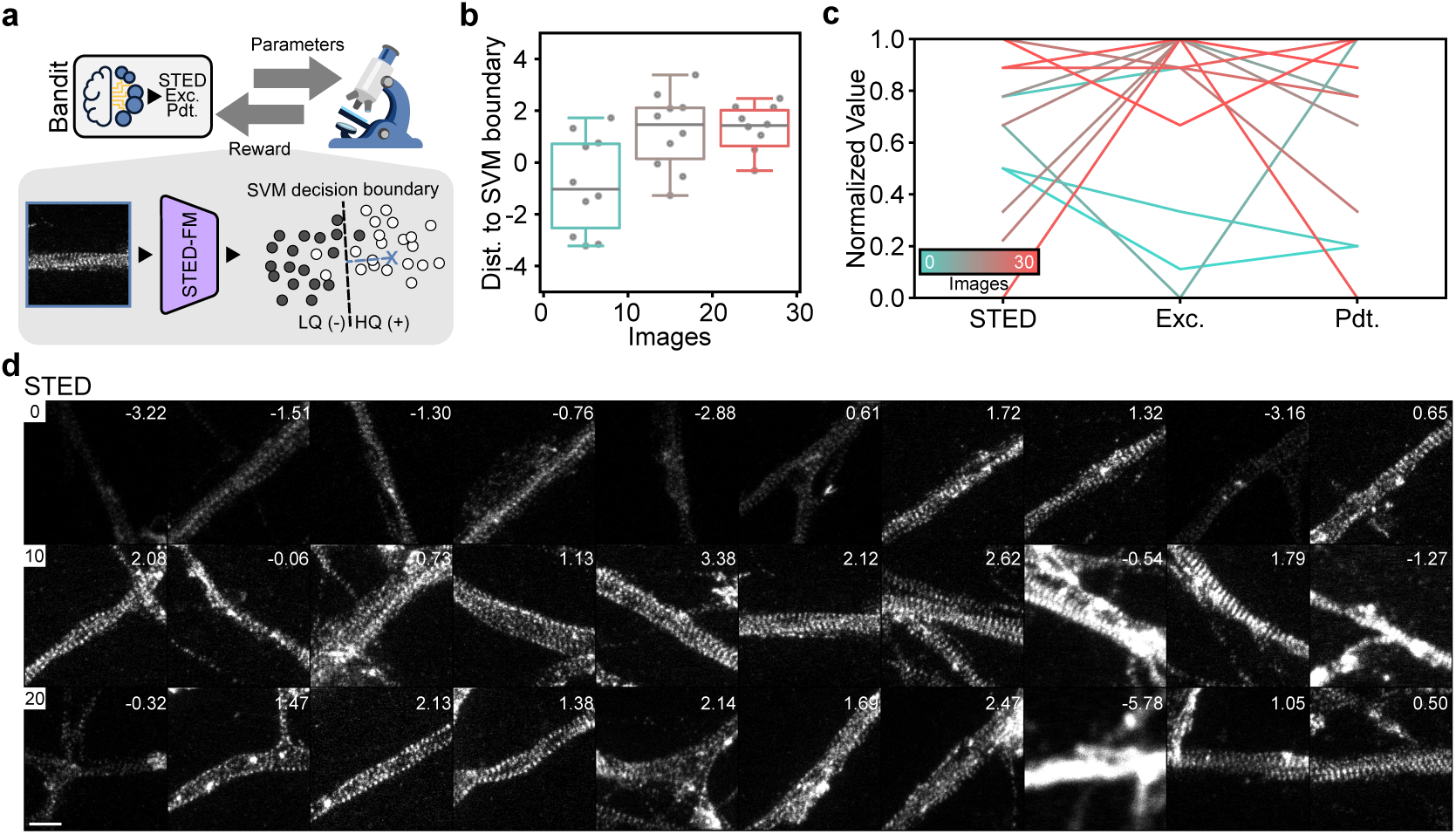
Bandit optimization using STED-FM as a quality metric. a) Schematic of the Bandit optimization routine. The Bandit algorithm selects imaging parameters and the reward is computed. The reward corresponds to the signed distance to the SVM decision boundary (Methods). b) Evolution of the distance to the SVM decision boundary. Boxplots in bins of 10 images. c) Evolution of the imaging parameters (STED power, excitation power, and pixel dwell time). Supplementary Tab. 17 presents the non-normalized imaging parameters. Boxplots present the the sample median, the first and third quantiles. The whiskers are 1.5 times the interquartile range. d) All images acquired with the Bandit model for the optimization of F-Actin STAR635. The measured quality is shown for each STED image in the top right corner. Scale bar: 1 μm.

