## Supplementary Materials for "A Self-Supervised Foundation Model for Robust and Generalizable Representation Learning in STED Microscopy"

### 6 Supplementary Material

### Contents

Tables . . . 3
Figures . . . 7

### Tables

| Dataset | Domain | # independent images | # crops |
|---|---|---|---|
| ImageNet [1] | Natural images | 1.3M | 1.3M |
| JUMP-CP [2] | Microscopy | 143 925 | 3 701 518 |
| HPA [3] | Microscopy | 124 288 | 1 118 592 |
| SIM [4] | Super-resolution microscopy | 3 656 | 205 946 |
| STED [5] | Super-resolution microscopy | 37 387 | 976 022 |

Supplementary Table 1: pretraining datasets overview.

| Feature | min | max |
|---|---|---|
| Area | 0.03 µm | 0.08 µm |
| Blur effect | 0.19 | 0.43 |
| Density | 1.41 proteins/µm | 4.83 proteins/µm |
| Eccentricity | 0.50 | 0.73 |
| Intensity | 0.03 | 0.17 |
| NN distance | 13.4 nm | 2.06 µm |
| # proteins | 3.74 | 32.97 |
| Entropy | 1.36 | 4.90 |
| SNR | 0.52 | 6.54 |

Supplementary Table 2: Values used to do min-max normalization in the clustering graphs of Supplementary Fig. 5. The min and max values correspond to quantiles 0.05 and 0.95, respectively.

| Dataset | Modality | # classes | # training instances |
|---|---|---|---|
| Optim [6] | STED | 4 | 1 536 |
| Neural activity states [7] | STED | 4 | 2 048 |
| Peroxisome [8] | STED | 4 | 352 |
| Polymer rings [9] | STED | 2 | 361 |
| DL-SIM [10] | SIM | 4 | 5 863 |
| BBBC026 [11] | Widefield | 2 | 6 120 |
| BBBC052 [12] | Confocal | 4 | 7 262 |
| BBBC053 [12] | Confocal | 2 | 1 809 |
| HPA-Classification [13] | Confocal | 28 | 24 857 (large images) |

Supplementary Table 3: Downstream classification datasets overview.

| Dataset | Modality | # classes | # training instances |
|---|---|---|---|
| F-actin [14] | STED | 2 | 941 |
| Synapses (semantic) [15] | STED | 4 | 1 191 |
| Foot processes [16] | STED | 2 | 3 049 |
| Lioness [17] | STED | 2 | 3 100 |
| LCN [18] | Confocal | 3 | 4 032 |
| DeepD3 [19] | Confocal and 2-photon | 2 | 1 017 |

Supplementary Table 4: Downstream segmentation datasets overview.

| | F-actin | SPZ | Foot processes | Lioness | LCN | DeepD3 |
|---|---|---|---|---|---|---|
| Random | N/A - 0.54 | N/A - 0.53 | N/A - 0.52 | N/A - 0.70 | N/A - 0.88 | N/A - 0.46 |
| ImageNet | 0.46 - **0.59** | 0.44 - **0.53** | 0.31 - **0.60** | 0.68 - **0.74** | 0.72 - **0.89** | 0.41 - 0.48 |
| JUMP | 0.51 - 0.57 | 0.55 - 0.57 | 0.41 - **0.59** | 0.72 - **0.74** | 0.73 - **0.90** | 0.45 - 0.48 |
| HPA | 0.51 - 0.56 | 0.54 - **0.56** | 0.41 - **0.59** | 0.72 - **0.74** | 0.74 - **0.90** | 0.47 - 0.48 |
| SIM | 0.49 - 0.56 | 0.55 - **0.56** | 0.40 - **0.59** | 0.72 - **0.74** | 0.72 - **0.90** | 0.44 - 0.48 |
| STED | **0.60 - 0.58** | **0.58 - 0.58** | **0.44 - 0.60** | **0.73 - 0.74** | **0.79 - 0.90** | **0.50 - 0.54** |

Supplementary Table 5: Evaluation of the segmentation performance of models pre-trained on different datasets (left column) on the downstream dataset using the F1-score as the evaluation metric. The average performance (N=5) is reported for both linear probing (left) and fine-tuning (right). The best performing models are in bold. For each training subset a Kruskal-Wallis test is performed followed by a post-hoc Mann-Whitney U rank test (two-tail). The complete statistical analysis is provided in Supplementary Data.

| | F-actin | SPZ | Foot processes | Lioness | LCN | DeepD3 |
|---|---|---|---|---|---|---|
| Random | N/A - 0.40 | N/A - 0.39 | N/A - 0.38 | N/A - 0.60 | N/A - 0.82 | N/A - 0.33 |
| ImageNet | 0.32 - **0.45** | 0.32 - **0.38** | 0.19 - **0.46** | 0.58 - **0.64** | 0.60 - **0.83** | 0.29 - 0.35 |
| JUMP | 0.38 - 0.43 | 0.42 - 0.43 | 0.28 - **0.46** | 0.62 - **0.64** | 0.60 - **0.84** | 0.32 - 0.34 |
| HPA | 0.38 - 0.43 | 0.40 - **0.42** | 0.27 - **0.46** | 0.62 - **0.64** | 0.61 - **0.84** | 0.34 - 0.35 |
| SIM | 0.36 - 0.43 | 0.41 - **0.42** | 0.26 - **0.46** | 0.62 - **0.64** | 0.59 - **0.84** | 0.31 - 0.35 |
| STED | **0.46 - 0.45** | **0.44 - 0.44** | **0.30 - 0.46** | **0.63 - 0.64** | **0.68 - 0.84** | **0.36 - 0.40** |

Supplementary Table 6: Evaluation of the segmentation performance of models pre-trained on different datasets (left column) on the downstream dataset using the Intersection over Union (IoU) as the evaluation metric. The average performance (N=5) is reported for both linear probing (left) and fine-tuning (right). The best performing models are in bold. For each training subset a Kruskal-Wallis test is performed followed by a post-hoc Mann-Whitney U rank test (two-tail). The complete statistical analysis is provided in Supplementary Data.

| Dataset | Modality | # training instances |
|---|---|---|
| $\alpha$-tubulin | STED | 1 145 |
| $\beta$-tubulin | STED | 1 200 |
| VGAT-LQHQ | STED | 881 |
| Gephyrin | STED | 881 |

Supplementary Table 7: Downstream denoising datasets overview.

| Dataset | Modality | # training instances |
|---|---|---|
| VGAT-SR | Confocal to STED | 881 |
| F-actin-SR | Confocal to STED | 4 331 |

Supplementary Table 8: Downstream super-resolution datasets overview.

| Metric | MSE | | | PSNR | | | SSIM | | |
|---|---|---|---|---|---|---|---|---|---|
| Model | pixpix | DDIM | DRaFT | pixpix | DDIM | DRaFT | pixpix | DDIM | DRaFT |
| pix2pix | - | $1.058 \times 10^{-7}$ | $0.5750 \times 10^{-7}$ | - | $1.058 \times 10^{-7}$ | $0.5750 \times 10^{-7}$ | - | 0.0015 | 0.0006 |
| DDIM | $1.058 \times 10^{-7}$ | - | 0.8476 | $1.058 \times 10^{-7}$ | - | 0.8476 | 0.0015 | - | 0.8333 |
| DRaFT | $0.5750 \times 10^{-7}$ | 0.8476 | - | $0.5750 \times 10^{-7}$ | 0.8476 | - | 0.0006 | 0.8333 | - |

Supplementary Table 9: P-values from Mann-Whitney U-tests (two-tailed) comparing the distribution of pixel-level metrics obtained from the different generative models on the *F-actin-SR* dataset (N=26 images). A Kruskal-Wallis test is performed before post-hoc comparison: $p$-value$_{\text{MSE}}$: $3.655 \times 10^{-9}$; $p$-value$_{\text{PSNR}}$: $3.655 \times 10^{-9}$; $p$-value$_{\text{SSIM}}$: 0.0006.

| Metric | Dice (rings) | | | Dice (fibers) | | |
|---|---|---|---|---|---|---|
| Model | pixpix | DDIM | DRaFT | pixpix | DDIM | DRaFT |
| pix2pix | - | 0.0161 | 0.0070 | - | 0.4418 | 0.3833 |
| DDIM | 0.0161 | - | 0.9339 | 0.4418 | - | 0.8777 |
| DRaFT | 0.0070 | 0.9339 | - | 0.3833 | 0.8777 | - |

Supplementary Table 10: P-values from Mann-Whitney U-tests (two-tailed) comparing the distribution of segmentation metrics obtained from the different generative models on the *F-actin-SR* dataset (N=26 images). A Kruskal-Wallis test is performed before post-hoc comparison: $p$-value$_{\text{Rings}}$: 0.011; $p$-value$_{\text{Fibers}}$: 0.614.

| Metric | Expert choice | | |
|---|---|---|---|
| Model | pixpix | DDIM | DRaFT |
| pix2pix | - | 0.0228 | 0.0228 |
| DDIM | 0.0228 | - | 0.0228 |
| DRaFT | 0.0228 | 0.0228 | - |

Supplementary Table 11: P-values from Mann-Whitney U-tests (two-tailed) comparing the distribution of expert (N=4) choices for the user of the algorithmic super-resolution *F-actin-SR* experiment (N=26 images). A Kruskal-Wallis test is performed before post-hoc comparison: $p$-value: 0.0059.

| Feature | Real data % | | Synthetic data % | |
|---|---|---|---|---|
| Age | DIV12 | DIV25 | sDIV12 | sDIV25 |
| Area | 4.71 | 3.70 | 1.08 | 11.83 |
| Density | 4.71 | 4.44 | 1.08 | 0.54 |
| Eccentricity | 7.06 | 2.96 | 9.14 | 5.91 |
| Intensity | 5.88 | 3.70 | 4.84 | 6.45 |
| Nanodomains | 5.88 | 3.70 | 2.15 | 12.90 |
| Perimeter | 3.53 | 5.19 | 0.54 | 11.29 |
| Solidity | 3.53 | 5.19 | 0.00 | 12.37 |
| Total | 4.48 | | 5.72 | |

Supplementary Table 12: Percentage of images whose features are significantly different ($p < 0.05$) from their corresponding target feature distributions. For example, for real DIV12 and synthetic DIV12 data, the target distribution is the real DIV12 data. The p-value of a data point is obtained from computing its z-score in the target distribution.

| Structure | # crops |
|---|---|
| Adducin | 1 711 |
| Bassoon | 44 649 |
| $\beta$-CaMKII | 6 481 |
| $\beta_2$-spectrin | 966 |
| CaMKII | 355 |
| F-actin | 34 547 |
| FUS | 20 820 |
| Gephyrin | 272 |
| Glur1 | 100 |
| Homer | 16 394 |
| Lifeact | 284 |
| Live tubulin | 124 |
| Map2 | 242 |
| NKCC2 | 1 381 |
| PSD95 | 58 074 |
| Rim | 8 453 |
| Sir-Actin | 2 137 |
| Sir-Tubulin | 202 |
| Tom20 | 35 400 |
| Tubulin | 312 |
| VGAT | 748 |
| VGLUT1 | 1 086 |
| VGLUT2 | 3 675 |
| Vimentin | 270 |

Supplementary Table 13: Number of crops which contain the different structures in the labeled subset of the *STED-FM dataset*.

| | Species | Supplier | Category | Dilution | Related Fig. |
|---|---|---|---|---|---|
| Bassoon | Rabbit | Synaptic System | 141 003 | 1:500 | Fig. 6 |
| PSD95 | Mouse | Invitrogen | MA1-045 | 1:500 | Fig. 6c |

Supplementary Table 14: Primary antibodies

| Fluorophores | Host | Reactivity | Supplier | Category | Dilution | Related Fig. |
|---|---|---|---|---|---|---|
| ATTO490LS | Goat | Rabbit | Hypermol | 2309-1MG | 1:250 | Fig. 6c |
| AF594 | Goat | Rabbit | Invitrogen | A32740 | 1:100 | Fig. 6a-b |
| AF594 | Goat | Mouse | Thermofisher | A32742 | 1:100 | Fig. 6c |

Supplementary Table 15: Secondary antibodies

| Imaging parameter | Optimization range |
|---|---|
| STED power | [0, 325] mW |
| Excitation power | [2.5, 6.6] µW |
| Pixel dwell time | [8, 20] µs |

Supplementary Table 16: Bandit optimization imaging parameters.

| Imaging parameter | Optimization range |
|---|---|
| STED power | [0, 210] mW |
| Excitation power | [0.9, 10.2] µW |
| Pixel dwell time | [5, 30] µs |

Supplementary Table 17: Bandit optimization imaging parameters with the STED-FM as quality metric.

| Parameter | F-actin STAR635 | PSD95 AF594 | Bassoon ATTO490LS |
|---|---|---|---|
| STED power | [0, 337.5] mW | [0, 337.5] mW | [0, 337.5] mW |
| Excitation power | [0.0, 4.1] µW | [0.0, 3.8] µW | [0.0, 14.9] µW |
| Pixel dwell time | [1, 100] µs | [1, 100] µs | [1, 100] µs |

Supplementary Table 18: Range of imaging parameters for the RL experiments.

### 11 Figures

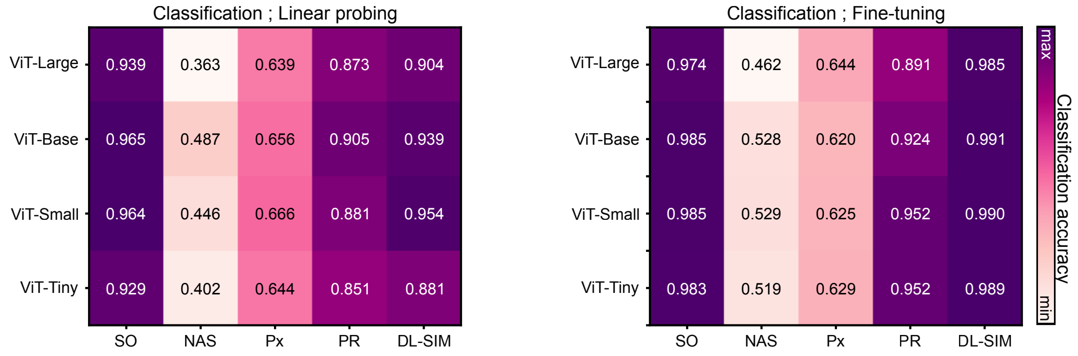


Supplementary Fig. 1: Classification accuracy as a function of model scale on the different downstream datasets. Left: linear probing. Right: end-to-end fine-tuning.

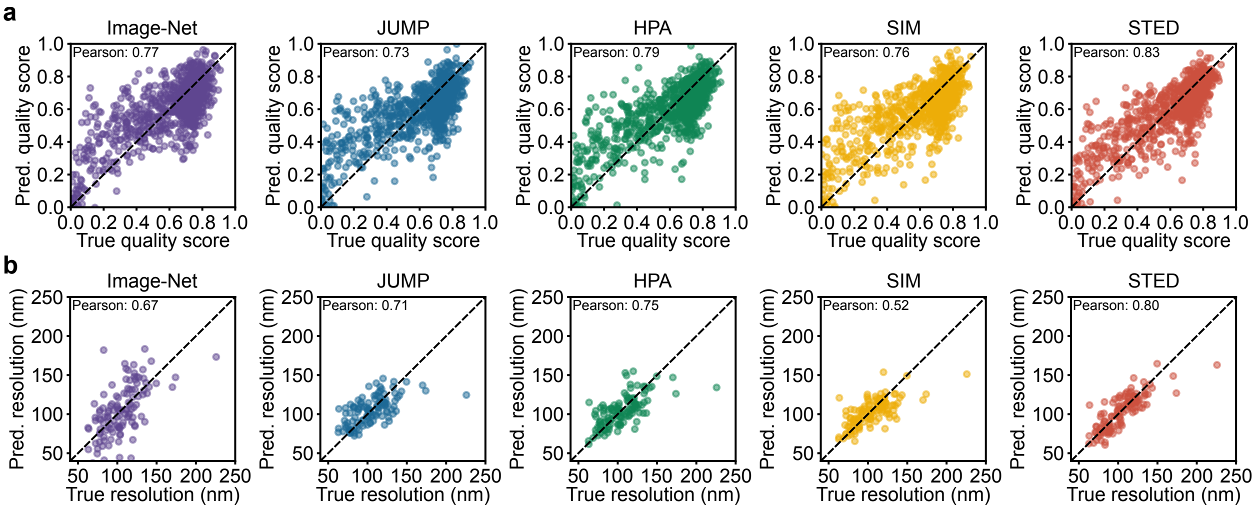


Supplementary Fig. 2: Predicted image attributes such as a) the image quality and b) the spatial resolution from the latent representation of all pretrained models. Pearson correlation between the ground truth and predicted values is shown on the graph for comparison.

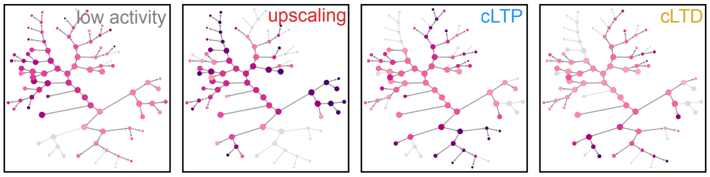


Supplementary Fig. 3: Graphs obtained from performing recursive clustering on the handcrafted feature space. Node sizes are given by the number of embeddings in the node. Nodes are color-coded by the proportion of the given class. Darker nodes are associated with a higher proportion.

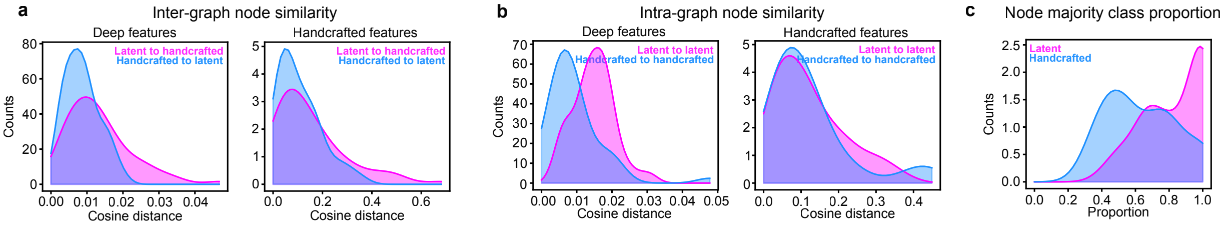


Supplementary Fig. 4: a) Distribution of the minimum cosine distance between nodes from two different graphs. For each node in one graph, this represents the smallest cosine distance to any node in the other graph. The cosine distance is computed using either features extracted by STED-FM (deep features, left) or manual feature extraction(handcrafted features, right). The feature vector of a node corresponds to the average feature vector of all embeddings in that node. Larger distances are associated with novel subtypes. b) Distribution of the minimum cosine distance between nodes from the same graph. The cosine distance is computed using either features extract by STED-FM (deep features, left) or manual feature extraction (handcrafted features, right). Larger distances are associated with more dissimilar nodes. c) Distribution of majority class proportion in the nodes (node purity). A perfect clustering would have a node purity of 1.0.

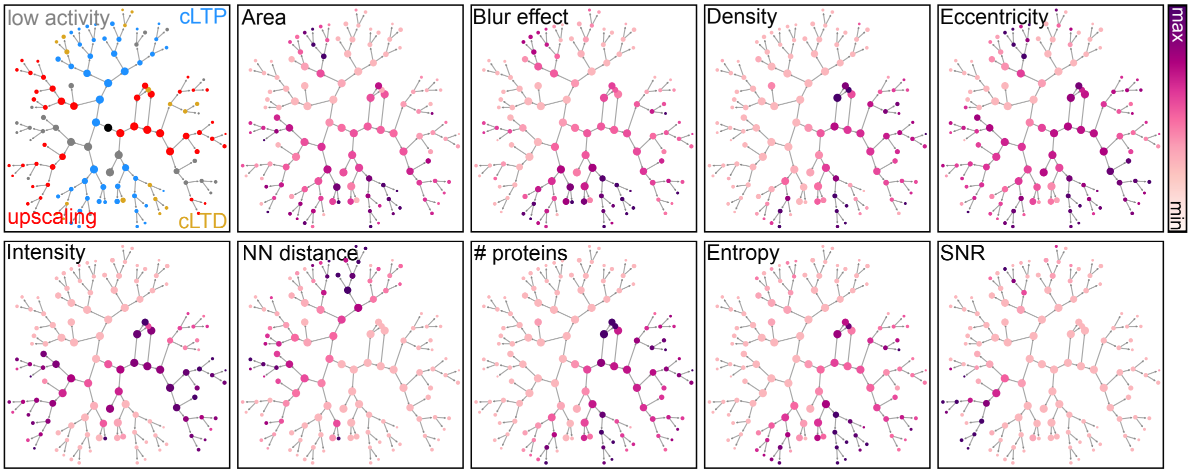


Supplementary Fig. 5: Recursive clustering graphs color-coded by feature value (except the top-left graph which is color-coded by majority class proportion). The min/max values are given in the Supplementary Table 2.

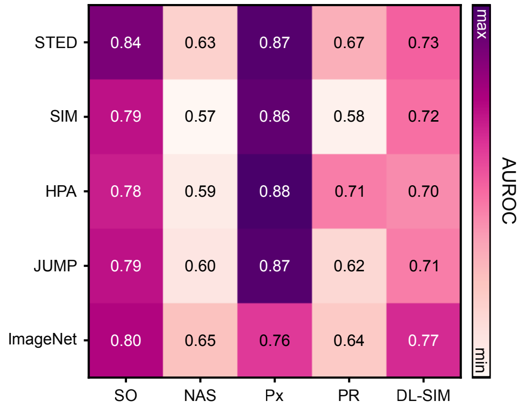


Supplementary Fig. 6: Image retrieval performance on the different downstream datasets for the pretraining datasets. The performance is reported in terms of AUROC.

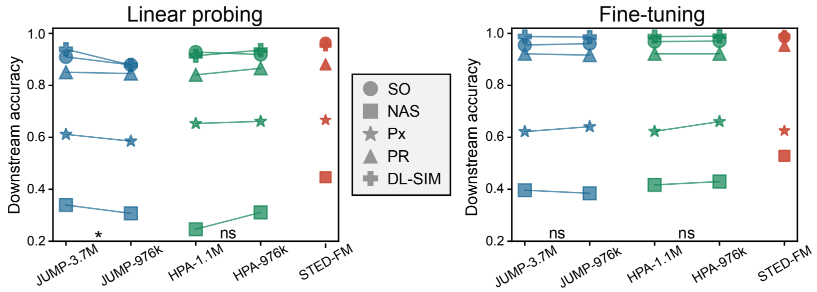


Supplementary Fig. 7: Downstream classification performance of the model on 5 different datasets for JUMP (blue) and HPA (green) pre-training after matching the dataset size to the STED pretraining dataset (red, 976 022 crops). Left: linear probing. Right: end-to-end fine-tuning. Statistical significance was measured using a t-test with the null hypothesis that the mean average difference between the accuracy of the full upstream dataset and the restricted upstream dataset is zero (Linear Probing: JUMP $p$ = 0.026. HPA $p$ = 0.147. Fine-tuning: JUMP $p$ = 0.858. HPA $p$ = 0.198). Data points represent the average of five independent runs (different random seeds).

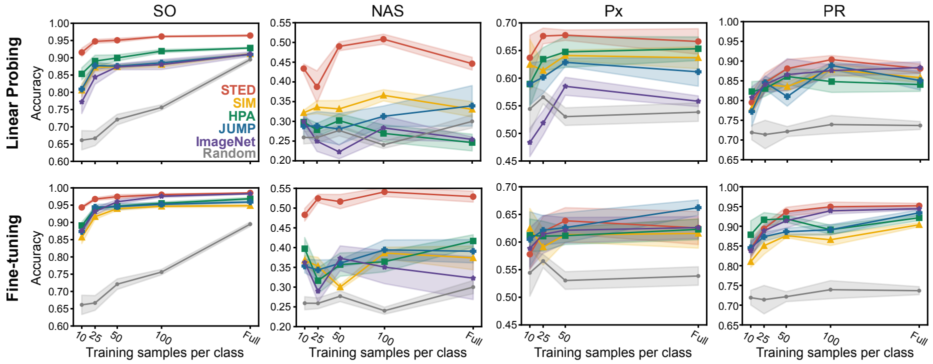


Supplementary Fig. 8: Classification accuracy for different training sample size measured for all downstream ID-STED and OOD-STED datasets. Top row: linear probing. Bottom row: end-to-end fine-tuning. *Random* refers to the random initialization of the weights of the models trained from scratch. Data points represent the average of five independent runs (different random seeds), with the shaded area representing the standard error of the mean (SEM). The complete statistical analysis is provided in Supplementary Data.

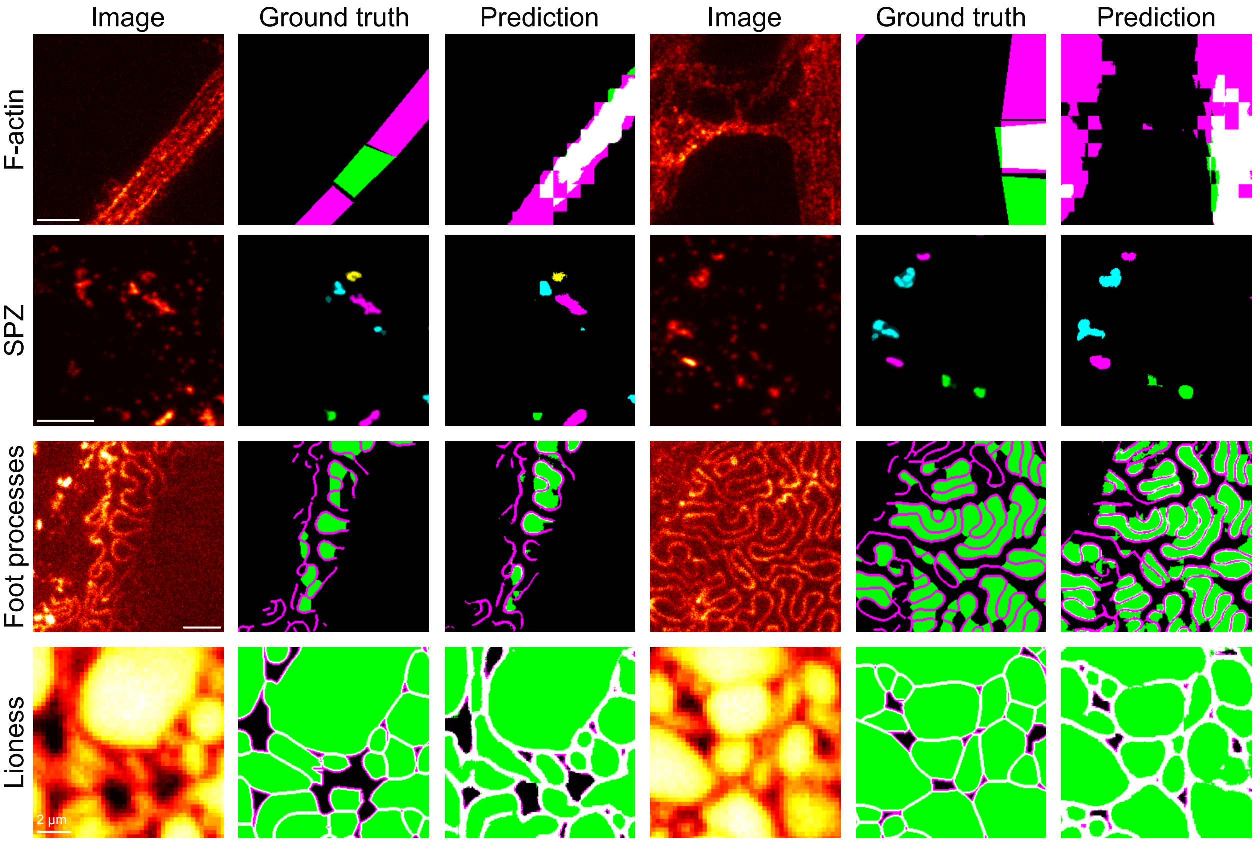


Supplementary Fig. 9: Example predictions of the segmentation masks from STED-FM fine-tuned on the downstream datasets. Scale bar: 1 µm unless otherwise noted.

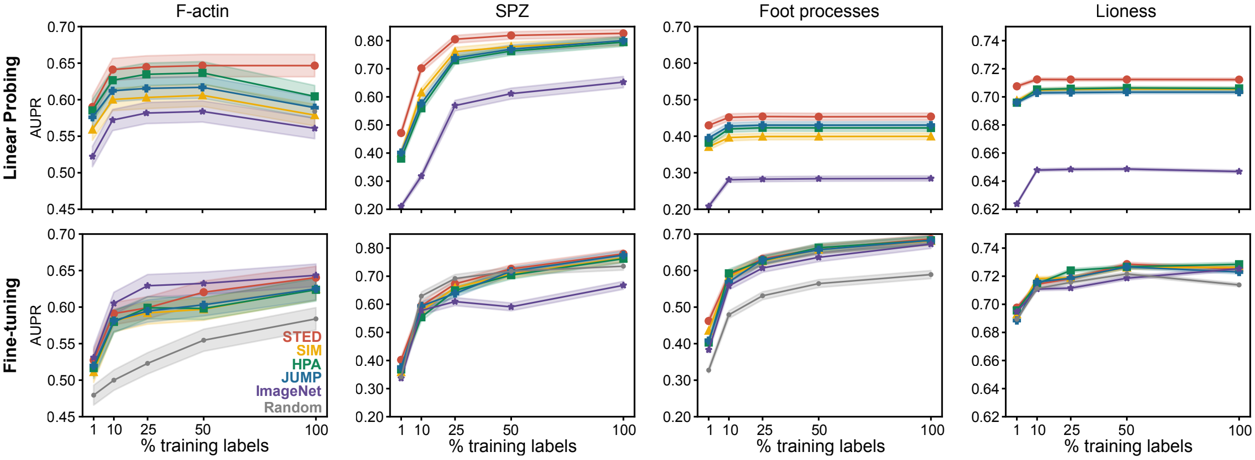


Supplementary Fig. 10: Segmentation AUPR for different training sample size measured for all downstream datasets. Top row: linear probing. Bottom row: end-to-end fine-tuning. The ground truths for the *SPZ* dataset are weak labels, i.e., not all synaptic proteins in the images have been annotated. Data points represent the average of five independent runs (different random seeds), with the shaded area representing the standard error of the mean (SEM). The complete statistical analysis is provided in Supplementary Data.

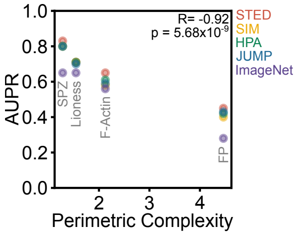


Supplementary Fig. 11: Effect of the perimeter complexity on the segmentation tasks. The segmentation performance of all pre-trained models decrease with increasing perimeter complexity.

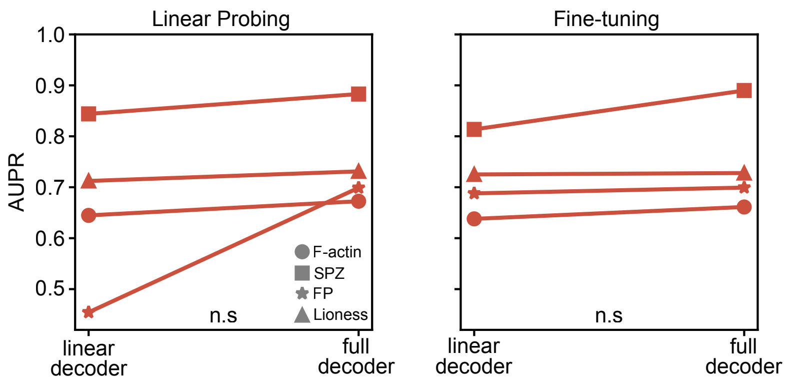


Supplementary Fig. 12: Segmentation performance improvement when using the full decoder compared to the lightweight linear decoder, in (left) linear probing and (right) end-to-end fine-tuning. The test for statistical significance was a t-test with null hypothesis that the mean average difference between the accuracy of the linear decoder and the full decoder is zero. Linear Probing: $p = 0.224$. Fine-tuning: $p = 0.183$. Data points represent the average of five independent runs (different random seeds).

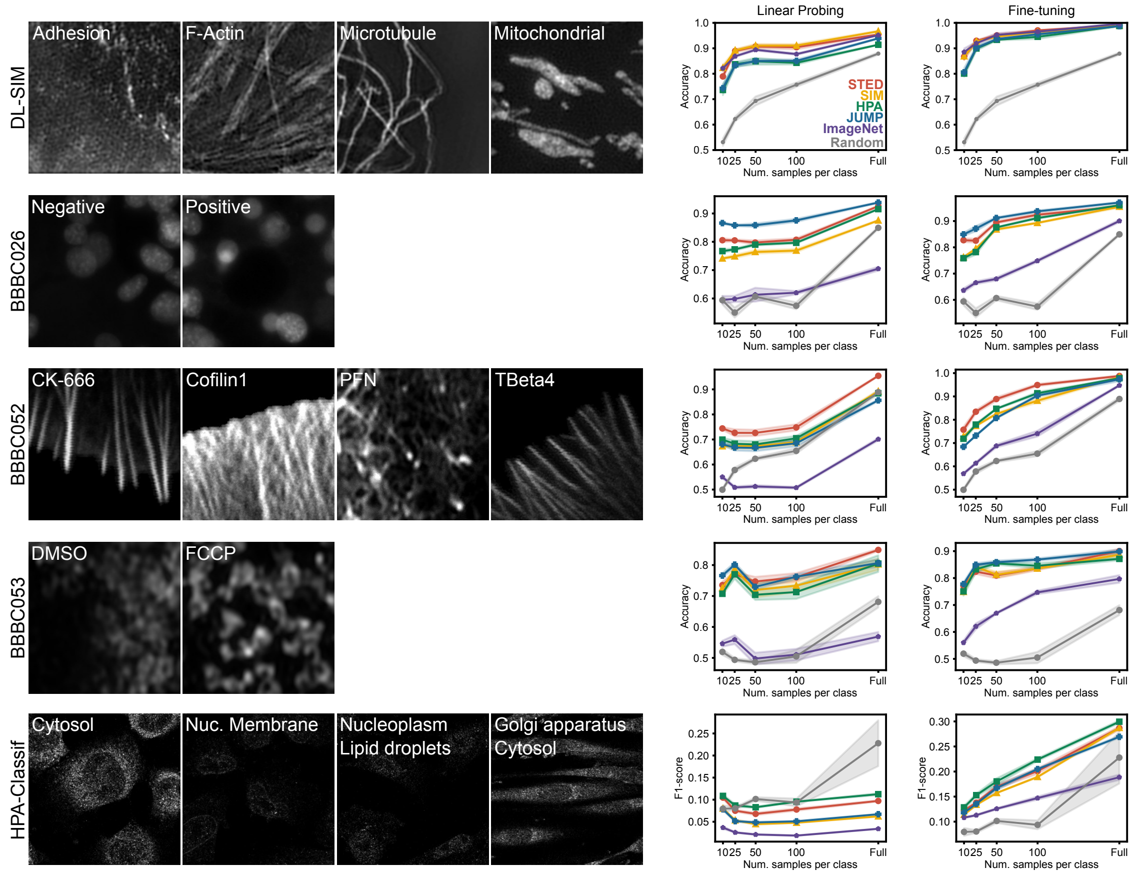


Supplementary Fig. 13: Classification accuracy for different training sample size measured for all downstream OOD-MIC datasets. Examples of training images are shown. For HPA-classification only a subset of classes are displayed. Left: Linear probing. Right: end-to-end fine-tuning. *Random* refers to the random initialization of the weights of models trained from scratch. Data points represent the average of five independent runs (different random seeds), with the shaded area representing the standard error of the mean (SEM). The complete statistical analysis is provided in Supplementary Data.

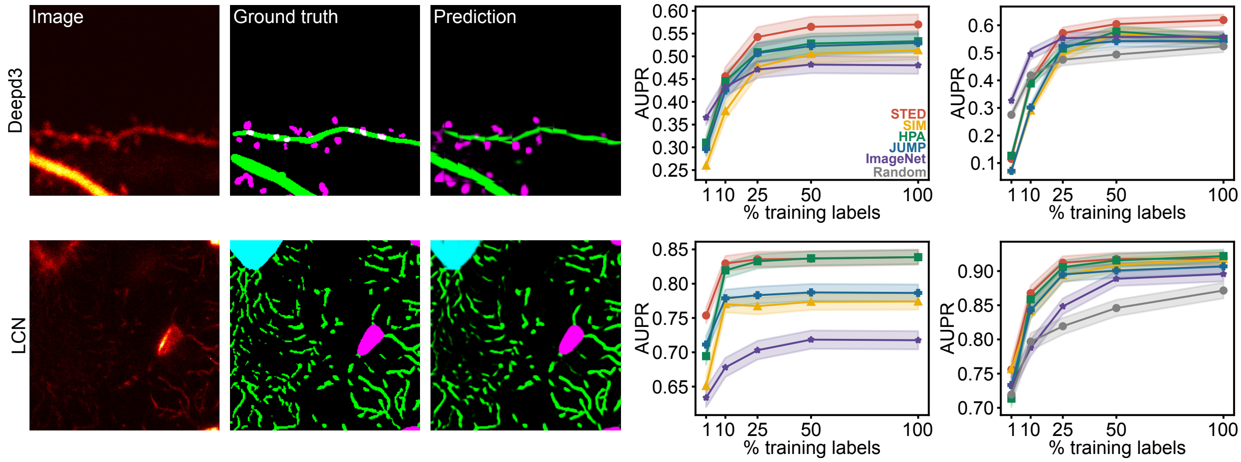


Supplementary Fig. 14: Segmentation task on the OOD-MIC datasets. Example images with their corresponding ground truth annotations and predictions from STED-FM fine-tuned on the downstream datasets. Segmentation AUPR for different training sample size measured for all downstream datasets. Left: Linear probing. Right: End-to-end fine-tuning. *Random* refers to the random initialization of the weights of models trained from scratch. Data points represent the average of five independent runs (different random seeds), with the shaded area representing the standard error of the mean (SEM). The complete statistical analysis is provided in Supplementary Data.

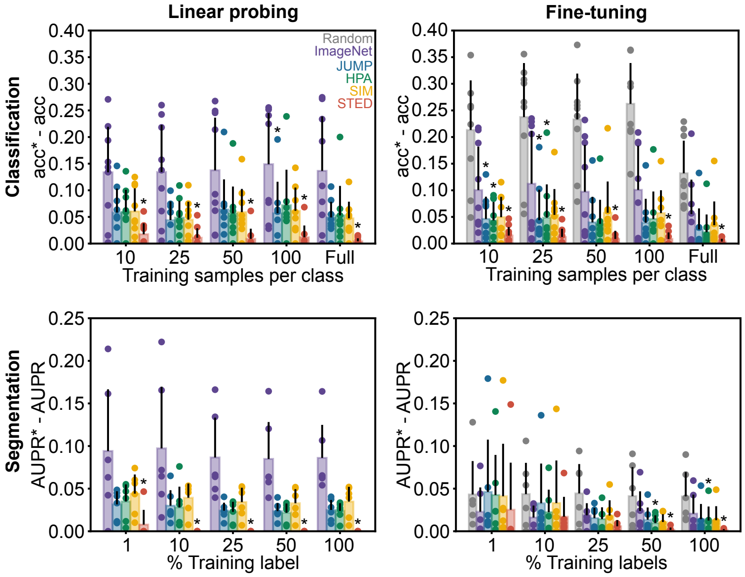


Supplementary Fig. 15: Performance on the classification and segmentation tasks for linear probing and fine-tuning. All datasets for classification (N=9) and segmentation (N=6) are combined. Performance is reported as the difference from optimal accuracy, where optimal accuracy represents the best performance achieved by any model for a given configuration. A model which performs best in all possible combinations of downstream dataset and number of training samples would have $\text{acc}^* - \text{acc} = 0$ (segmentation: $\text{AUPR}^* - \text{AUPR} = 0$). A $*$ is added on the best performing models. For each training subset a Kruskal-Wallis test is performed followed by a post-hoc Mann-Whitney U rank test (one-tailed). Data points represent the average of five independent runs (different random seeds). Bar plot presents the average of the distribution and the black line the standard deviation. The complete statistical analysis is provided in Supplementary Data.

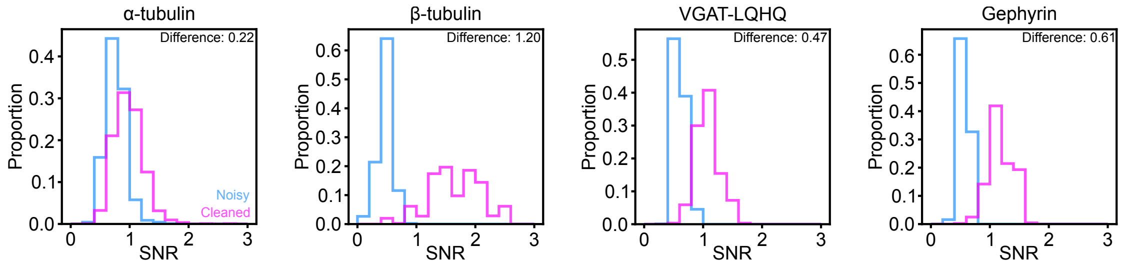


Supplementary Fig. 16: Quantification of the change in signal-to-noise ratio (SNR) across the training datasets used for the denoising task ($\alpha$-tubulin, $\beta$-tubulin, VGAT-LQHQ, Gephyrin). The difference in average SNR between high SNR (ground truth) and the low SNR input images is reported ($\mathrm{SNR}_{\mathrm{high}} - \mathrm{SNR}_{\mathrm{low}}$). A larger reported value indicates a greater disparity in signal quality between the input and target images, representing a more challenging denoising task.

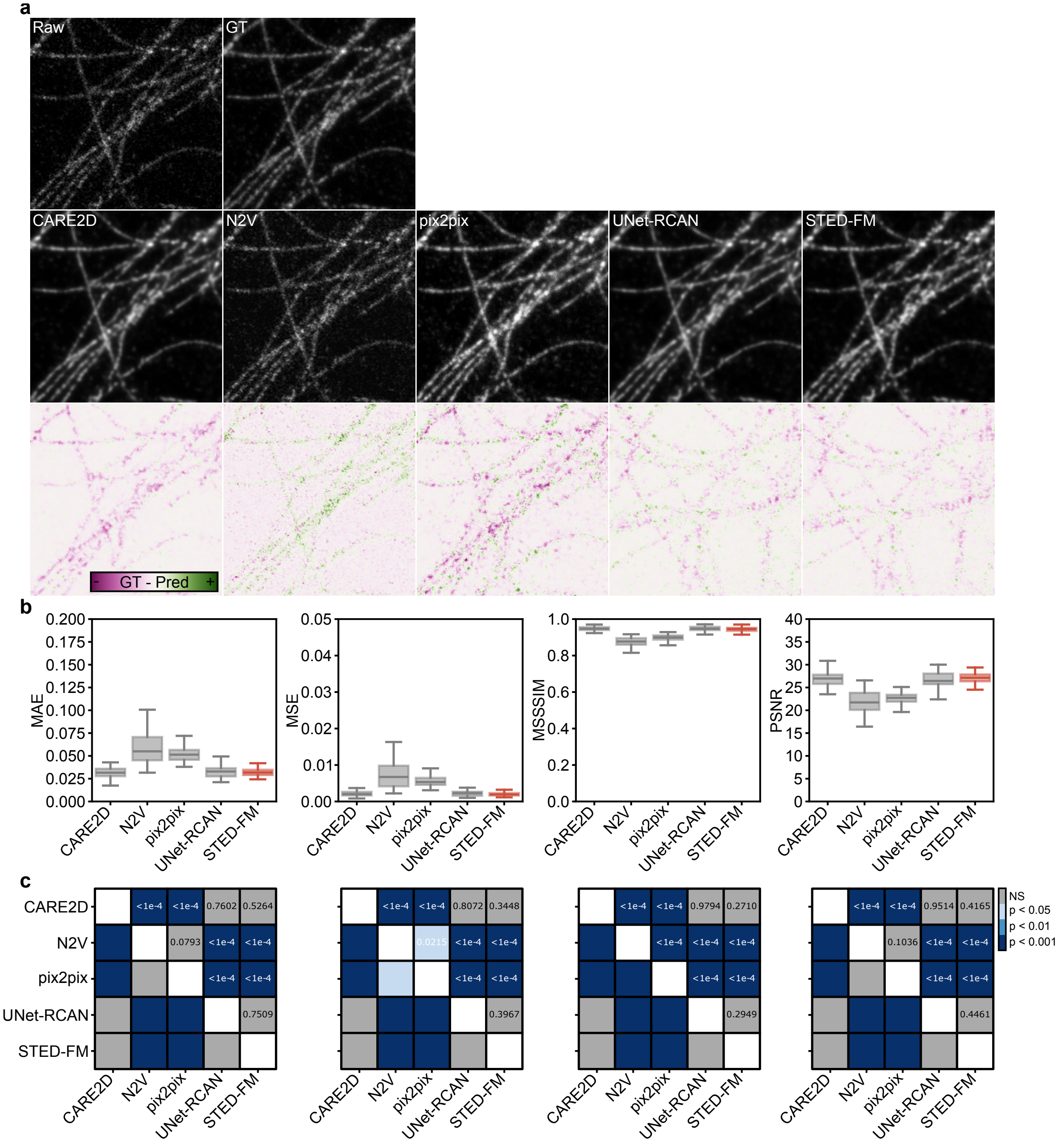


Supplementary Fig. 17: Denoising performance on the $\alpha$-tubulin (OOD-STED) dataset. a) Qualitative comparison and residual analysis. (Top row) Presents the raw, noisy input image and the corresponding ground truth image. (Middle row) Displays the denoised outputs generated by the baselines and the STED-FM model. (Bottom row) Shows the residual image, calculated by subtracting the denoised output from the ground truth. The residual map highlights errors: magenta indicates regions where the denoised signal is higher, while green indicates regions where the signal is lower. b) Evaluation of the denoising performance across all models using pixel-wise metrics: MAE (mean absolute error), MSE (mean squared error), MSSSIM (multiscale structural similarity), and PSNR (peak signal to noise ratio). Higher MSSSIM and PSNR indicate better performance, while lower MAE and MSE are favorable. Boxplots present the the sample median, the first and third quantiles. The whiskers are 1.5 times the interquartile range. (N=47) c) Statistical comparison of model performance. The matrix summarizes the results of the resampling test comparing the performance of each baseline (row) against every other baseline (column) for the metrics presented in (b). The matrix includes the raw $p$-values for the pair-wise comparisons, allowing for the assessment of the significance of performance differences.

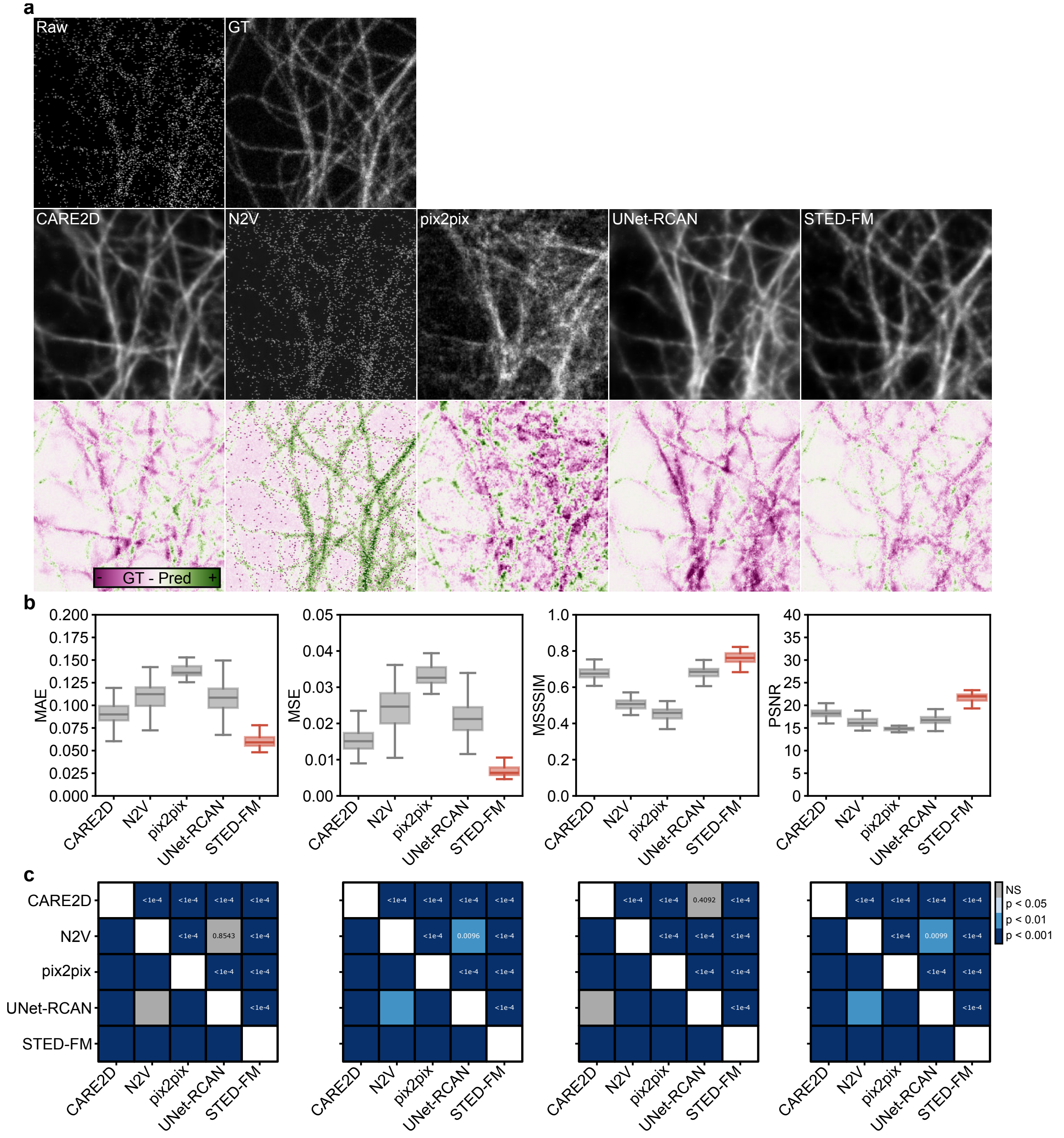


Supplementary Fig. 18: Denoising performance on the $\beta$-tubulin (OOD-STED) dataset. a) Qualitative comparison and residual analysis. (Top row) Presents the raw, noisy input image and the corresponding ground truth image. (Middle row) Displays the denoised outputs generated by the baselines and the STED-FM model. (Bottom row) Shows the residual image, calculated by subtracting the denoised output from the ground truth. The residual map highlights errors: magenta indicates regions where the denoised signal is higher, while green indicates regions where the signal is lower. b) Evaluation of the denoising performance across all models using pixel-wise metrics: MAE (mean absolute error), MSE (mean squared error), MSSSIM (multiscale structural similarity), and PSNR (peak signal to noise ratio). Higher MSSSIM and PSNR indicate better performance, while lower MAE and MSE are favorable. Boxplots present the the sample median, the first and third quantiles. The whiskers are 1.5 times the interquartile range. (N=100) c) Statistical comparison of model performance. The matrix summarizes the results of the resampling test comparing the performance of each baseline (row) against every other baseline (column) for the metrics presented in (b). The matrix includes the raw $p$-values for the pair-wise comparisons, allowing for the assessment of the significance of performance differences.

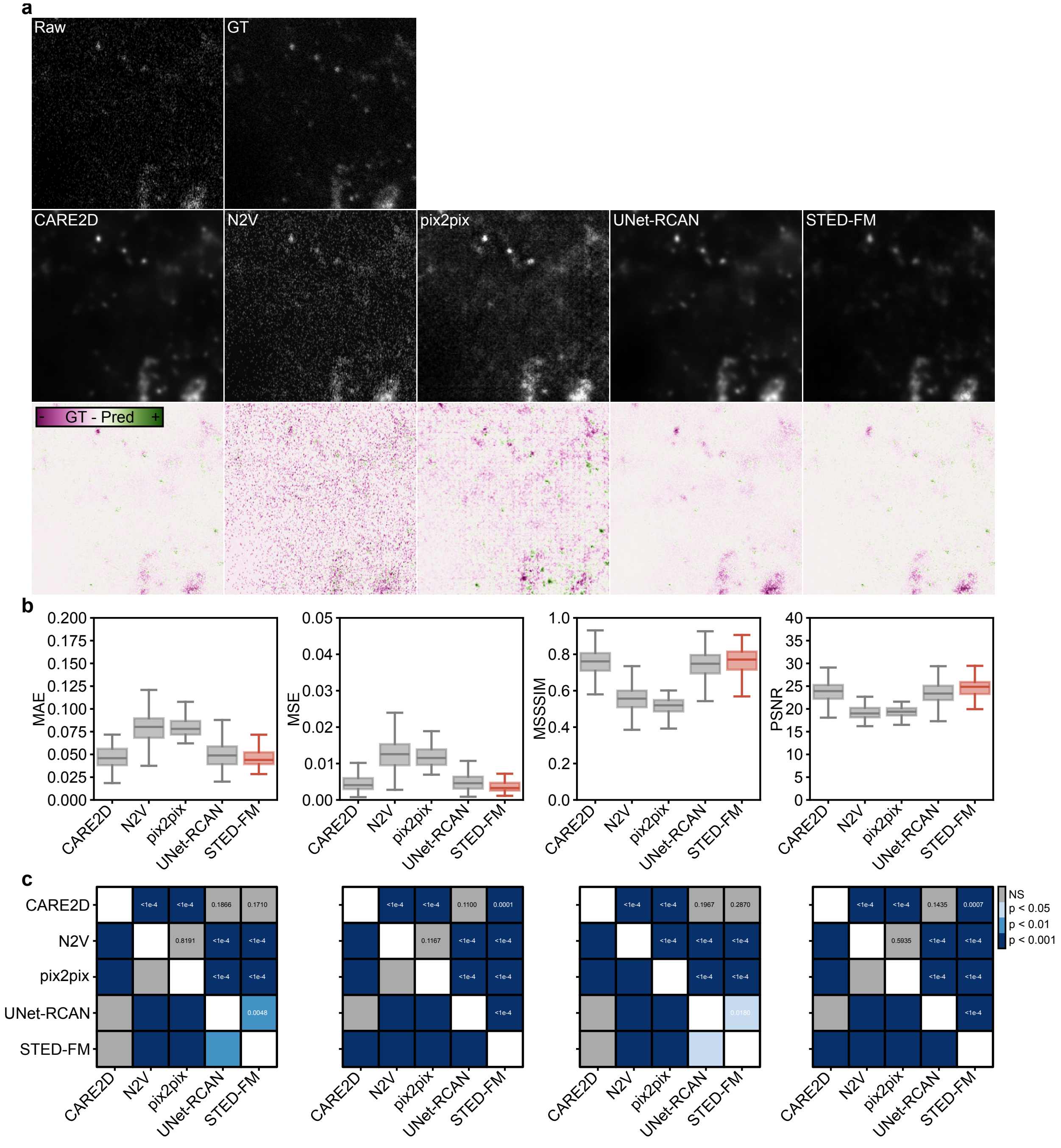


Supplementary Fig. 19: Denoising performance on the VGAT-LQHQ (OOD-STED) dataset. a) Qualitative comparison and residual analysis. (Top row) Presents the raw, noisy input image and the corresponding ground truth image. (Middle row) Displays the denoised outputs generated by the baselines and the STED-FM model. (Bottom row) Shows the residual image, calculated by subtracting the denoised output from the ground truth. The residual map highlights errors: magenta indicates regions where the denoised signal is higher, while green indicates regions where the signal is lower. b) Evaluation of the denoising performance across all models using pixel-wise metrics: MAE (mean absolute error), MSE (mean squared error), MSSSIM (multiscale structural similarity), and PSNR (peak signal to noise ratio). Higher MSSSIM and PSNR indicate better performance, while lower MAE and MSE are favorable. Boxplots present the the sample median, the first and third quantiles. The whiskers are 1.5 times the interquartile range. (N=187) c) Statistical comparison of model performance. The matrix summarizes the results of the resampling test comparing the performance of each baseline (row) against every other baseline (column) for the metrics presented in (b). The matrix includes the raw $p$-values for the pair-wise comparisons, allowing for the assessment of the significance of performance differences.

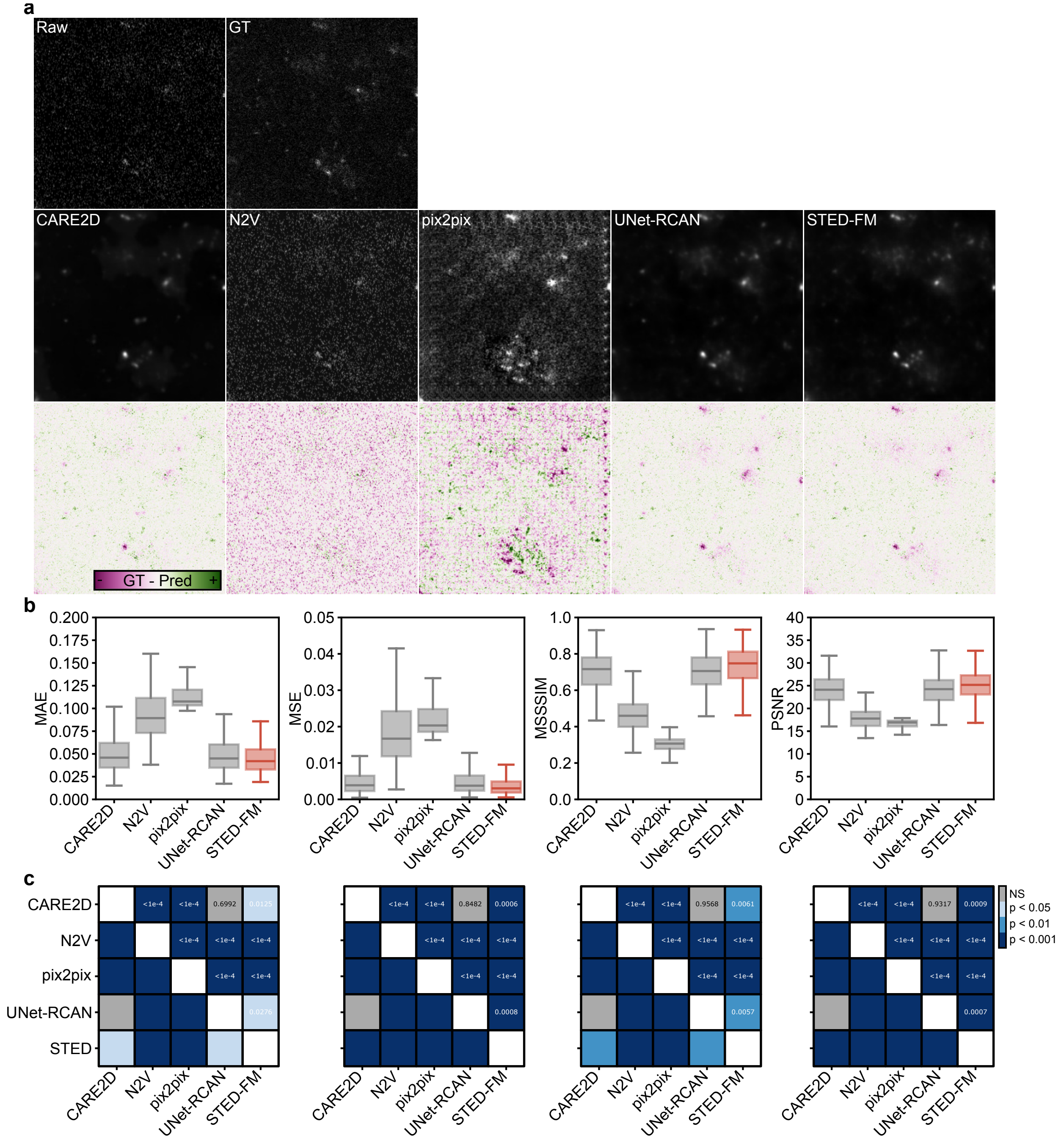


Supplementary Fig. 20: Denoising performance on the Gephyrin-LQHQ (OOD-STED) dataset. a) Qualitative comparison and residual analysis. (Top row) Presents the raw, noisy input image and the corresponding ground truth image. (Middle row) Displays the denoised outputs generated by the baselines and the STED-FM model. (Bottom row) Shows the residual image, calculated by subtracting the denoised output from the ground truth. The residual map highlights errors: magenta indicates regions where the denoised signal is higher, while green indicates regions where the signal is lower. b) Evaluation of the denoising performance across all models using pixel-wise metrics: MAE (mean absolute error), MSE (mean squared error), MSSSIM (multiscale structural similarity), and PSNR (peak signal to noise ratio). Higher MSSSIM and PSNR indicate better performance, while lower MAE and MSE are favorable. Boxplots present the the sample median, the first and third quantiles. The whiskers are 1.5 times the interquartile range. (N=187) c) Statistical comparison of model performance. The matrix summarizes the results of the resampling test comparing the performance of each baseline (row) against every other baseline (column) for the metrics presented in (b). The matrix includes the raw $p$-values for the pair-wise comparisons, allowing for the assessment of the significance of performance differences.

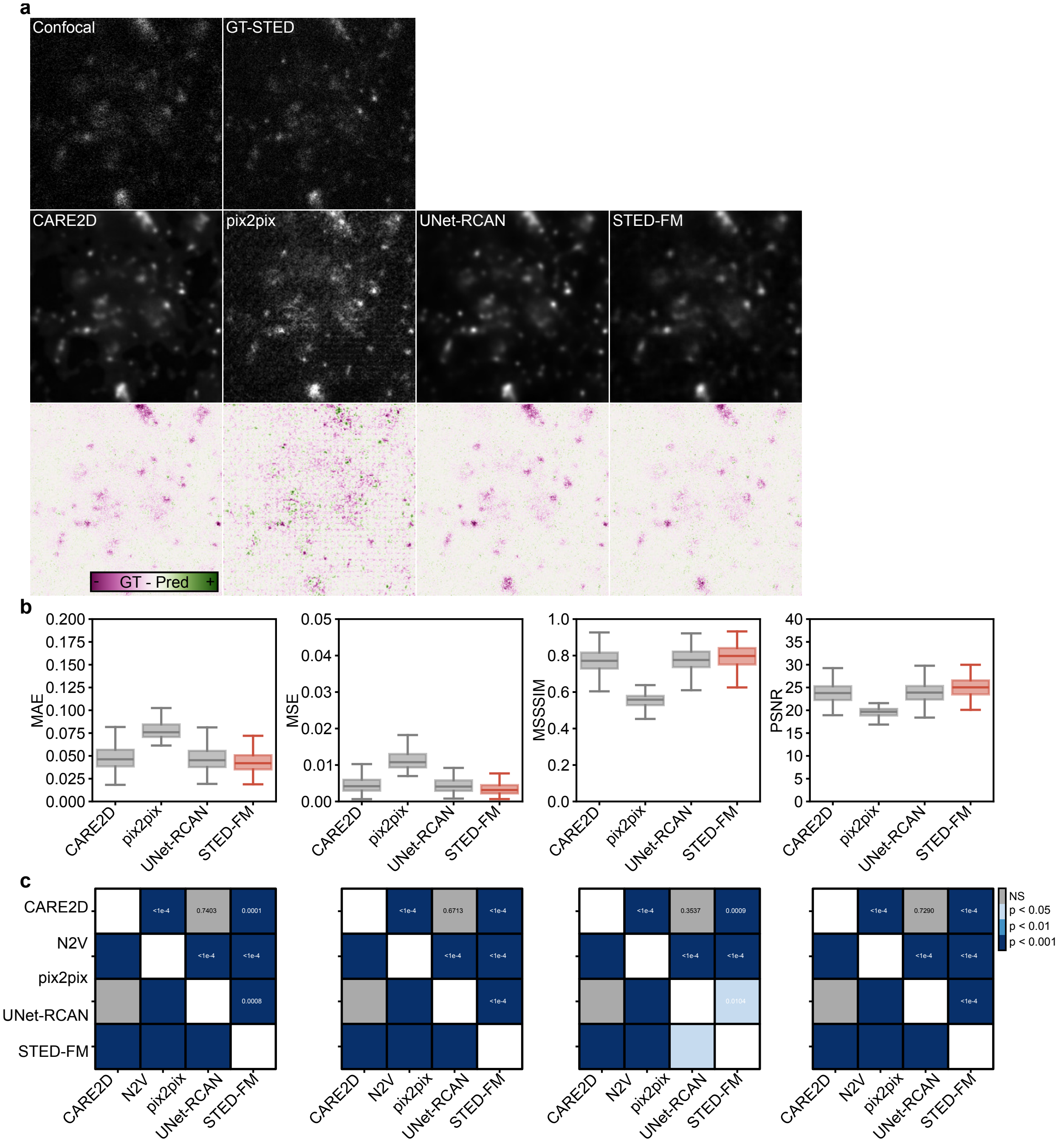


Supplementary Fig. 21: Super-resolution performance on the VGAT-SR (OOD-STED) dataset. a) Qualitative comparison and residual analysis. (Top row) Presents the confocal input image and the corresponding ground truth STED image. (Middle row) Displays the super-resolution outputs generated by the baselines and the STED-FM model. (Bottom row) Shows the residual image, calculated by subtracting the super-resolved output from the ground truth STED. The residual map highlights errors: magenta indicates regions where the super-resolved signal is higher, while green indicates regions where the signal is lower. b) Evaluation of the super-resolution performance across all models using pixel-wise metrics: MAE (mean absolute error), MSE (mean squared error), MSSSIM (multiscale structural similarity), and PSNR (peak signal to noise ratio). Higher MSSSIM and PSNR indicate better performance, while lower MAE and MSE are favorable. Boxplots present the the sample median, the first and third quantiles. The whiskers are 1.5 times the interquartile range. (N=187) c) Statistical comparison of model performance. The matrix summarizes the results of the resampling test comparing the performance of each baseline (row) against every other baseline (column) for the metrics presented in (b). The matrix includes the raw $p$-values for the pair-wise comparisons, allowing for the assessment of the significance of performance differences.

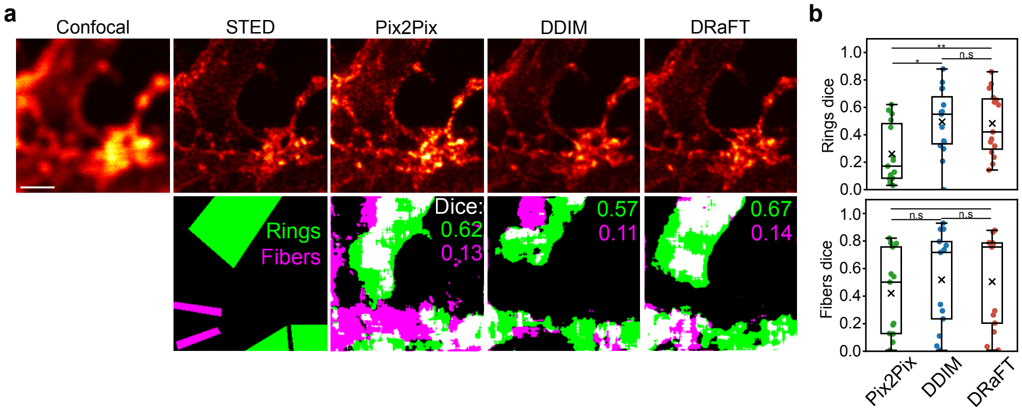


Supplementary Fig. 22: Algorithmic super-resolution performance of F-actin nanostructures from reference confocal images (N=26). a) Example images of generated F-actin nanostructures with the pix2pix, DDIM and DRaFT models from a reference confocal image and compared with the ground truth STED image. Segmentation masks of F-actin rings and fibers are shown for each examples. The ground truth masks correspond to the annotations from [14]. Segmentation of rings and fibers were obtained by using a pre-trained U-Net (Methods). b) Evaluation of the fidelity of reconstructed F-actin structures. Fidelity was quantified using the Dice score after segmenting the F-actin rings and fibers structures with a U-Net model pretrained on real STED images. From pixel-wise and segmentation metrics, DDIM and DRaFT are similar. Boxplots present the the sample median, the first and third quantiles, and 'x' the mean. The whiskers are 1.5 times the interquartile range. The p-values for the segmentation metrics are given in Supplementary Table 10.

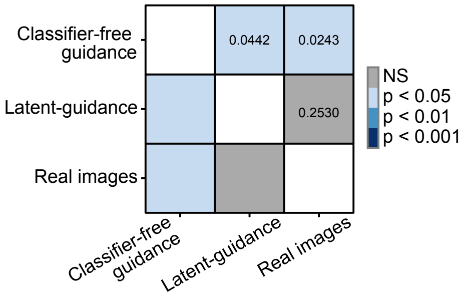


Supplementary Fig. 23: Results from the resampling statistics for the experiment from Fig. 4b. The F-statistic is significantly different ($p$-value=0.0079). A post-hoc resampling test is done to compare distributions in a one-to-one manner. N=4 experts.

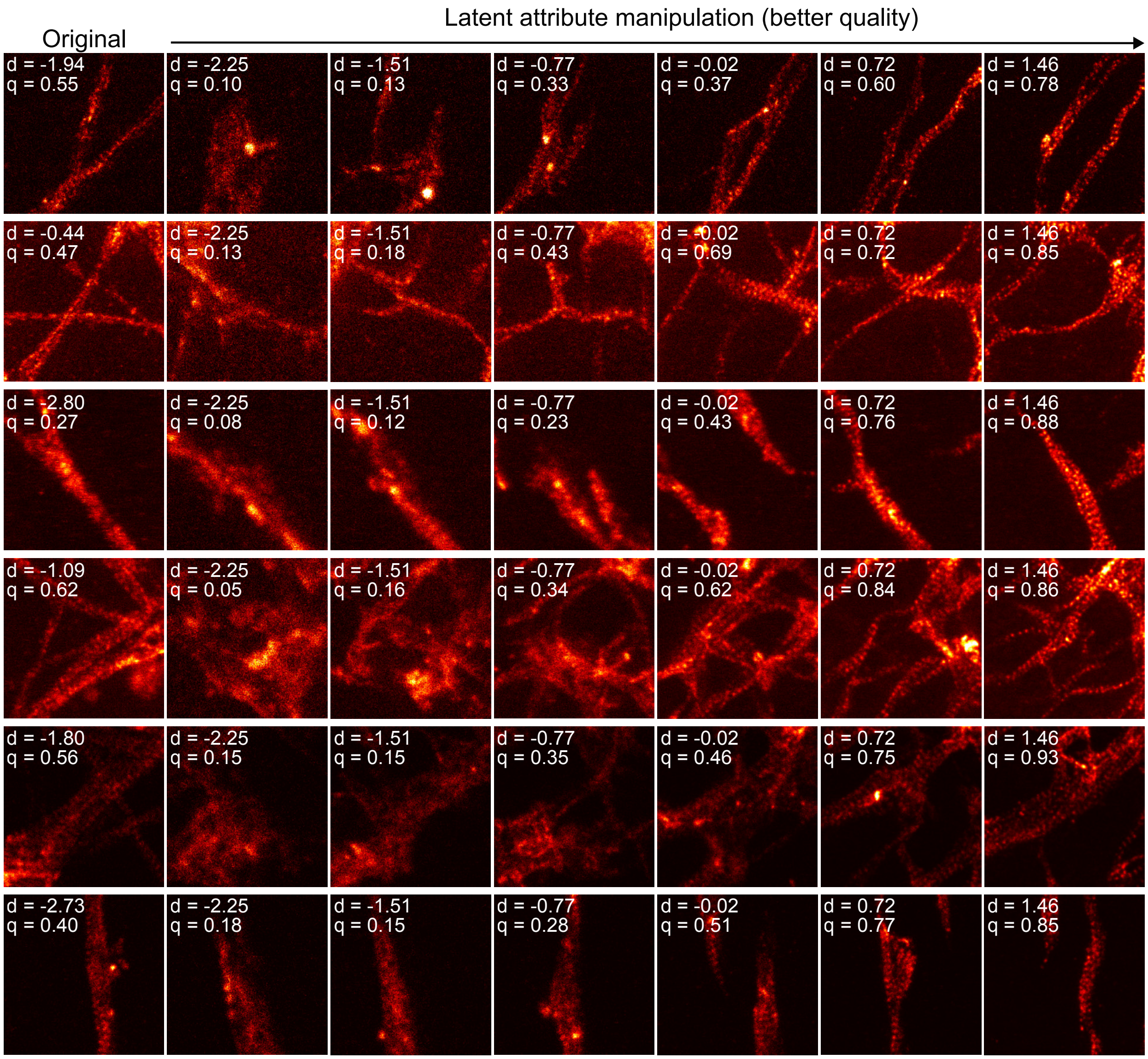


Supplementary Fig. 24: Example trajectories from the latent quality manipulation experiment. The quality score measured with a small convolutional neural network (Methods) is shown for each image. All images are 4.48 µm×4.48 µm

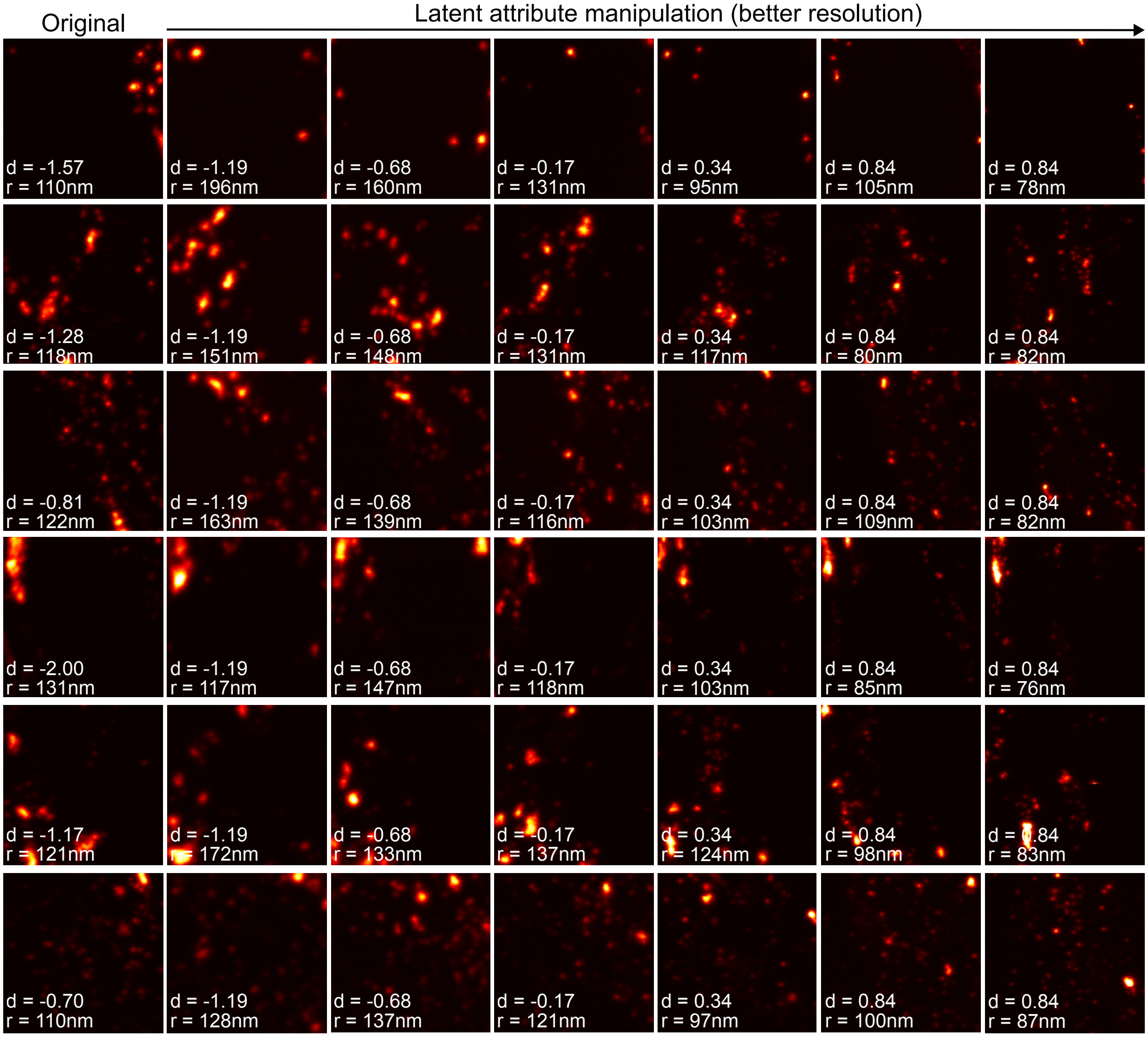


Supplementary Fig. 25: Example trajectories from the latent resolution manipulation experiment. Resolution was measured using the decorrelation analysis [20]. All images are 4.48 µm × 4.48 µm

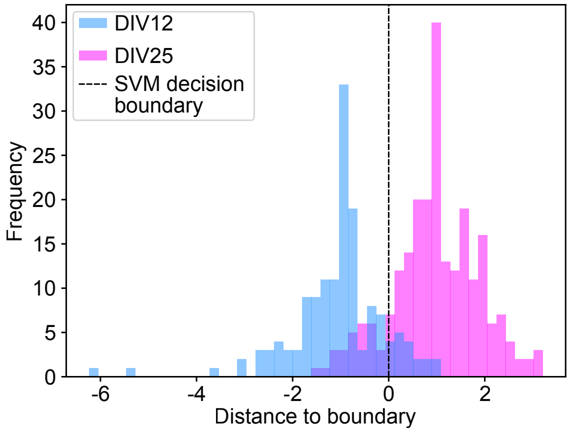


Supplementary Fig. 26: Histogram of DIV12 and DIV25 image embeddings' distance to the SVM decision boundary.

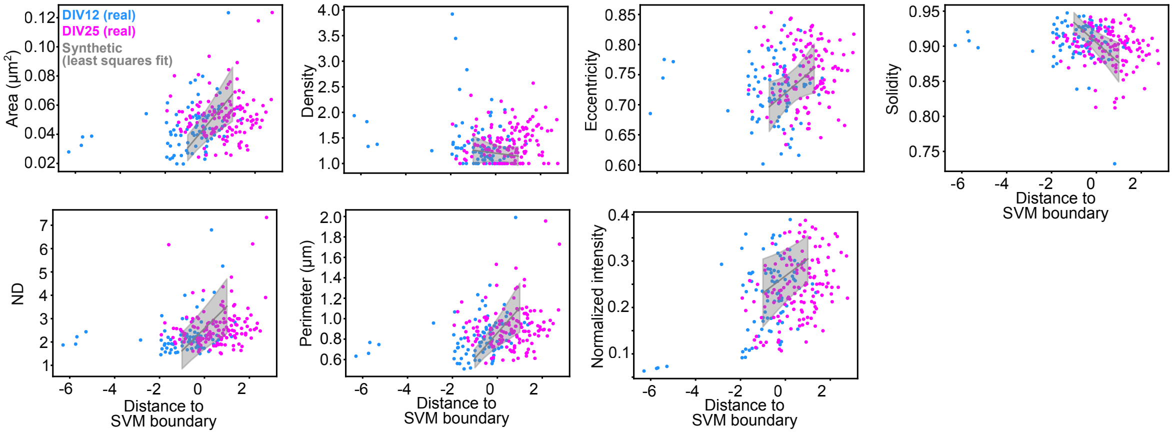


Supplementary Fig. 27: Raw image feature values as a function of distance to the SVM boundary for real DIV12 and DIV25 images. In gray is the average curve and standard deviation obtained from doing a least squares fit on the data points of the synthetic image trajectories generated by the conditioned diffusion model.

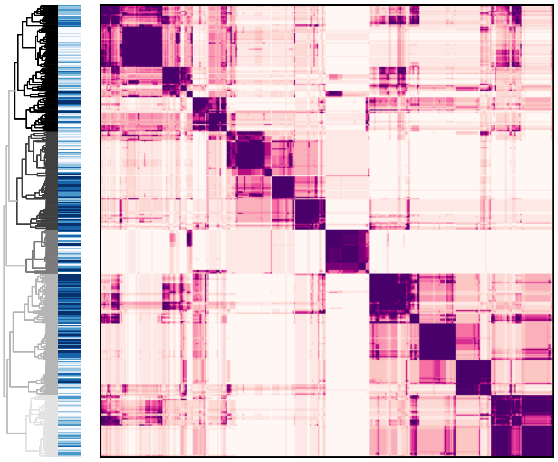

Supplementary Fig. 28: Hierarchical clustering on the consensus matrix from the *Synaptic Development* latent attribute manipulation experiment (Fig. 5a-e). *Dendrogram*: The dendrogram is color-coded by clusters. For each item in the dendrogram we report the distance to boundary in the blue colormap. The color gradient (from light to dark) represent the transition from DIV12 to DIV25. *Heatmap*: Darker colors are associated with images often clustering together.

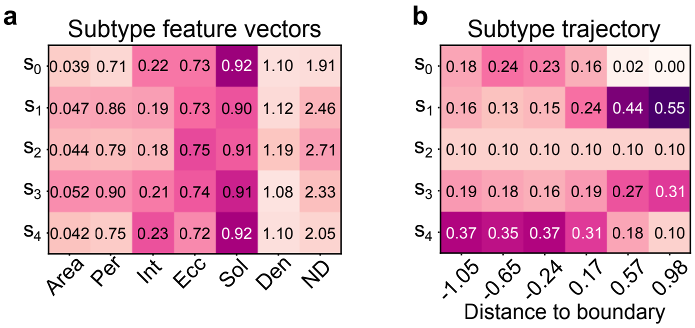


Supplementary Fig. 29: **a)** Average hand-crafted feature vectors of the different PSD95 subtypes. **b)** Proportion of PSD95 subtypes in the different trajectory time steps showing continuous transitions in subtype proportions.

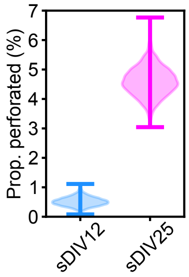


Supplementary Fig. 30: The proportion of perforated synapses is greater in synthetic images of DIV25 neurons compared to DIV12 neurons.

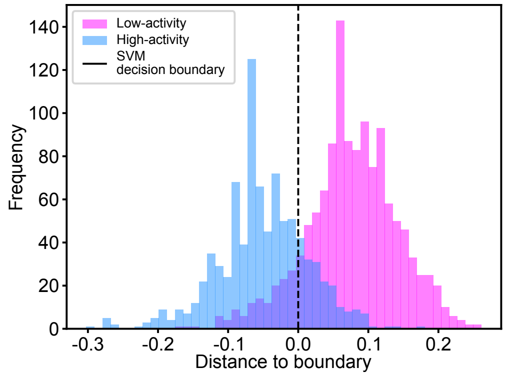


Supplementary Fig. 31: Histogram of Low- and High-activity image embeddings' distance to the SVM decision boundary of the *Activity-dependent F-actin dataset*.

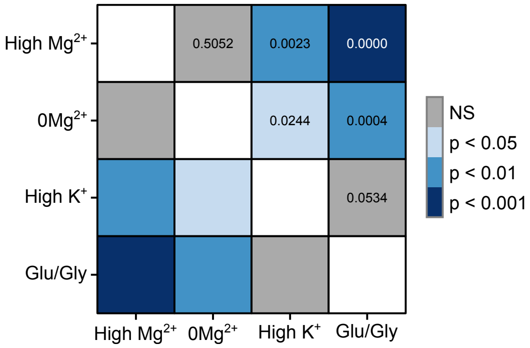


Supplementary Fig. 32: Results from the resampling statistics for the experiment from Fig. 5i. The F-statistic is significantly different ($p$-value=0.0002). A post-hoc resampling test is done to compare distributions in a one-to-one manner. High $Mg^{2+}$/Low $Ca^{2+}$ (N=86); $0Mg^{2+}$/Gly/Bic (N=70); High $^{+}$ (N=83); Glu/Gly (N=48)

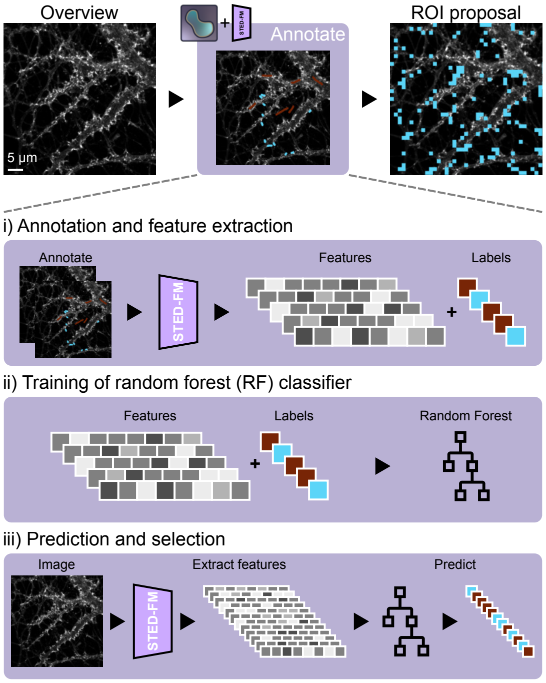


Supplementary Fig. 33: Overview of the automated microscopy approach. A three-step process is used for identifying the ROIs for subsequent imaging. i) Annotation and feature extraction: The overview image is imported into a napari plugin, allowing a user to manually annotate training patches. The latent features (embeddings) corresponding to each annotated patches are then extracted using the pre-trained STED-FM model. ii) Model training: These extracted features, along with their corresponding user-defined labels (positive/negative), form the training dataset for a random forest (RF) classifier. iii) Prediction and selection: STED-FM features are extracted for all patches selected by the classifier in the full field of view. The RF model is used to detect ROIs for the automated imaging step.

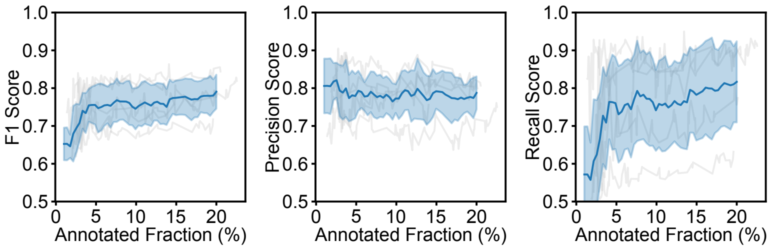


Supplementary Fig. 34: An evaluation strategy that simulated a human annotator was designed, allowing us to benchmark the performance of the random forest on a set of unseen images using a leave-on-out cross-validation scheme across five images. The detection metrics (F1-score, Precision, and Recall) are reported as a function of percentage of annotated patches in an image. The line and shaded area correspond to the average and standard deviation obtained over five repetitions.

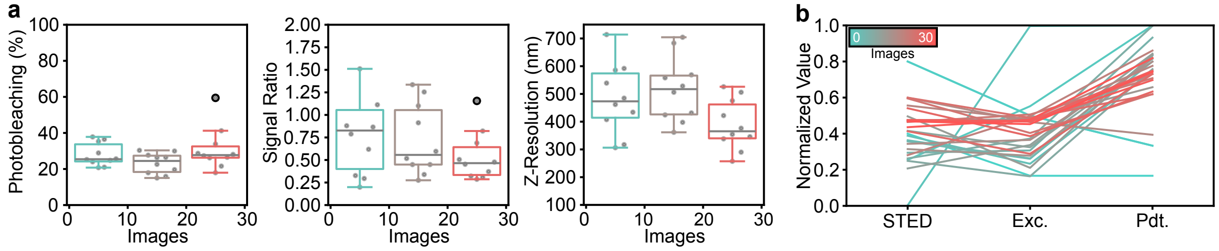


Supplementary Fig. 35: Bandit optimization. a) Evolution of the imaging optimization objectives (Photobleaching, Signal Ratio, and Z-Resolution; Methods). Boxplots in bins of 10 images. b) Evolution of the imaging parameters (STED power, Excitation power, and pixel dwell time). Supplementary Tab. 16 presents the non-normalized imaging parameters. Boxplots present the the sample median, the first and third quantiles. The whiskers are 1.5 times the interquartile range.

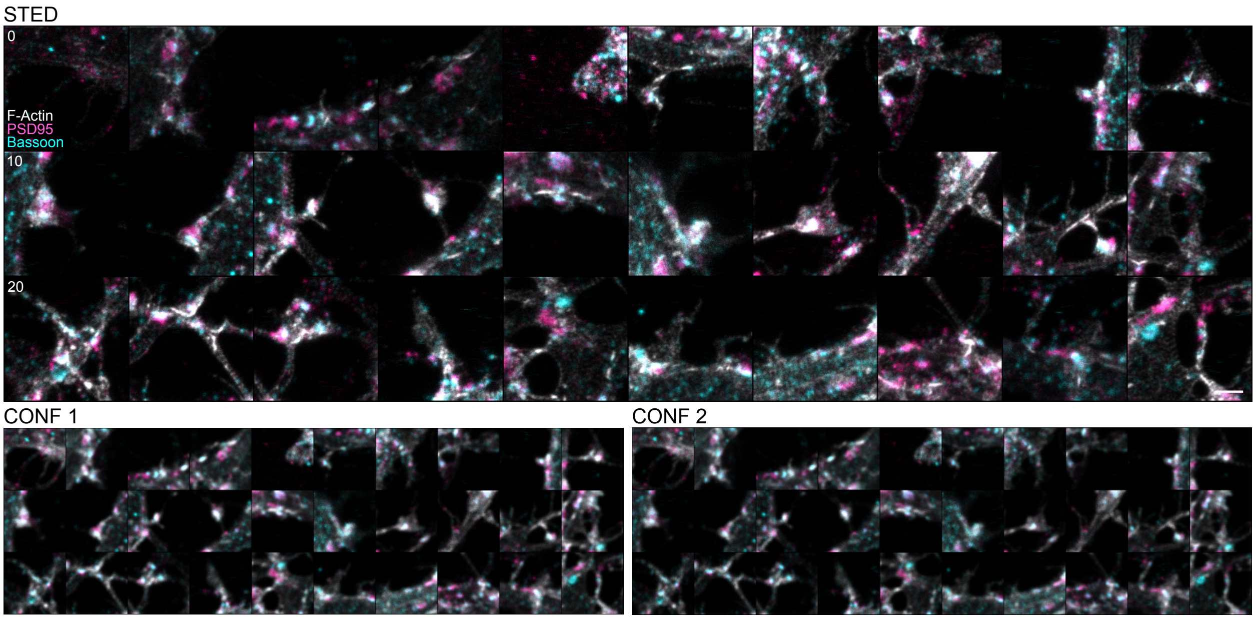


Supplementary Fig. 36: All 3-color images acquired with the RL models, one model per protein (F-actin STAR 635; PSD95 Alexa 594; Bassoon ATTO 490LS). A confocal image is acquired before (CONF1) and after the STED image (CONF2) to measure photobleaching. The displayed signal of the STED and CONF 2 images are normalized to their corresponding CONF 1 image. Scale bar: 1 µm.

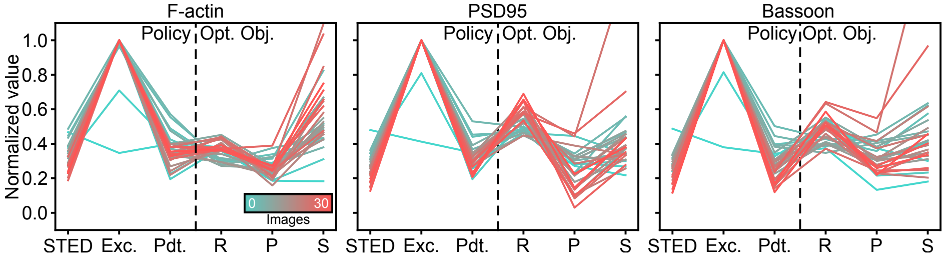


Supplementary Fig. 37: Evolution of the imaging parameter selection (STED and Excitation power and pixel dwell time) and imaging optimization objectives (R: Resolution, P: Photobleaching, and S: Signal ratio) for the 3-color STED imaging. From left to right: F-actin, PSD95, and Bassoon.

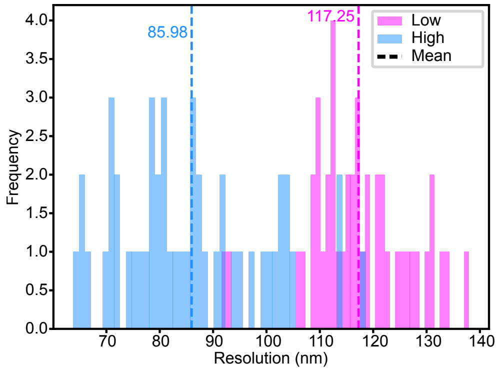


Supplementary Fig. 38: Histogram of image resolutions in the *Image resolution dataset*.
